# Universal inverse modelling of point spread functions for SMLM localization and microscope characterization

**DOI:** 10.1101/2023.10.26.564064

**Authors:** Sheng Liu, Jianwei Chen, Jonas Hellgoth, Lucas-Raphael Müller, Boris Ferdman, Christian Karras, Dafei Xiao, Keith A. Lidke, Rainer Heintzmann, Yoav Shechtman, Yiming Li, Jonas Ries

**Affiliations:** Department of Physics and Astronomy, University of New Mexico, Albuquerque, NM, USA; Department of Biomedical Engineering, Guangdong Provincial Key Laboratory of Advanced Biomaterials, Southern University of Science and Technology, Shenzhen, China; European Molecular Biology Laboratory, Cell Biology and Biophysics, Heidelberg, Germany; Department of Biomedical Engineering, Technion–Israel Institute of Technology, Haifa, Israel; Institute of Physical Chemistry and Abbe Center of Photonics, Friedrich-Schiller-University Jena, Jena, Germany; Leibniz Institute of Photonic Technology, Albert-Einstein-Straße 9, 07745, Jena, Germany; Max Perutz Labs, Vienna Biocenter Campus (VBC), Dr.-Bohr-Gasse 9, 1030, Vienna, Austria; University of Vienna, Center for Molecular Biology, Department of Structural and Computational Biology, Dr.- Bohr-Gasse 9, 1030, Vienna, Austria; University of Vienna, Faculty of Physics, Boltzmanngasse 5, 1090 Vienna, Austria; Department of Mechanical Engineering, The University of Texas at Austin, Austin, TX, USA; Collaboration for joint PhD degree between Southern University of Science and Technology and Harbin Institute of Technology, Harbin, 150001, China; Currently at JENOPTIK Optical Systems GmbH, Jena, Germany

## Abstract

The point spread function (PSF) of a microscope describes the image of a point emitter. Knowing the accurate PSF model is essential for various imaging tasks, including single molecule localization, aberration correction and deconvolution. Here we present uiPSF (universal inverse modelling of Point Spread Functions), a toolbox to infer accurate PSF models from microscopy data, using either image stacks of fluorescent beads or directly images of blinking fluorophores, the raw data in single molecule localization microscopy (SMLM). The resulting PSF model enables accurate 3D super-resolution imaging using SMLM. Additionally, uiPSF can be used to characterize and optimize a microscope system by quantifying the aberrations, including field-dependent aberrations, and resolutions. Our modular framework is applicable to a variety of microscope modalities and the PSF model incorporates system or sample specific characteristics, e.g., the bead size, depth dependent aberrations and transformations among channels. We demonstrate its application in single or multiple channels or large field-of-view SMLM systems, 4Pi-SMLM, and lattice light-sheet microscopes using either bead data or single molecule blinking data.

## Introduction

A microscope image is formed by the convolution of the fluorescent light emitted by the sample structure with the point spread function (PSF). Precise knowledge of the PSF is of fundamental importance for many applications: quantification and optimization of microscope performance^1–3^, deconvolution of microscopy images to increase contrast and resolution^4,5^, estimation and correction of system and sample induced aberrations^6^, and evaluation of single molecule properties (e.g., 3D localization, dipole orientation and color)^7–10^.

PSF modelling is especially important for single-molecule localization microscopy (SMLM), a super-resolution method that relies on sparse activation of switchable fluorophores over many camera frames, followed by precise localization of the emitter positions^11^. In 3D SMLM, the z coordinate of the emitter can be inferred from the shape of the PSF after introducing suitable aberrations (PSF engineering)^12–14^, when measured at two or more different focal planes (bi-plane^15,16^, LLS-PAINT^17^) or from relative intensities in 3 or 4 channels after interference of the fluorescence detected with two opposing objectives (iPALM or 4Pi-SMS)^18,19^. All these approaches rely on knowing the precise shape of the PSF, as any inaccuracy of the PSF model (‘model mismatch’) will result in a z-dependent bias in the position estimation and reduced accuracy.

A microscope PSF can be calculated in either the spatial domain or the Fourier domain. In spatial domain modelling, PSFs are described by their intensity values on a 3D grid. The Gaussian PSF model is extensively used in 3D SMLM because of its ease of calibration and fitting. However, it can result in large localization bias (Fig.1)^12,20^. Realistic PSF models, built directly from a z stack of images of fluorescent beads immobilized on a coverslip^21^ or after spline interpolation^22–25^, have greatly improved the accuracy of 3D SMLM. However, spatial-domain based PSF models usually cannot describe variations of experimental conditions (e.g., depth or sample induced aberrations) and are difficult to calibrate for complex microscope modalities (e.g., 4Pi-SMLM).

**Figure 1.**
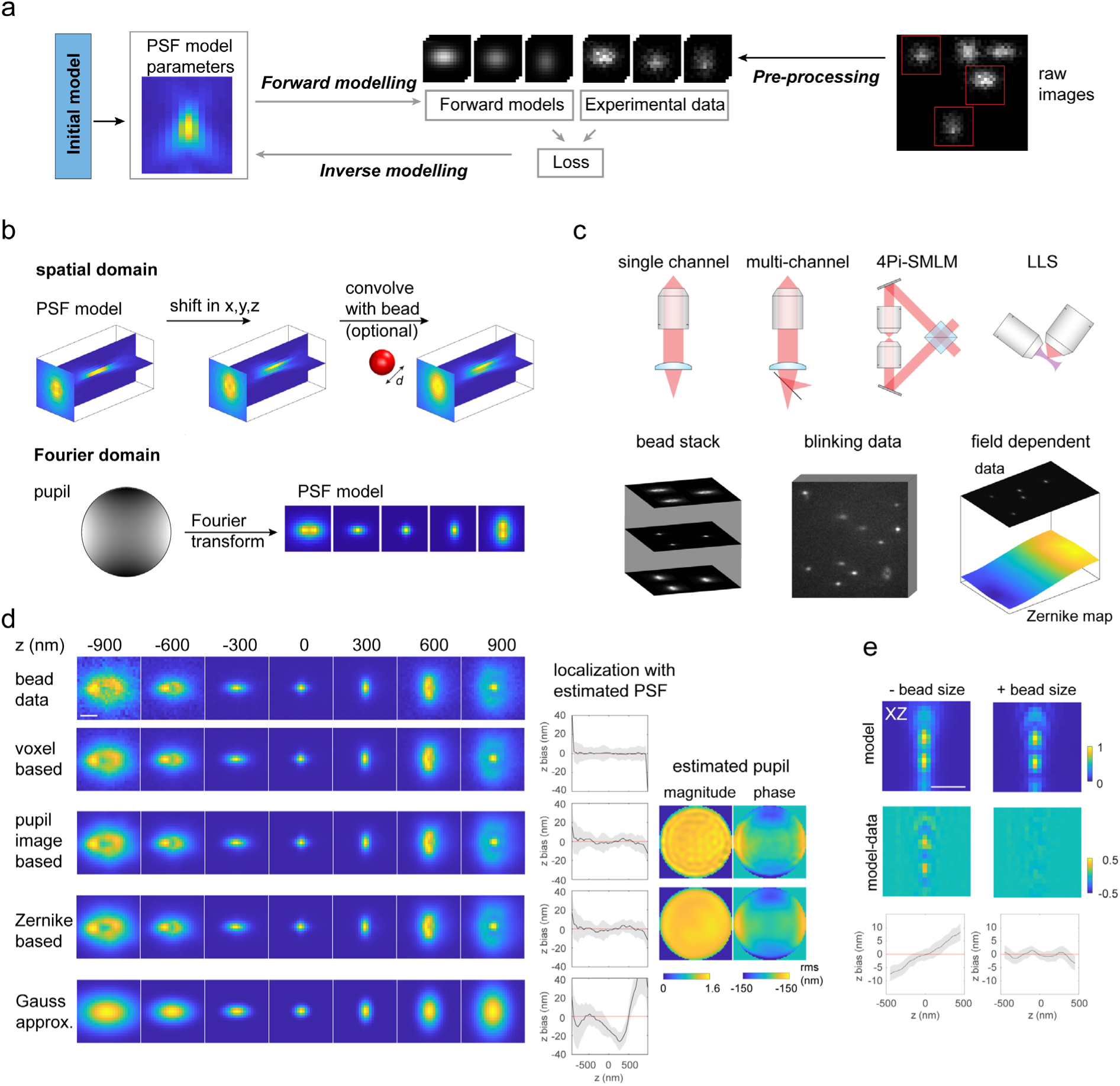
Concept and modalities of uiPSF. (a) Inverse modelling concept of uiPSF. (b) PSF modelling methods in both the spatial and the Fourier domain. (c) Supported imaging and data modalities. (d) Example PSF modelling results from uiPSF and Gaussian approximation for a single-channel astigmatic imaging system. (e) Effect of bead size on modelling 4Pi-SMLM PSFs using the voxel-based PSF model. Scale bar, 1 µm (d,e).

Alternatively, instead of expressing PSFs in terms of a 3D volume in the spatial domain, they can be fully parametrized in the Fourier domain by a complex-valued 2D image, the pupil function. The spatial domain 3D PSF can then be calculated using the scalar or vectorial diffraction theory^26–28^. Different experimental conditions (e.g., aberrations, depths, wavelength) can be easily incorporated. Decomposing the phase of the pupil function into the orthogonal basis of Zernike polynomials reduces the number of parameters^29^ and results in intuitive aberration modes^30^. Pupil functions can be determined by phase retrieval using a modified Gerchberg-Saxton (GS) algorithm^31–33^ or by direct fitting to Zernike coefficients^28,34,35^ or a 2D pupil image^36,37^, or by deep learning^38–40^. Lately, Xu *et. al.* applied a GS algorithm to retrieve an *in situ* PSF model from single blinking fluorophores by averaging single fluorophore images at similar z positions to construct a 3D PSF image stack^41^. We recently determined the spatially variant PSFs across a large field of view (FOV) by retrieving changes of Zernike coefficients^42^.

Accurate PSF modelling often requires many parameters to model the complex image formation process. However, it is often too complicated to incorporate the complete experimental conditions, e.g., the finite size of beads, field dependent aberrations, and sample induced aberrations. Therefore, current approaches are often limited to simple imaging modalities, and lack accuracy in multi-channel applications (multi-color, bi-plane or 4Pi-SMLM) that require not only knowledge of the precise PSF model in each channel, but also an accurate transformation among the channels. Additionally, existing methods usually work with experimental data of small size (e.g. 1-3 bead stacks) and simplified experimental conditions, and cannot extract the abundant information embedded in the massive microscopy data. Thus, a universal, robust and easy-to-use approach that can infer accurate PSF models from relatively large experimental data for various microscope modalities is still missing.

To accurately retrieve all parameters of a PSF model, a differentiable imaging model is required. However, such a model including derivatives is complex and lengthy in its explicit form. Thus, we took advantage of the automatic differentiation functionality of TensorFlow and developed a versatile and modular toolbox, termed uiPSF (universal inverse modelling of Point Spread Functions), that uses inverse modelling to extract accurate PSF models for most SMLM imaging modalities from bead stacks or *in situ* from blinking emitters. Supported modalities include astigmatic, Tetrapod, double-helix, 4Pi and multi-channel PSFs, both in the spatial domain and in the Fourier domain (Fig. 1a-c). We account for realistic experimental conditions, such as bead size, intensity fluctuations, pixelation effects, sample drift and field-dependent aberrations.

## Results

### Inverse modelling of PSFs

Determining a microscope PSF from measurements is an inverse problem, which requires building a continuous model function through noisy and pixelated image observations under certain imaging conditions. As the image formation process in a microscope is well studied, it is relatively easy to model microscopy images given the PSF and the positions of the emitters (‘forward modelling’). The inverse problem – determining a parameterized PSF model from experimental data – is much more difficult, as the forward model cannot be inverted easily. This inverse problem can be solved by an iterative approach called inverse modelling, in which the forward models are built from initial parameters and are compared to the experimental data by means of a value (‘loss function’) measuring the current quality of agreement, and an optimization algorithm adjusts the model parameters to decrease the loss (Fig.1a). This process is repeated until convergence. Efficient optimization requires the derivatives of the forward model with respect to the model parameters. We thus implemented inverse modelling of PSFs using the TensorFlow package, where the differentiation is automatically performed, allowing us to use increasingly complex forward models without explicitly calculating derivatives.

### PSF modelling in the spatial domain

We now describe a simple implementation in the spatial domain, in which the PSF, parametrized by a 3D image stack (‘voxels’), is determined from bead data: In a pre-processing step, the beads are identified and regions of interest (ROIs) around the beads are cropped from the raw camera frames (SI Note 1). We initialize the PSF model as a 3D array with all values set to a small value close to zero and set the initial bead positions to zero (Fig. 1a and SI Note 2). The forward model then consists of shifting the 3D array to the estimated bead positions using sub-pixel interpolations, which are achieved by applying phase-ramps in Fourier space (Fig. 1b). We used the mean square error (MSE) between the forward models and the data as the loss function. Regularization terms are added to the loss function to reduce overfitting and edge effects, and to ensure that the PSF model and all photon counts remain positive (SI Note 2). A suitable optimizer (L-BFGS)^43^ then minimizes the loss to optimize the voxel values in the PSF model and the positions, photons and backgrounds of the beads (SI Table 1).

### PSF modelling in the Fourier domain

The parametrization of the PSF as a 3D array of intensity values has the advantage that it directly measures the impulse function of a microscope in the spatial domain where it forms the microscope image. No prior knowledge of the image formation process is needed. However, this comes at the expense of typically tens of thousands of fitting parameters and the difficulty to incorporate specific imaging conditions (e.g., depth dependent aberrations) into the model. An alternative representation of the PSF is the pupil function *h*(*k_x_, k_y_*), which describes the electric field at the pupil plane. Scalar or vectorial diffraction theory can then be used in the forward model to calculate the PSF, *U*_PSF_(*x* − *x_i_*, *y* − *y*_*i*_, *z*_*i*_), of a emitter at location (*x_i_*, *y_i_*, *z_i_*) from the pupil function using Fourier transforms and knowledge about the microscope such as the numerical aperture (NA) of the objective, the refractive index of the immersion oil *n* and the emission wavelength *λ* (Fig. 1b). The scalar PSF model is given by (see SI Note 3 for vectorial PSF model),

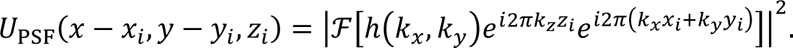

Here ℱ denotes a 2D Fourier transform and *k*_*x*_, *k*_*y*_, *k*_*z*_ are the Cartesian components of the wave vector ***k***, where its magnitude *k* = *n*/*λ*. The term *e*^*i*2*π*_*z*_*z*_*i*_^ accounts for the defocus phase and 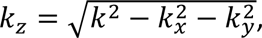 and the term *e*^*i*2*π*(*k*_*x*_*x*_*i*_+*k*_*y*_*y*_*i*_)^ accounts for the shift phase to location (*x*_*i*_, *y*_*i*_). The pupil function of most imaging system has a circular boundary, and its cutoff frequency is defined by *NA*/*λ*.

The complex pupil function is typically expressed in terms of its magnitude *A*(*k*_*x*_, *k*_*y*_) and its phase Φ(*k*_*x*_, *k*_*y*_):

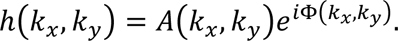

*A* and Φ as 2D arrays can be directly estimated, we term this modelling method as pupil-image based modelling. To reduce the number of parameters even further and to increase the robustness in the presence of noise, we can express *A* and Φ in the basis of Zernike polynomials *Z*_*p*_,

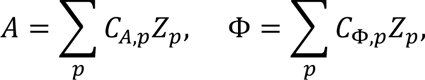

parametrizing the PSF in terms of the Zernike coefficients *C*_*p*_. We term this modelling method as Zernike-based modelling. Both pupil-image based and Zernike-based modelling are Fourier-domain or pupil-based modelling methods. We found that including up to 8th radial order (Noll index) is sufficient, resulting in 45 Zernike coefficients for each expansion. We can further decrease the number of parameters by assuming *A* to be rotationally symmetric or constant, with a small sacrifice in the axial bias, but it sometimes led to larger lateral biases (SI Figs. 1-2).

For Fourier-domain based modelling, we use a maximum-likelihood based loss-function to account for the shot noise and to improve the precision especially for low signal-to-noise conditions (SI Note 3).

### Validation

We validated our framework extensively with simulations, in which we can directly compare the results to the ground truth (SI Figs. 3-7). We found that the estimated PSF model can achieve a localization precision at the information limit (the Cramér–Rao lower bound (CRLB)) (SI Figs. 3-4).

We then performed an extensive experimental validation of uiPSF: (a) we compared the estimated PSF model with the raw data visually and by calculating residuals (SI Figs. 8-9). (b) We fitted the bead stacks with the PSF model to retrieve emitter’s *x*, *y* and *z* positions in each frame. A frame-dependent bias in any direction denotes a model-mismatch (Fig. 1d and SI Fig. 1-2, 8-10). The localization test shows that the estimated PSF model can achieve a localization precision close to the information limit (SI Fig. 8-9). (c) We used nuclear pore complexes (NPC) as 3D reference standards^44^. As the labeled protein Nup96 forms two rings with a separation of ∼50 nm, we can investigate distortions in *z* by evaluating how this separation changes with the axial position of the nuclear pores (ED Fig. 1).

Compared to previous approach that aligned and averaged multiple beads using cross-correlation^23^, the voxel-based PSF model from uiPSF show less bias for both simulated and experimental data (SI Fig. 11). Compared to Zola 3D^34^, a previous approach that estimates Zernike polynomials from bead data, uiPSF, extracting Zernike coefficients from multiple beads (typically >20), shows a smaller bias by incorporating challenging experimental conditions in a vectorial PSF model (SI Fig. 12).

### Extensions of the forward model

The forward models introduced above can be made more realistic by including additional experimental conditions.

#### Bead size

Large fluorescent beads (100 nm or 200 nm) are substantially brighter than smaller ones. They contain more fluorophores and thus show less intensity fluctuations and photobleaching. As the bead images can be described as a convolution of the PSF with the bead shape, they are more blurred than the PSF itself. We can include this blurring in the forward model *U*_*i*_ by a convolution with a uniform fluorophore distribution in the bead *g*^28^:

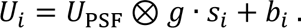

Here, *s*_*i*_ is the total photon count of bead *i* and *b*_*i*_ is the average background photon count per pixel in the ROI. For fitting of single molecules, *U*_PSF_ is then directly used without the bead convolution. Indeed, when we calibrated a PSF with 200 nm beads and validating the calibration on 40 nm beads mimicking single fluorophores, we found that considering the bead size substantially improves the accuracy of the PSF model and reduces the bias in z-localization for both astigmatic PSFs (SI Fig. 8) and 4Pi PSFs (Fig. 1e and SI Fig. 9, SI Note 2).

#### Apodization

Most objectives are designed to be aplanatic, where at the nominal focal plane the image is free of coma and spherical aberrations. The imaging system satisfies the Abbe Sine condition. In this case, we assume that the objective can convert a spherical wavefront coming from an emitter at the nominal focal plane to a plane wavefront at the back focal plane (BFP) of the objective. To satisfy energy conservation, an apodization term, describing the amount of the incident area at the spherical surface projected to an unit area of the BFP, is incorporated into the transmission of the rays^45^ (SI Note 3). This apodization term also assumes that the refractive index is matched between the sample medium and the immersion medium. In the case of refractive index mismatch, the apodization term is modified so that the retrieved pupil magnitude is nearly uniform (SI Fig. 13).

#### Pixilation

The finite size of the pixels results in a difference between the PSF value at the pixel center compared to the intensity averaged over the pixel. Neglecting this effect can lead to localization errors^46^. We can decrease the impact of pixilation by oversampling the PSF followed by binning, which we implemented into uiPSF. We found that pixilated PSFs are similar to slightly blurred PSFs, allowing us to replace the computationally expensive oversampling by applying an extra Gaussian blur (ED Fig. 2 and SI Note 3).

#### Intensity fluctuations, drift, and shear

Beads can flicker intrinsically, especially when imaged under high intensities, or when using multi-mode excitation that produces speckles^47^. Thus, we have the option to treat the intensity of each bead in each frame as a free fitting parameter (SI Fig.6). For pupil-based modelling, a per-frame intensity is always assumed unless otherwise set. Similarly, we can allow for lateral drifts (frame-wise shifts) in our forward model. We found that by estimating the lateral drift, the resulting PSF model will be less affected by lateral drift from the bead data (SI Fig. 14). Additionally, some microscopes, like the lattice light-sheet microscope (Fig. 1c)^48^ or single-objective light-sheet microscopes utilizing remote-focusing^49,50^, translate the sample not along the optical axis, leading to sheared bead stacks. We can directly take this shearing into account in our forward model, avoiding computationally expensive and imprecise interpolations (ED Fig. 3).

#### Extra blurring

After considering the imperfections described above, we found that even the full vectorial forward model generally results in slightly sharper images than the experimental data, as has been observed previously^31,32,34,51^. The reasons for this are not entirely clear but could stem from non-isotropic dipole radiation, or system vibrations. By adding a convolution with a 2D Gaussian kernel with user-defined or estimated standard deviations in both x and y dimensions (SI Note 3), we can successfully describe this extra blurring, leading to more accurate PSFs and a reduced bias (ED Fig. 2).

#### Refractive index mismatch and supercritical angle emission

When using an oil objective, the refractive index mismatch results in depth-dependent aberrations^6^. Additionally, for high-NA objectives, emitters close to the coverslip can emit into super-critical angles (an angle in the immersion medium which is beyond the critical angle), leading to PSFs that are different from those of emitters a few hundred nanometers above the coverslip. When using a forward model in the Fourier domain (pupil-image based and Zernike-based models), the supercritical-angle fluorescence and depth-induced aberrations are incorporated and a theoretical PSF model can be calculated for arbitrary imaging depths (ED Fig. 4).

### Multi-channel PSFs

Two or more detection channels are used in SMLM for multi-color imaging^52–55^, for bi-plane 3D localization^15,16^, for 3D dense emitter fitting^56^ or single-molecule orientation imaging^57^. Individual PSFs for each channel together with a transformation among the channels define a multi-channel PSF, which can improve the accuracy in global fitting^58^ compared to fitting channels individually. We extended our inverse modeling approach to multiple channels to determine a PSF for each channel together with an affine transformation matrix (Fig. 2a-b and SI Table 2) and validated the estimated PSF model both with bead data (SI Fig. 2) and in 3D dual-color imaging of NPCs (Fig. 2c-d).

**Figure 2.**
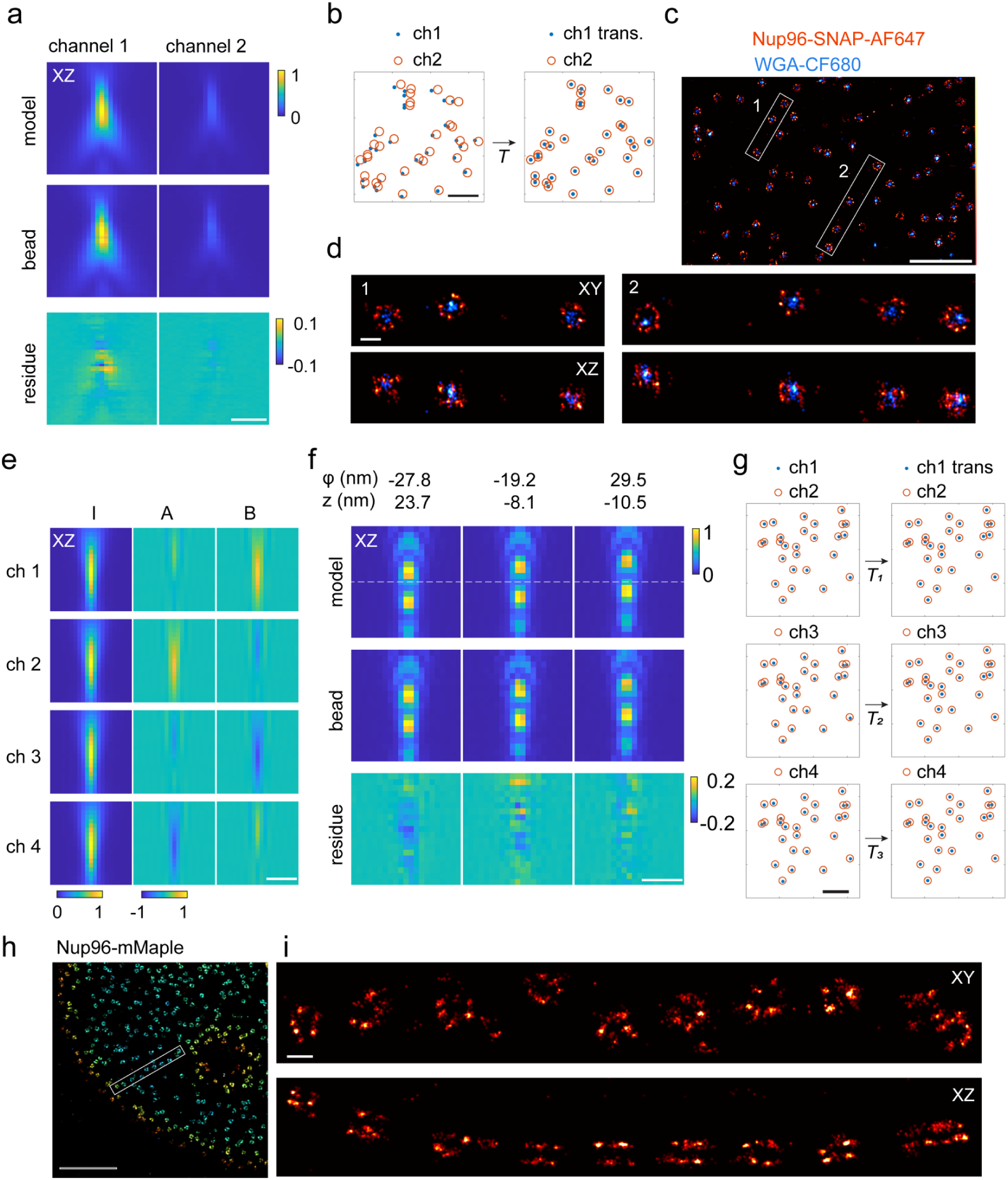
Application of uiPSF to multi-channel ratiometric dual-color SMLM and 4Pi-SMLM. PSF models are estimated from bead data. (a, b) Estimated PSF model and the transformation for a dual-color system. (c, d) 3D dual-color SMLM imaging of NPC, where Nup96-SNAP is labeled with Alexa Fluor 647 and WGA is labeled with CF680. (e) Estimated IAB model for each channel of a 4Pi-SMLM system. (f) Example beads and corresponding forward models at various z and *φ* positions. (g) Estimated transformation of each target channel to the reference channel of a 4Pi-SMLM system. (h,i) 4Pi-SMLM imaging of Nup96-mMaple. Scale bars, 5 µm (b, g), 2 µm (h), 1 µm (a, c, e, f), 100 nm (d, i).

Interferometric 4Pi-SMLM^18,19,59^ achieves excellent axial localization precisions by interfering the emission of fluorophores detected with two opposing objectives (Fig. 1c). Depending on the z position of the fluorophore, the fluorescence intensity is modulated by constructive or destructive interference and is detected in 3 or 4 channels at different interference phases. The PSF in each channel is described by an incoherent enveloping term plus a modulation term^19^. A small change in the emitter’s z position or interference phase *φ*_0_ will shift the modulation and affect the PSF shape. This prevents averaging of multiple beads and makes it difficult to recover the 4Pi-PSF model. Here we overcome this limitation by choosing a PSF representation (IAB-based 4Pi-PSF model) in which the interference phase term is decoupled from the emitter’s z position^25^. In this representation, the 4Pi PSFs in each channel are generated from a model that consists of three 3D matrices (*I*, *A*, and *B*, Fig. 2e) and a phase parameter *φ* = 2*πkz* + *φ*_0_, where the two coupling parameters, *z* and *φ*_0_, are absorbed into one parameter (SI Table 3),

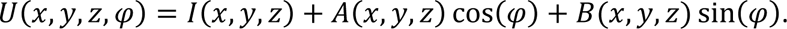

We treat *z* and *φ* as independent variables for each bead (Fig. 2f and SI Fig. 9). An affine transformation matrix for each channel is also estimated (Fig. 2g). This model is used for estimating a voxel-based representation of the 4Pi-PSF model. For inverse modelling in the Fourier domain, we coherently add PSF models from both objectives, each described by the pupil function images or Zernike coefficients (SI Fig. 10 and SI Note 3). The resulting 4Pi PSFs in combination with global fitting allowed us to reach high 3D resolution even on dim but live-cell compatible photoconvertible fluorescent proteins, as demonstrated on NPCs labeled with mMaple (Fig. 2h-i and SI Fig. 15).

### *In situ* PSF modelling

Although PSF models estimated from fluorescence bead data can achieve excellent localization precisions, they are still different from the emission patterns of a single fluorophore *in situ* in the cell. First, the bead size of 40-200 nm is large compared to the size of a fluorophore^60^, ∼2 nm. Second, bead data is usually collected with beads at the coverslip, while fluorophores in cells can be far above the coverslip. In this case, aberrations induced by a refractive index mismatch of the imaging buffer, or the sample, cannot be captured by the bead data. The resulting model mismatch can cause a bias in the fitted position and a loss in localization accuracy (ED Fig. 5).

Here we overcome this limitation by directly estimating an *in situ* PSF model from the blinking single fluorophores, the raw data in SMLM. Our *in situ* PSF modelling approach simultaneously determines the 3D positions of the emitters and either image-based or Zernike-based pupil functions. It consists of four major steps: 1) select candidate emitters from raw SMLM data; 2) localize emitters using an initial PSF model; 3) select several thousand of emitters spanning the entire z-range, 4) estimate the PSF model from the selected emitters. This estimated PSF model is then used as the initial PSF model and steps 2-4 are repeated. Usually, two iterations are sufficient for most tested data. For data with low signal-to-noise ratio (SNR) or complex emission patterns, 5-6 iterations are required to converge (SI Fig. 16).

We initially evaluated the *in situ* PSF modelling of uiPSF on simulated data (SI Fig. 17). Specifically, we generated a single-molecule dataset based on a vectorial PSF model (SI Note 4) with a set of Zernike aberrations. The estimated pupil and *x*, *y*, *z* positions demonstrated excellent agreement with the ground truth of the simulated data and the position estimates reached their theoretical precision limit, defined by the CRLB. We further examined how the number of single molecules and axial sampling methods affect the fitting precision. We found that the precision of Zernike coefficients improves with larger number of single molecules (SI Fig. 18) and can reach values below 1 nm (the root-mean-square (RMS) error of the wavefront) with 250 single molecules (5000 photons / localization, 10 background photons per pixel, z-range -600 nm to 600 nm). Moreover, selecting a larger fraction of defocused single molecules provided additional information, resulting in a higher precision of estimating the Zernike coefficients (SI Fig. 18).

We experimentally tested the *in situ* PSF estimation on beads embedded in agarose gel for which images were collected at stage positions from 0 to 25 µm. The per-frame z positions from each bead are treated as independent variables to mimic *in situ* single-molecule data and an *in situ* PSF model is estimated for each bead to verify uiPSF’s capability of modelling the emission patterns at different imaging depths. Compared with the PSF estimated from beads on the coverslip, the *in situ* PSF models can accurately model the depth of each bead and the fitted z positions using the *in situ* PSF model shows good linearity with the stage positions (ED Fig. 5).

We then validated our approach on single-molecule data taken from labeled microtubules. With a deformable mirror (DM) in the detection beam path, we introduced defined aberrations corresponding to individual Zernike modes. Comparing those DM-generated aberrations to the Zernike modes estimated from the *in situ* PSF modelling from the single-molecule data, we found an excellent correlation (Fig. 3a,b). Residual differences, apparent as non-zero off-diagonal terms, are likely due to remaining aberrations from the imaging system and the fact that the aberration modes generated by DM can only approximate pure Zernike modes.

**Figure 3.**
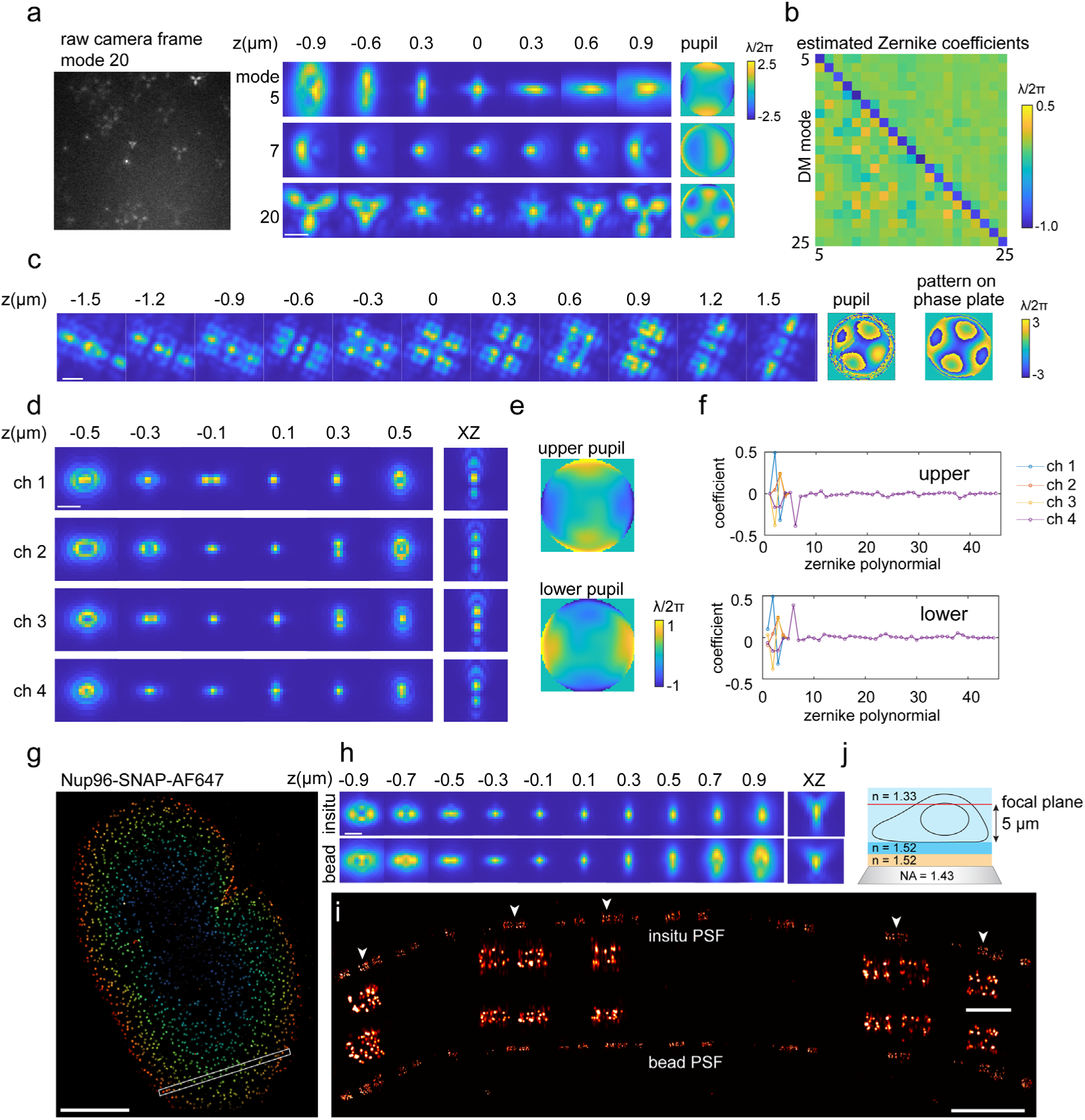
Modelling PSFs *in situ* from single blinking fluorophores. (a) uiPSF was used to determine a PSF model from single blinking fluorophores when specific Zernike aberrations were applied by a deformable mirror (DM) with a magnitude of −*λ*/2*π*. (b) The estimated Zernike coefficients from single blinking fluorophores for 25 different Zernike aberrations applied by the DM. Samples for a,b are microtubules labeled with AF647. (c) Tetrapod PSF model generated from a phase plate, the pupil-image based PSF model is estimated from single blinking fluorophores. The sample is TOMM20 labeled with AF647. (d-f) Estimation of a Zernike-based 4Pi-PSF model (estimated PSF model (d) pupil function (e) and Zernike coefficients (f)) from blinking single fluorophores. The sample is Nup96-mMaple in U2OS cells. (g-i) Comparison of PSF models from SMLM data and from bead data for astigmatic 3D SMLM (h). (j) The sample is Nup96-AF647 imaged at 5 µm above the coverslip using an oil immersion objective. (i) XZ view of the selected region in (g), a few zoom-in NPCs under the white arrows shows consistent ring separation for the in-situ PSF model. Scale bars, 1 µm (a,c,d,h,i), 5 µm (g), 200 nm (i, zoom in).

INSPR^41^ is a recent software that determines Zernike coefficients from blinking fluorophores after phase retrieval with a modified GS algorithm. In comparison to uiPSF, it resulted in larger off-diagonal terms (ED Fig. 6a-b). As a further comparison, we imaged AF647 dye molecules on coverslip at different z positions, extracted *in situ* PSF models (ED Fig. 6c-d) and used those to fit the single moleucles again. Our *in situ* PSF estimation method exhibited a linear relationship between the fitted z-position and the objective z-position, whereas INSPR resulted in a discontinuity, suggesting a mismatch of the PSF model from the data.

As the aberrations generated by the DM are usually smooth across the pupil plane, they can be modeled well by Zernike polynomials. However, some PSFs generated with a phase plate or a spatial light modulator (SLM) can have discontinuities, requiring an image-based pupil function (SI Fig. 19). We demonstrated pupil-image based *in situ* PSF estimation on Tetrapod PSF patterns generated from a phase plate (Fig.3c and ED Fig.7). Due to discontinuities of the phase pattern (Fig. 3c), the Zernike-based *in situ* PSF modelling requires more accurate initial values and the obtained PSF model is blurrier than the one from pupil-image based modelling (ED Fig. 7).

Our *in situ* PSF modelling method can easily be extended to multiple channels. We demonstrated this on a 4Pi-SMLM system by imaging Nup96 in U2OS cells, using the relatively dim photo-convertible fluorescent protein mMaple as a label (Fig. 3d-f and SI Fig. 20). We used uiPSF to extract the Zernike coefficients for the upper and lower interference arm independently and included a relative piston phase between the two arms (Fig. 3f). To reduce the effect of data noise, we linked the Zernike coefficients between the 4 channels in each arm, except the tip, tilt and defocus, which describes the relative x, y and z shifts between the two objectives in each channel. The transformations among the channels are also obtained during the inverse modelling process.

We next tested the performance of uiPSF for *in situ* PSF learning at the presence of large sample-induced aberrations (Fig. 3g-i). We imaged the Nup96 labeled with AF647 at the top of the nucleus, ∼5 µm above the coverslip, using an oil immersion objective lens (NA=1.43) and standard imaging buffer (n=1.35) (Fig. 3j and Methods). The refractive index mismatch resulted in strong aberrations and clear differences between the *in situ* PSF and bead-based PSF models (Fig. 3h and ED Fig. 8). The reconstructed NPCs show consistent ring distances for the *in situ* PSF, but strong squeezing in the z-direction for the bead PSF at larger depth (Fig. 3i). To quantitively evaluate the accuracy of both the bead PSF and *in situ* PSF models, we measured the distance between the two rings in 3D across thousands of NPCs^61^ (ED Fig. 1) and found that the *in situ* PSF model indeed resulted in more accurate and consistent measurements of the ring distance. We confirmed the robustness of the *in situ* PSF model by imaging Nup96 at various depths by placing U2OS cells on the top coverslip of a sandwiched sample (∼25 µm between the two coverslips). The sample was imaged by a silicon oil objective lens (NA=1.35) and standard imaging buffer (n=1.35). Data from both the upper and lower nuclear envelopes demonstrated that the *in situ* PSF model resulted in consistent distances between the two rings (ED Fig. 1).

### Field-dependent aberrations

Up to now, we assumed that all beads or all fluorophores have identical emission patterns. However, this assumption is not generally fulfilled, as field-dependent (FD) aberrations lead to a PSF that is different at different positions in the FOV. This can be a major limitation for SMLM with large FOVs, which are important to increase the throughput of the slow SMLM imaging^62^. Here, we overcome this limitation by considering an FD-PSF model in the Zernike representation. Instead of estimating one set of Zernike coefficients, we represent every Zernike aberration as a map where each pixel value represents the Zernike coefficient at the center of the subregion. In the forward model, the Zernike coefficients for each emitter are calculated from the linear interpolation of the aberration maps at the pixel coordinates of the bead or fluorophore. During the estimation, we impose a certain degree of smoothness to the aberration maps, leading to meaningful values even when no emitter was found in a subregion (SI Note 3).

We first validated modelling of FD-PSFs using bead images across a FOV of 180 µm × 180 µm. Since the bead images were taken at equal axial spacing, the x, y, z positions and background parameter of all images in each bead stack were linked. In contrast, the photon parameter of each bead stack was fitted individually in each frame to account for bleaching and intensity fluctuations. The aberration maps across the whole FOV were directly estimated by maximum likelihood estimation using all beads simultaneously by globally optimizing the aberration maps. The resulting aberration maps were much smoother than those obtained by FD-DeepLoc^42^, which fits every bead individually. Thus, uiPSF reduced artifacts introduced by abnormal bead images (e.g. aggregated beads, non-uniform beads) which often appear in the single bead fitting process (Fig. 4a, SI Fig. 21a). Finally, we validated the FD-PSF model by fitting the z position of each bead and compared them to the objective positions and could show that localization biases were greatly reduced compared to fitting with the average PSF model (Fig. 4b,c and SI Fig. 21b).

**Figure 4.**
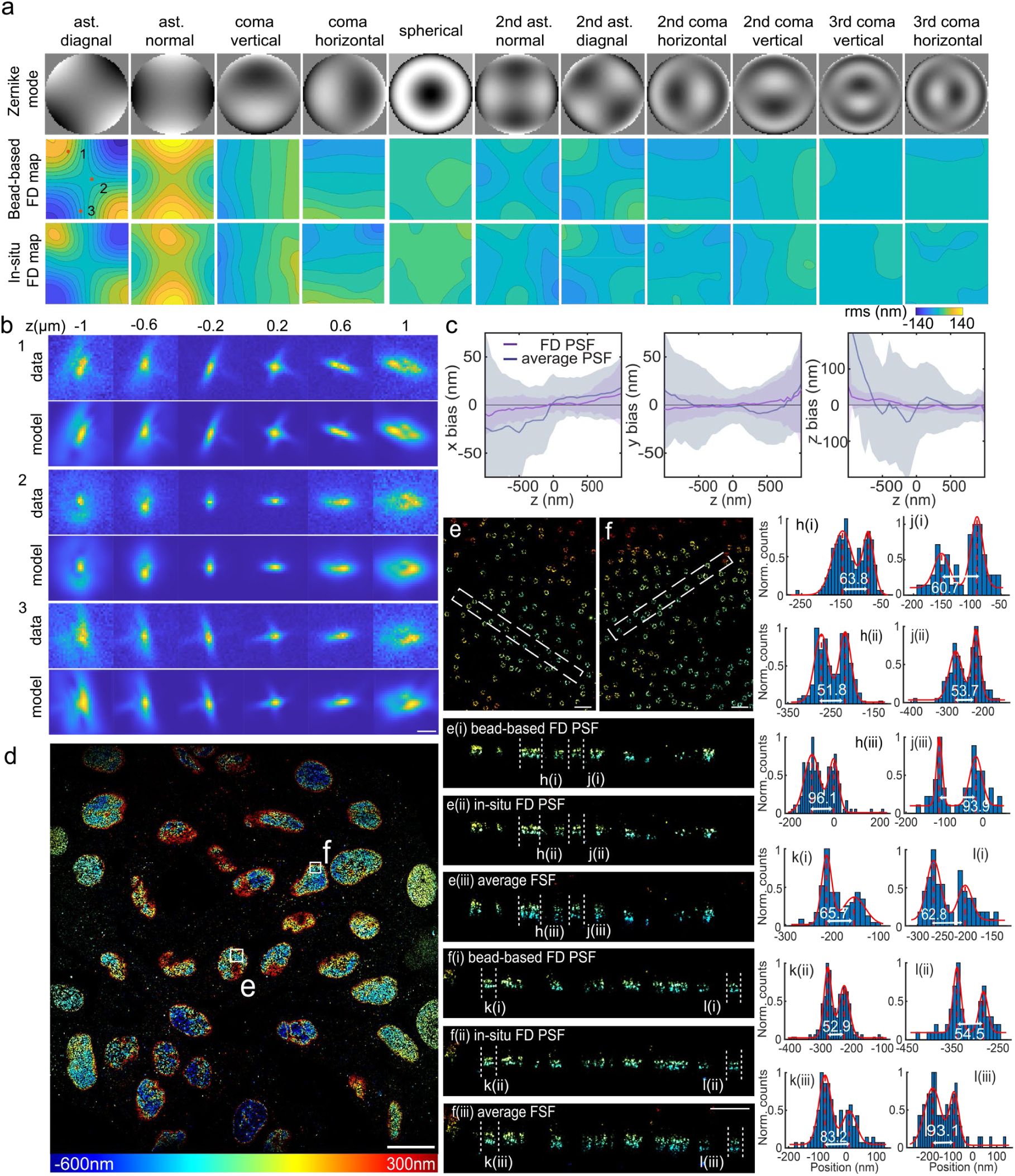
Field-dependent PSF modeling. (a) FD map of Zernike coefficients obtained from beads and single fluorophores in an FOV of about 180 µm × 180 µm. (b) uiPSF successfully recovers FD PSFs from beads at different positions indicated in (a). (c) Comparison of localization bias in x, y, and z obtained through the localization of beads data using the FD PSF and an average PSF. (d-l) in-situ FD PSF. (d) Nup96-AF647 in U2OS cells imaged, fitted with FD-DeepLoc using a FD *in situ* PSF model. The different colors represent various z positions. (e) and (f) zooms as indicated in (d). e(i-iii) and f(i-iii) are side view reconstructions of the selected region indicated in (e-f) and intensity profiles through individual NPCs (h-l) demonstrate the higher quality for the FD *in situ* PSF. Scale bars, 50 µm (a), 1 µm (b), 20 µm (d), and 0.5 µm (e-f).

Next, we showed that uiPSF estimates an *in situ* FD-PSF model from single-molecule blinking data. From imaging NPCs in many cells across a FOV of 180 µm × 180 µm, we observed a high degree of similarity in the estimated aberration maps between bead and *in situ* PSF models (Fig. 4a). We used the deep-learning based fitter, FD-DeepLoc^42^, to reconstruct the NPCs with *in situ* and bead FD-PSF models and found that both FD-PSF models lead to higher quality reconstructions compared to that of using an averaged PSF model (Fig. 4d-l).

### Microscope characterization

uiPSF is a useful toolbox for characterization and optimization of a microscope system, as the estimated PSFs, Zernike coefficients and aberration maps provide useful information on the aberrations of the imaging system. uiPSF provides a demo notebook that takes a Zernike-based PSF model as an input and displays major aberrations modes, the Strehl ratio and full width of half maximum (FWHM) of the PSF model for each channel. For a 4Pi-SMLM system, it will report the modulation depth instead of the Strehl ratio. For FD-PSF modelling, the notebook will display the aberration maps of the major aberrations, as well as the Strehl ratio and FWHM maps. The user can use this information to minimize system aberrations, or, if adaptive optics (AO) systems (deformable mirror or spatial light modulator) are integrated, to directly compensate for system or sample-induced aberrations.

We demonstrated this by imaging NPCs at the upper nuclear envelope, ∼4 µm above the coverslip, ED Fig.9a). Spherical aberrations caused by the refractive index mismatch, reduced localization precisions, making it difficult to resolve the two-layer ring structure of the NPC without aberration correction (ED Fig.9b-c). By employing *in situ* PSF modelling, we determined the Zernike aberrations and directly fed them back to the DM for AO correction, resulting in a significant improvement in the quality of the reconstructed NPCs (ED Fig.9d).

## Conclusion

We developed uiPSF, a modular toolbox that reconstructs accurate PSF models for most fluorescence microscopes, including single channel, multi-channel, 4Pi and lattice light-sheet systems, and demonstrated its use for SMLM. It can parameterize the PSFs in the form of 3D matrices, Zernike-based and image-based pupil functions. It can also account for various experimental imperfections such as field-dependent aberrations, the finite size of beads or camera pixels, intensity fluctuations, drifts, and shot noise. This versatility in terms of representations and extensions on one hand allows for optimizing the approach for highest robustness and accuracy, but also might be overwhelming for non-experts. Here we give a general guideline for selecting PSF representation and extensions (SI Note): 1) When bead data exists and when SMLM data were collected near the coverslip, we recommend using voxel-based learning (i.e., inverse modelling) if the number of beads is sufficient or Zernike-based learning if SNR is low. 2) For SMLM data at large depths with index mismatched buffer, or when bead data is not available, *in situ* PSF learning is always recommended. However, as discussed above, *in situ* PSF learning requires single fluorophores spanning a large axial range, thus if most emitters are located at similar z positions, *in situ* PSF learning might fail. 3) Pupil-image based learning is recommended only when the pupil function cannot be presented by Zernike polynomials. 4) For pupil-based learning (includes both Zernike-based and pupil-image based methods), we always recommend using the vectorial PSF model if computational resources permit, otherwise the scalar PSF model can be used for faster learning. 5) Voxel-based learning only works for bead data. Frame-wise intensity and lateral drift are not required, unless bead data exhibit large intensity fluctuation or lateral shifts between frames. 6) For pupil-based learning, an extra blurring should be always learned or set. 7) Oversampling and binning is usually not required. 8) The bead size can be ignored when bead is smaller than 100 nm.

The robustness of uiPSF allows extracting PSFs not only from bead stacks, but directly from the images of single blinking fluorophores, the raw data of SMLM. This not only renders a cumbersome bead calibration obsolete, but it also improves the accuracy of the PSF model, as the PSF is extracted precisely where it is measured and can include sample-induced aberrations.

The modularity of the software allows for simple extensions of the forward model in the future to include for instance fixed dipoles for single-molecule orientation microscopy^57^, polarized detection (e.g. using a polarized beam splitter in the emission path), spectral SMLM^63,64^, and imaging systems using structured illuminations, such as confocal, image scanning microscopy, structured illumination microscopy, SIMFLUX methods^65–67^, and MINFLUX^68^.

To make uiPSF easily accessible, we developed a user-friendly, extendable, and performant software in Python/TensorFlow that can be used via simple Jupyter Notebooks and integrates readily into existing software for SMLM analysis^23,42,58,69,70^. Computation times depend on the number of beads or single molecules used to learn the PSF, as well as on the representation, extensions and number of detection channels, but typically lie between 40 s (single-channel voxelated PSF from 20 beads) and 35 min (4Pi *in situ* model from 10000 single fluorophores) on a consumer GPU (Nvidia RTX3080). uiPSF will enable many microscopists to quantify and optimize the performance of their microscopes and to perform SMLM with improved accuracy.

## Acknowledgements

We thank Y.E. Katrukha and L.C. Kapitein from Utrecht University, Netherlands; B. Hajj and L. Regnier from Institut Curie, France; K.J.A. Martens from Bonn University, Germany; A. Tschanz from European Molecular Biology Laboratory, Germany; S. Khan and S. Pani from the University of New Mexico, USA; Y. Li from Purdue University, USA for testing the software. We thank X. Liu from the University of Hong Kong for valuable suggestions on inverse modelling. This work was supported by the European Research Council (grant no. ERC CoG724489 to J.R.) and the European Molecular Biology Laboratory. This work was conducted with support from the University of New Mexico Office of the Vice President for Research Program for Enhancing Research Capacity. This research was supported by grants from NVIDIA and utilized an NVIDIA A6000 GPU. S.L. was supported by EMBL ARISE fellowship no. 945405. S.L. and K.A.L. were supported by NIH Grant 1R01GM140284. Y.L. was supported by National Natural Science Foundation of China (62375116); Key Technology Research and Development Program of Shandong (2021CXGC010212); Shenzhen Science and Technology Innovation Commission (Grant No. JCYJ20220818100416036 and KQTD20200820113012029); Guangdong Provincial Key Laboratory of Advanced Biomaterials (2022B1212010003); Startup grant from Southern University of Science and Technology. L.-R.M. was supported by the German Federal Ministry of Education and Research (BMBF; project SIMALESAM, FKZ 01|S21055A-B). R.H. was supported by the German research foundation (Deutsche Forschungsgemeinschaft, DFG, project TR 1278, TP C04, 316213987). C.K. was supported by the German research foundation (Deutsche Forschungsgemeinschaft, DFG, project SFB TR166 ‘‘Receptorlight’’, TP B05). This project has received funding from the European Union’s Horizon 2020 research and innovation program under grant agreement No. 802567-ERC-Five-Dimensional Localization Microscopy for Sub-Cellular Dynamics (to Y.S), and from the ISRAEL SCIENCE FOUNDATION (grant No. 450/18 to Y.S). Y.S. was supported by the Zuckerman Foundation.

## Data availability

Example data for uiPSF is available at https://zenodo.org/records/10027718.

## Code availability

uiPSF is available at https://github.com/ries-lab/uiPSF.

## Contributions

J.R., Y.L., R.H. and Y.S. conceived the project. J.R., Y.L., and K.A.L. provided supervision. S.L., J.H., J.C., C.K. and L.-R.M. wrote the software. S.L. and J.C. analyzed the data. S.L., J.C., J.R., B.F. and D.X. acquired the data. S.L., J.C., Y.L. and J.R. wrote the paper with input from all authors.

## Methods

### Microscopes

#### 4Pi-SMLM

The 4Pi-SMLM system is a custom-built microscope as described previously^59,71^. It consists of two opposingly configurated objectives (silicone oil objective, NA = 1.35, UPLSAPO100XS, Olympus), collecting fluorescence emission into both upper and lower emission arms. Four lasers 405 nm (iBEAM-SMART-405-S, 150 mW, TOPTICA Photonics), 488 nm (iBEAM-SMART-488-S-HP, 200 mW, TOPTICA Photonics), 561 nm (2RU-VFL-P-1500-560-B1R, MPB Communications) and 642 nm (2RU-VFL-2000-642-B1R, MPB Communications) were coupled into a single mode fiber for fluorescence imaging. The lasers were then filtered through a clean-up filter (ZET405/488/561/640xv2, Chroma) and reflected by a quadband dichroic (Di03-R405/488/561/635, Semrock) into the lower objective. The fluorescence emission was collected by both objectives and filtered by a quadband emission filter (FF01-432/515/595/730-25, Semrock). A band pass emission filter (FF01-600/37-25, Semrock) was also used before the camera for imaging Nup96-mMaple. The microscope is controlled through a LabView-based software. Two deformable mirrors (Multi-DM 5.5, Boston Micromachines), one in each emission arm, were used to minimize systematic aberrations and apply astigmatism aberration. The DMs were calibrated using a Python-based software to generate accurate Zernike aberrations.

#### Dual-color ratiometric imaging systems

The dual-color SMLM system is a custom-built microscope as described previously^58^. We used the same microscope for both dual-color and single-color imaging of NPCs (Fig. 2a-d, Fig. 3g-i). It equipped with a high NA oil immersion objective (160x, 1.43-NA oil immersion, Leica, Wetzlar, Germany). A commercial laser box (LightHub®, Omicron-Laserage Laserprodukte, Dudenhofen, Germany) equipped with Luxx 405, 488 and 638, Cobolt 561 lasers and an additional 640 nm booster laser (iBeam Smart, Toptica, Munich, Germany) were combined for wide field illumination. Lasers were focused onto a speckle reducer (LSR-3005-17S-VIS, Optotune, Dietikon, Switzerland) and coupled into a multi-mode fiber (M105L02S-A, Thorlabs, Newton, NJ, USA). The lasers were triggered using an FPGA (Mojo, Embedded Micro, Denver, CO, USA) allowing microsecond pulsing control of lasers. The output of the fiber was magnified by an achromatic lens and imaged into the sample plane. A laser clean-up filter (390/482/563/640 HC Quad, AHF, Tübingen, Germany) was placed in the excitation beam path to remove the fluorescence generated by the fiber. The focus of microscope was stabilized by a 785 nm infrared laser (iBeam Smart, Toptica, Munich, Germany) that was projected through the objective and reflected by the coverslip onto a quadrant photodiode, which was used as closed-loop feedback signal to the objective piezo stage (P-726 PIFOC, Physik Instrument, Karlsruhe, Germany). The astigmatic 3D imaging was acquired using a cylindrical lens (f = 1,000 mm; LJ1516L1-A, Thorlabs) to introduce astigmatism. For astigmatic dual-color imaging with AF647 and CF680, the fluorescence was split by a 665 nm long pass dichroic (ET665lp, Chroma), filtered by a 685/70 (ET685/70 m, Chroma) bandpass filter for the transmitted light and a 676/37 (FF01-676/37-25, Semrock) bandpass filter for the reflected light. An EMCCD camera (Evolve512D, Photometrics, Tucson, AZ, USA) was used to collect final fluorescence. The microscope is entirely controlled by Micro-Manager41 using htSMLM, a custom EMU plugin^72^.

#### Large FOV imaging system

Large FOV imaging system is a custom-built microscope designed for high-throughput SMLM imaging at room temperature (24 °C). The sample illumination is achieved by combining lasers with a multimode fiber (WFR CT200x200/230x230/440/620/1100N, NA 0.22, CeramOptec). The lasers pass through a laser cleanup filter (ZET405/488/561/640xv2, Chroma) and a main dichroic (ZT405/488/561/640rpcxt-UF2, Chroma) before entering the objective for sample illumination. The fluorescence emitted by the sample is collected by a high-NA objective (NA 1.5, UPLAPO ×100 OHR) and filtered using a quad-band emission filter (ZET405/488/561/640mv2, Chroma). A cylindrical lens is placed at a specific distance in front of the imaging camera for astigmatism-based 3D imaging. Prior to reaching the sCMOS camera (PRIME 95B, Teledyne Photometrics), the residual laser is eliminated using a band-pass filter (ET680/40m, Chroma). A 785 nm near-infrared laser (iBEAM-SMART-785-S, Toptica Photonics) is introduced to stabilize the sample focus using a dichroic mirror (FF750-SDi02, Semrock). The reflected laser is detected by a quadrant photodiode (SD197-23-21-041, Advanced Photonix), and the position-dependent output voltage serves as feedback for the z-axis of the objective (P-726.1CD, Physik Instrumente). The microscope is controlled using Micro-Manager integrated with EMU^72^. Typically, we acquire 50,000 to 100,000 frames with exposure times ranging from 15 to 25 ms.

#### Adaptive optics single-channel SMLM system

Adaptive optics single-channel SMLM system is a custom-built microscope as described before^73^. A combination of laser beams of different wavelengths is used in the experiment. A 405 nm laser (iBEAM-SMART-405-S, 150 mW, TOPTICA Photonics), a 488 nm laser (iBEAM-SMART-488-S-HP, 200 mW, TOPTICA Photonics), a 561 nm laser (MGL-FN-561nm, 300mW, CNI), and a 640 nm laser (iBEAM-SMART-640-S-HP, 200mW, TOPTICA Photonics) are combined using a fiber coupler (PAF2-A4A, Thorlabs) and transmitted through a single-mode fiber (P3-405BPM-FC-2, Thorlabs). The position of the fiber output can be adjusted using a translation stage to achieve different illumination angles. To eliminate fluorescence induced by the fiber, the illumination beam is filtered using a laser clean-up filter (ZET405/488/561/640xv2, Chroma). The main dichroic mirror (ZT405/488/561/640rpcxt-UF2, Chroma) reflects the beam, which then enters the objective to illuminate the sample. The emitted fluorescence is collected by a high NA objective (NA 1.35, UPLSAPO 100XS or NA 1.5, UPLAPO 100XOHR, Olympus) and imaged using the tube lens (TTL-180-A, Thorlabs). Two bandpass filters (NF03-405/488/561/635E-25 and FF01-676/37-25, Semrock) are used to separate the emitted fluorescence from the excitation laser. A deformable mirror (DM140A-35-P01, Boston Micromachines) is placed in the Fourier plane is employed for PSF engineering. To accurately determine the influence function of each actuator in the DM, we adopted the method outlined in reference^74^. Finally, the images are captured using an sCMOS camera (ORCA-Flash4.0 V3, HAMAMATSU) with a pixel size of 108 nm on the sample. Typically, 50,000-100,000 frames are acquired with a 15-25 ms exposure time.

#### Optical system for generating Tetrapod PSFs

Plan Apo 100X/1.45 NA Nikon objective was fitted on a Nikon Eclipse-Ti inverted fluorescence microscope with a sCMOS camera (Prime95B, Photometrics). The sample was illuminated by a high-intensity 638 nm 2000 mW red dot laser module, filtered through a 25 µm pinhole (Thorlabs) and low intensity 405nm laser (iChrome MLE, Toptica, <5mw). The emission path was extended by a 4f system consisting of two achromatic doublet lenses (focal length 15 cm) and a dielectric phase mask (a 4 µm z-range Tetrapod, photolithographically fabricated) was placed in the conjugate back focal plane. The emission light was filtered through a 500 nm long pass dichroic and a 650 nm long pass (Chroma).

### Sample preparation

#### Cell culture

COS-7 cells (catalog no. 100040, BNCC) were grown in DMEM (catalog no. 10569, Gibco) containing 10% (v/v) fetal bovine serum (FBS; catalog no. 10099-141C, Gibco), 100 U ml−1 penicillin and 100 μg ml−1 streptomycin (PS; catalog no. 15140-122, Gibco). U2OS cells (Nup96-SNAP no. 300444, Cell Line Services) were grown in DMEM containing 10% (v/v) FBS, 1× PS and 1× MEM non-essential amino acids (NEAAs; catalog no. 11140-050, Gibco). Cells were cultured in a humidified atmosphere with 5% CO2 at 37 °C and passaged every 2 or 3 d. Before cell plating, high-precision 25-mm round-glass coverslips (no. 1.5H, catalog no. CG15XH, Thorlabs) were cleaned by sequentially sonicating in 1 M potassium hydroxide (KOH), Milli-Q water and ethanol and finally irradiated under ultraviolet light for 30 min. For super-resolution imaging, COS-7 and U2OS cells were cultured on the clean coverslips for 2 d with a confluency of 80–90%. All cells were cultured at 37 °C in a humidified atmosphere containing 5% CO2. Mycoplasma detection was conducted routinely to ensure Mycoplasma-free conditions throughout the study.

#### Nup96-mMaple in U2OS cells

U2OS-Nup96-mMaple cells were prepared as previously reported. Cells were prefixed for 30 s in 2.4% (w/v) formaldehyde in PBS and then permeabilized with 0.4% (v/v) Triton X-100 in PBS for 3 min. To complete the fixation, samples were incubated for 30 min in 2.4% (w/v) formaldehyde in PBS. Formaldehyde was subsequently quenched in 100 mM NH4Cl in PBS for 5 min before washing the coverslip twice for 5 min in PBS.

#### Nup96-SNAP-AF647 in U2OS cells

To label Nup96, U2OS-Nup96-SNAP cells were prepared as previously reported. Cells were prefixed in 2.4% paraformaldehyde (PFA) for 30 s, permeabilized in 0.4% Triton X-100 for 3 min and subsequently fixed in 2.4% PFA for 30 min. Then, cells were quenched in 0.1 M NH4Cl for 5 min and washed twice with PBS. To decrease unspecific binding, cells were blocked for 30 min with Image-iT FX Signal Enhancer (catalog no. I36933, Invitrogen). For labeling, cells were incubated in dye solution (1 μM SNAP-tag ligand BG-AF647 (catalog no. S9136S, New England Biolabs), 1 mM dithiothreitol (catalog no. 1111GR005, BioFroxx) and 0.5% bovine serum albumin (BSA) in PBS) for 2 h and washed three times in PBS for 5 min each to remove excess dyes. Last, cells were post-fixed with 4% PFA for 10 min, washed with PBS three times for 3 min each and stored at 4 °C until imaged.

#### TOMM20-AF647 imaging with Tetrapod PSFs

Sample preparation included cleaning coverslips (22 × 22 × 0.17 mm) in an ultrasonic bath with 5% Decon90 at 60°C for 30 min, then washing with water and incubating in ethanol absolute for 30 min. Finally sterilized with 70% filtered ethanol for another 30 min. COS7 cells were grown for 24 hours on the coverslips in 6-well plate in phenol red free Dulbecco’s Modified Eagle Medium (DMEM) With 1g/l D-Glucose (Low Glucose) supplemented with 10% Fetal bovine serum, 100 U/ml penicillin 100 ug/ml streptomycin and 2 mM glutamine, at 37oC, and 5% CO2. The cells were fixed with 4% paraformaldehyde and 0.2% glutaraldehyde in PBS, pH 6.2, for 45 min, washed and incubated in 0.3M glycine/PBS solution for 10 minutes. The coverslips were transferred into a clean 6-well plate and incubated in a blocking solution (10% goat serum, 3% BSA, 2.2% glycine, and 0.1% Triton-X in PBS, filtered with 0.45 µm PVDF filter unit, Millex) for 2 hours at 4oC. The cells were then immunostained overnight at 4°C with anti TOMM20-AF647 antibody (diluted 1:230 in the blocking buffer) and then washed X5 with PBS. Finally, blinking buffer was made from 100 mM β-mercaptoethylamine hydrochloride, 20% sodium lactate, and 3% OxyFluor (Sigma, SAE0059). At the time of imaging, a PDMS chamber was attached to the glass coverslip containing the fixed cells with a second coverslip on top to prevent evaporation. The camera exposure time was 30 ms.

#### Fluorescence bead sample

The fluorescence beads used in this study include: 40 nm red bead (580/605 nm, Thermo Fisher, cat. no. F8793), 100 nm red bead (580/605 nm, Thermo Fisher, cat. no. F8801), 200 nm red bead (580/605 nm, Thermo Fisher, cat. no. F8810) and 100 nm TetraSpeck bead (Thermo Fisher Scientific, cat. no. T7279). For the single objective system, except the measurement of the Tetrapod PSF, the same preparation procedure is used. A clean 25 mm coverslip was incubated with 1 ml of bead dilution in 100 mM MgCl_2_ for 10 minutes. The coverslip was then washed with PBS for three times. The coverslip was then transferred to a custom sample holder and 1 ml of PBS was added to the coverslip. The sample was then mounted for imaging. The concentration of the bead dilution in above mentioned beads are 1:10^7^, 1:10^6^, 1:10^5^ and 1:10^6^ respectively. For measuring Tetrapod PSF, 40 nm red beads (580/605 nm, Thermo Fisher, cat. no. F8793) immobilized on the coverslip with 1% Polyvinyl alcohol were imaged.

#### Imaging buffer

Samples were imaged in refractive index matching buffer, including 50 mM Tris-HCl (pH 8.0), 10 mM NaCl, 10% (w/v) glucose, 0.5 mg/ml glucose oxidase (G7141, Sigma), 40 μg/ml catalase (C100, Sigma), 35 mM cysteamine and 28.5% (v/v) 2,2′-thiodiethanol (166782, Sigma). The refractive index of the final imaging buffer was n = 1.406.

### Data acquisition

#### Imaging of Nup96-mMaple on 4Pi-SMLM

The fixed Nup96-mMaple cells were incubated with 100 nm red fluorescent beads in a dilution of 1:10^6^ in 100 mM MgCl_2_ for 10 minutes. The cells were then washed three times with PBS. The coverslip was picked up with a tweezer and the excess buffer was removed by gently touching the edge of the coverslip on a Kimwipe tissue. The coverslip was then transferred to a custom sample holder, 200 µl imaging buffer (43.2% (w/w) sucrose in 50 mM Tris in D2O, pH 8.0) was added to the coverslip. A clean 24 mm coverslip was slowly placed on top of the imaging buffer. The top coverslip was slightly touched with the blunt side of a tweezer and the excess buffer was extracted with Kimwipes. The coverslips were then sealed with a two-component dental glue (Picodent, 1300 1000). After the dental glue solidifies in ∼20 minutes, the sample was mounted for imaging.

Before data collection, the deformable mirror (DM) in each arm was optimized to minimize the system aberrations. The optimization procedure is as follows: a z-stack of bead data was collected from each emission arm and the Zernike coefficients were retrieved using a GS-based phase retrieval algorithm. The corresponding correction voltage-map based on the retrieved Zernike coefficients were applied to the respective DM. This process was repeated 2-3 times until the obtained Zernike coefficients were within ±0.02 radian.

140,000 frames were collected at an exposure time of 25 ms, with a 561 nm excitation laser power of 2.3 kW/cm^2^, and a 405 nm activation laser power of 0-40 W/cm^2^ which was automatically adjusted to maintain the number of emitters per frame at ∼3.

#### Dual-color ratiometric imaging of Nup96 and WGA

The sample for dual-color imaging of Nup96 and wheat germ agglutinin (WGA) was prepared as previously reported^75^. Nup96-SNAP-tag cells (catalog no. 300444, CLS Cell Line Service) were rinsed twice with warm PBS. Prefixation was carried out in a 2.4% [w/v] formaldehyde in PBS solution for 40 s before the samples were permeabilized in 0.4% [v/v] Triton X-100 in PBS for 3 min. Complete fixation was carried out in 2.4% [w/v] formaldehyde in PBS for 30 min followed by three 5 min washing steps in PBS after fixation. Subsequently, the sample was incubated for 30 min with Image-iT FX Signal Enhancer (catalog no. I36933, Thermo Fisher Scientific) before staining with SNAP dye buffer (1 µM BG-AF647 (catalog no. S9136S, New England Biolabs) and 1 µM dithiothreitol in 0.5% [w/v] BSA in PBS) for 2 h at room temperature. To remove unbound dye, coverslips were washed three times for 5 min in PBS. For WGA staining, the sample was then incubated for 10 min with 400 ng ml−1 WGA-CF680 (catalog no. 29029-1, Biotium) in 100 mM Tris pH 8.0, 40 mM NaCl, and rinsed three times with PBS. Before imaging, samples were mounted on a custom sample holder in imaging buffers. The holder was sealed with parafilm.

#### Collection of SMLM data at various DM generated aberrations

We tested the accuracy of distortion wavefront estimation from a single-molecule blinking datasets (SI Fig. 20). The deformable mirrors were calibrated to introduce single-aberration modes based on Zernike polynomials (SI Note 5). We utilized 21 Zernike modes within the Fringe order, ranging from vertical astigmatism to third-order spherical aberration. Each mode was applied to distort fluorescent beads and single molecules with Zernike-based aberrations (amplitude ±1, units λ/2π) introduced by the DM. These aberrations were then used in the in situ PSF construction algorithm to globally fit the coefficients of the 21 Zernike modes post-acquisition. After processing the 21 Zernike modes, we created a heat map to illustrate the relationship between the DM input of the Zernike modes and the output amplitude of the algorithm.

#### Imaging of Nup96-AF647 at large FOV

Nup96-SNAP-AF647 labeled U2OS cells were imaged with the sCMOS camera over an FOV covering the full chip (1608×1608 pixels). The camera was operated under rolling shutter readout mode with an exposure time of 20 ms. 100,000 frames were acquired. The position of the cylindrical lens before camera was adjusted so that ∼80 nm astigmatism aberration was introduced to the system.

#### Collection of bead data on 4Pi-SMLM

Fluorescence beads were added to the cell sample as described in “Imaging of Nup96-mMaple on 4Pi-SMLM”. To reconstruct the 4Pi-PSF model from bead data, 20-30 bead stacks were collected. Each bead stack was collected by moving the sample stage from -0.5 µm to 0.5 µm with a step size of 50 nm, and three phase positions were collected at each z position by applying a piston phase to the DM at -2π/3, 0 2π/3. One frame per phase position was collected at an exposure time of 20 ms. The three phase positions were imaged with minimum delay where the DM change triggers the frame capture through software trigger.

#### Collection of bead data on single objective SMLM systems

The bead data for dual-color ratio-metric imaging and single-color imaging were collected in similar procedures. The fluorescence bead sample was prepared as described above. 10-30 bead stacks were collected where each bead stack was acquired by moving the sample stage from -1 µm to 1 µm with a step size of 10-50 nm. One frame per z position was collected at an exposure time of 10-35 ms. Beads embedded in agarose gel were collected by moving the sample stage from -1 µm to 5 µm with a step size of 20 nm. One frame per z position was collected at an exposure time of 50 ms.

#### uiPSF guided aberration correction with a DM

First, we acquired ∼1000 frame of single molecule data without adaptive optics (AO) correction. We then utilized uiPSF to perform *in situ* PSF modeling from single molecule data, resulting in Zernike aberrations. We applied 21 Zernike modes within the Fringe order, ranging from vertical astigmatism to third-order spherical aberration to the deformable mirror to compensate these aberrations using calibrated Zernike mirror modes (SI Note 5). Finally, we performed a complete single molecule data acquisition with AO correction activated.

## Extended Figures

**ED Fig 1.**
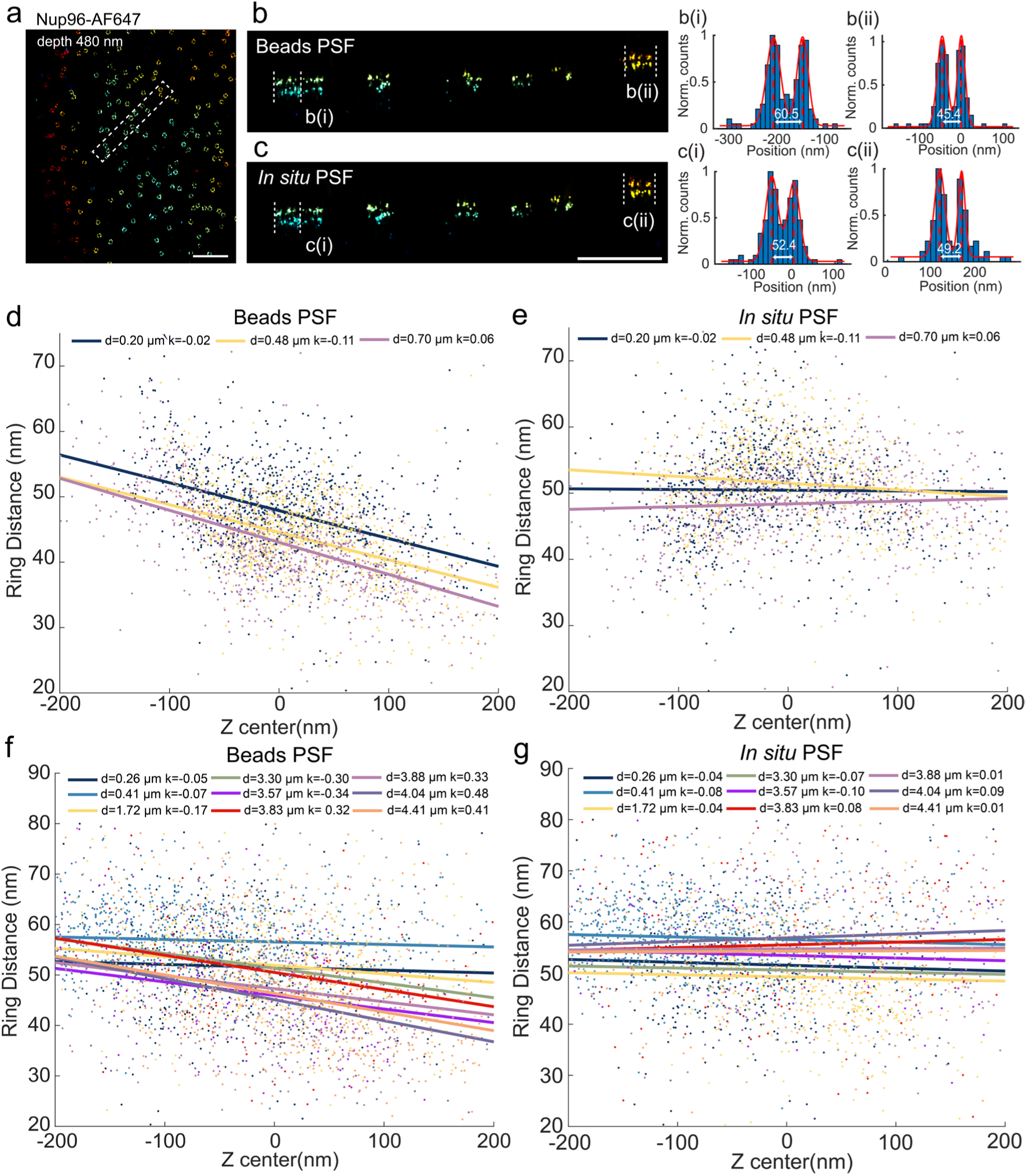
In situ PSF model reduces the local deformation in z measured by the ring distance of Nup96. (a) A subregion of the reconstructed Nup96 using the *in situ* PSF model. (b-c) XZ view of selected region in a from bead and *in situ* PSF models. (d) and (e) are the ring distance of Nup96 as a function of z-position at different depth reconstructed by using beads PSF model and in situ PSF model respectively. The points of various colors represent different imaging depths of Nup96. The straight lines represent the linear regression of the axial distance and z position for each imaging depth, with k representing the corresponding slope. An oil immersion objective lens (NA = 1.43) was used for imaging. (f) and (g) have the same meaning as (d) and (e), respectively, but imaged by a silicone oil immersion objective lens (NA = 1.35). Scale bars, 1 µm (a), 500 nm (b).

**ED Fig 2.**
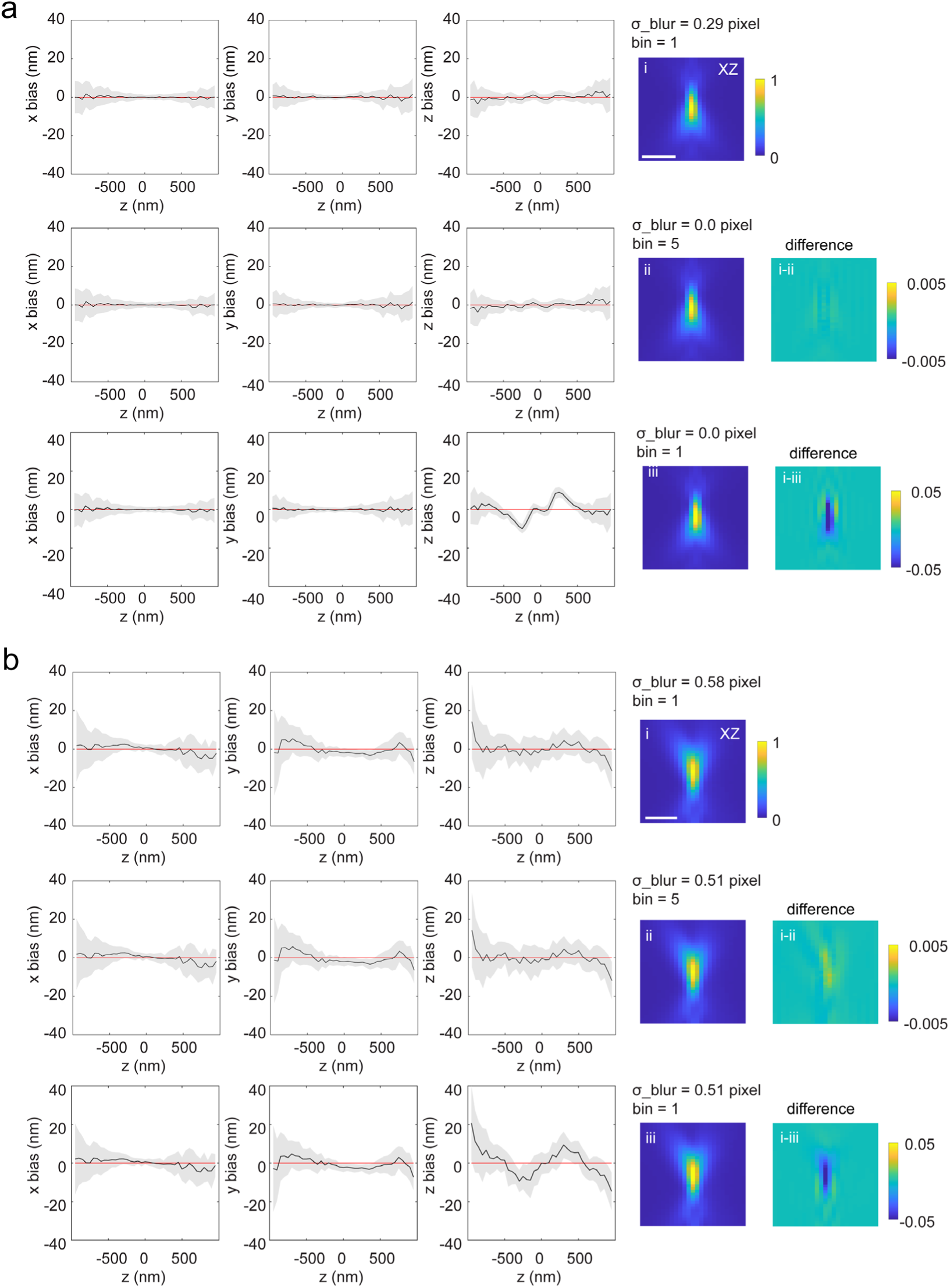
Effect of pixelation using the vectorial PSF modelling method. (a) Test on simulated data. Data were simulated from vectorial PSF model, with a binning of 5 in both x and y dimensions, and without extra blurring to the PSF model. (b) Test on experimental data. The estimated blur factor is slightly smaller when adding a binning of 5 in the forward model. However, the estimated PSF model and the localization bias are nearly identical with or without additional binning in the forward model. In contrast, under the same blur factor but without binning (bin=1), the axial bias increases dramatically and the estimated PSF model differs substantially. Scale bar, 1 µm.

**ED Fig 3.**
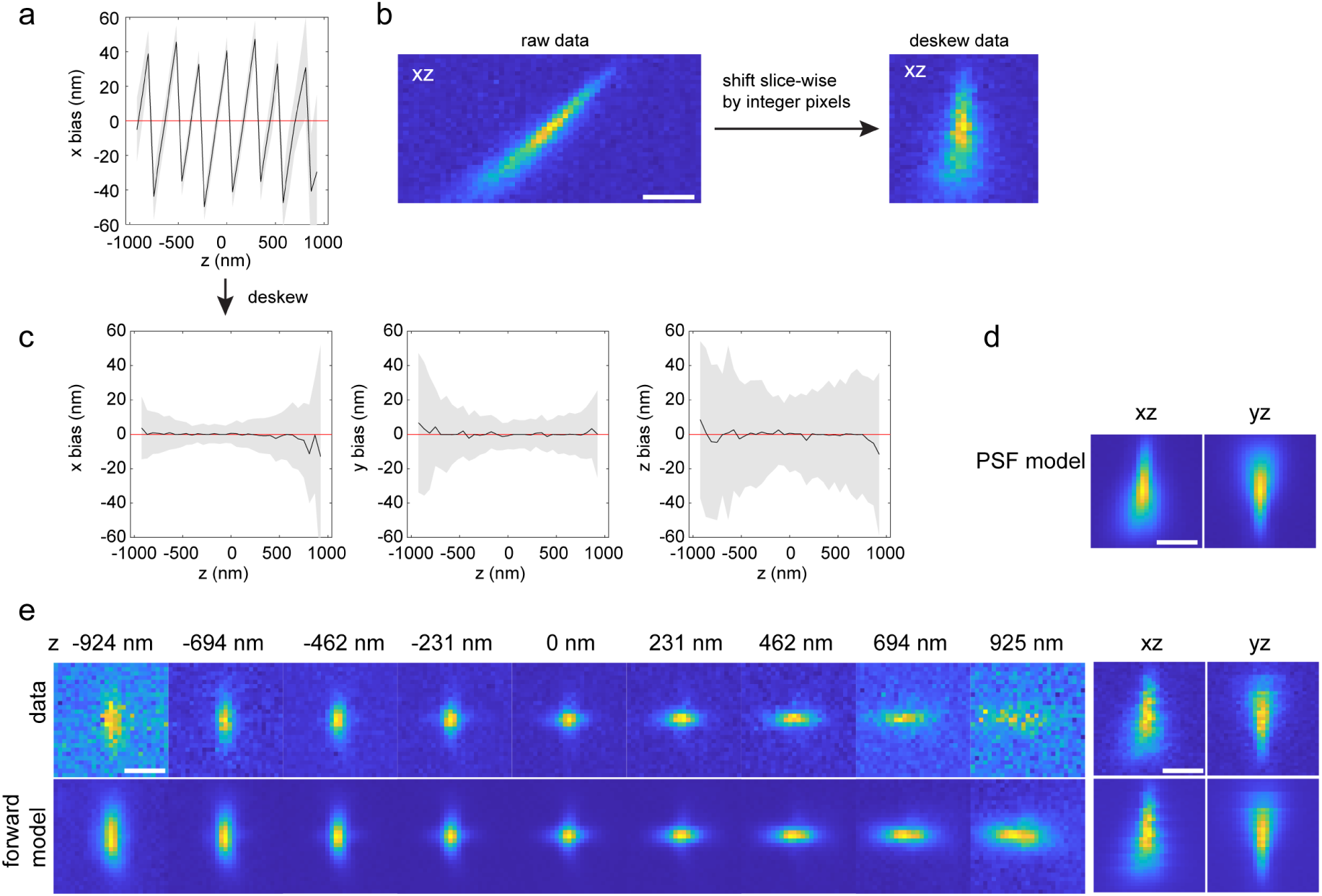
Estimation of voxel-based PSF model of a lattice light-sheet system from bead data. The data were collected by imaging beads in agarose gel at sample stage positions from -50 µm to 50 µm, with a step size of 50 nm. Translation of the sample stage will translate the beads both in x and z dimensions in the coordinate of the detection objective. For this data, translating the sample stage by 50 nm, will translate the beads by 40.8 nm in x and 28.9 nm in z relative to the detection objective. (a) Localization in x of the deskewed data used for inverse modelling. (b) The raw data stack of each bead was deskewed using the above translation relationship between x and z dimensions. Only shifts of integer pixels were applied to maintain the photon statistics of the raw data. The deskewed bead stacks were used for inverse modelling. The sub-pixel shifts were incorporated in the forward model so that the estimated PSF model has no skew as in d. (c) localization of the data used for inverse modelling, sub-pixel shifts were applied to the localization result in a to remove the skew effect. (d) Estimated PSF model. (e) Comparison of an example bead stack and its corresponding forward model. Scale bars, 1 µm (b,d,e).

**ED Fig. 4.**
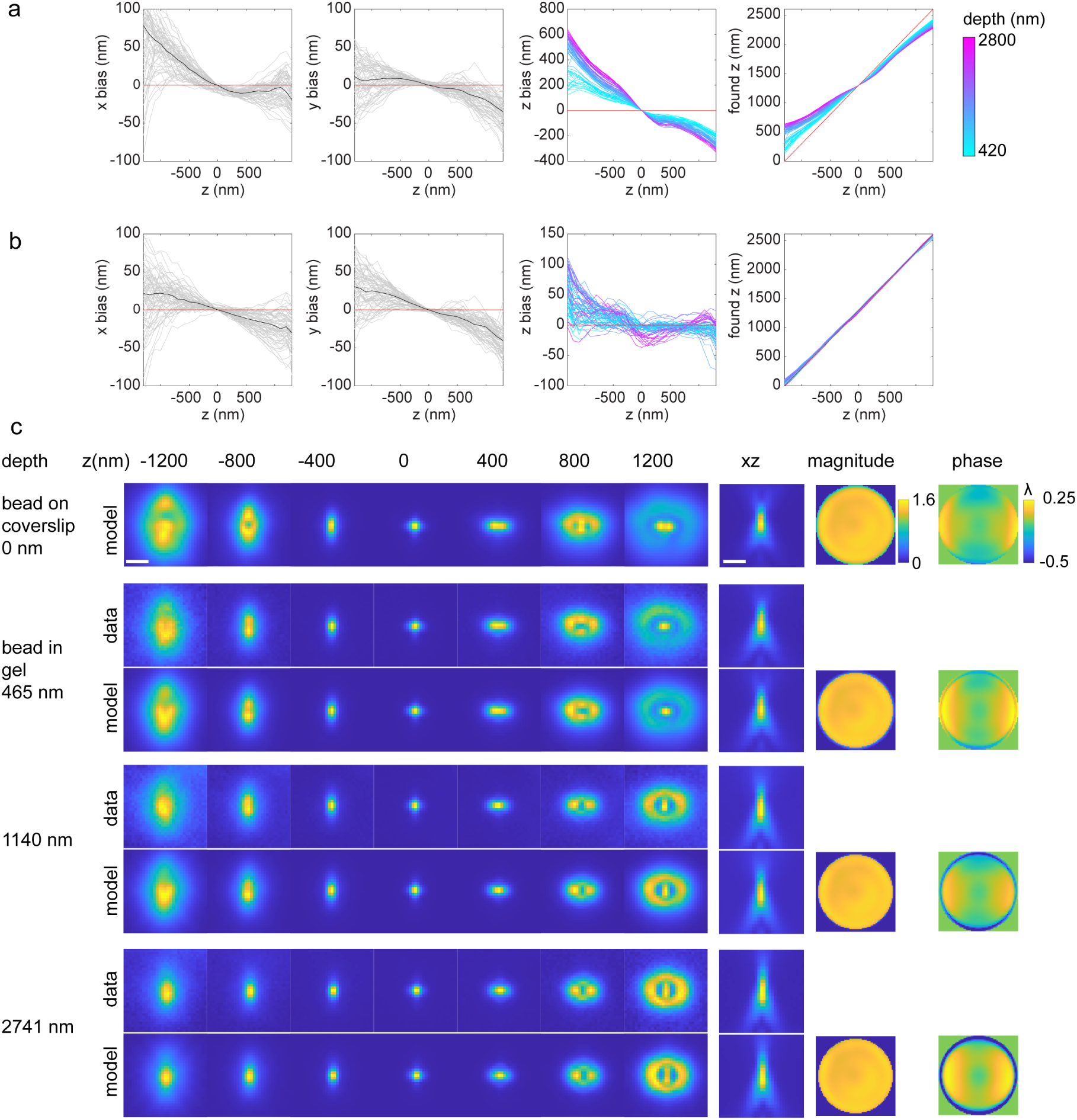
Incorporating refractive index mismatch aberrations in the estimated PSF model. Bead data at different imaging depth were collected by imaging beads in agarose gel at stage positions from -1 µm to 5 µm, with a step size of 20 nm. Bead data at the coverslip were collected at stage positions from -1.5 µm to 1.5 µm, with a step size of 10 nm. (a) Localization of beads in agarose gel using the PSF model estimated from beads at the coverslip. (b) Localization of beads in agarose gel using the PSF model estimated from beads at the coverslip and modified by adding an index mismatch aberration at the estimated imaging depth of each bead. Therefore, each bead stack was localized by its own modified PSF model. Depth is defined as the estimated emitter’s z position to the coverslip. (c) Comparison of the PSF models at the coverslip, bead data at different imaging depth and their corresponding PSF models. With increasing depth, the pupil size slightly decreases as the effective numerical aperture (NA) decreases from 1.43 to 1.33 (the refractive index of water), and the pupil phase show larger spherical aberration kind of patterns. Scale bars, 1 µm (c).

**ED Fig. 5.**
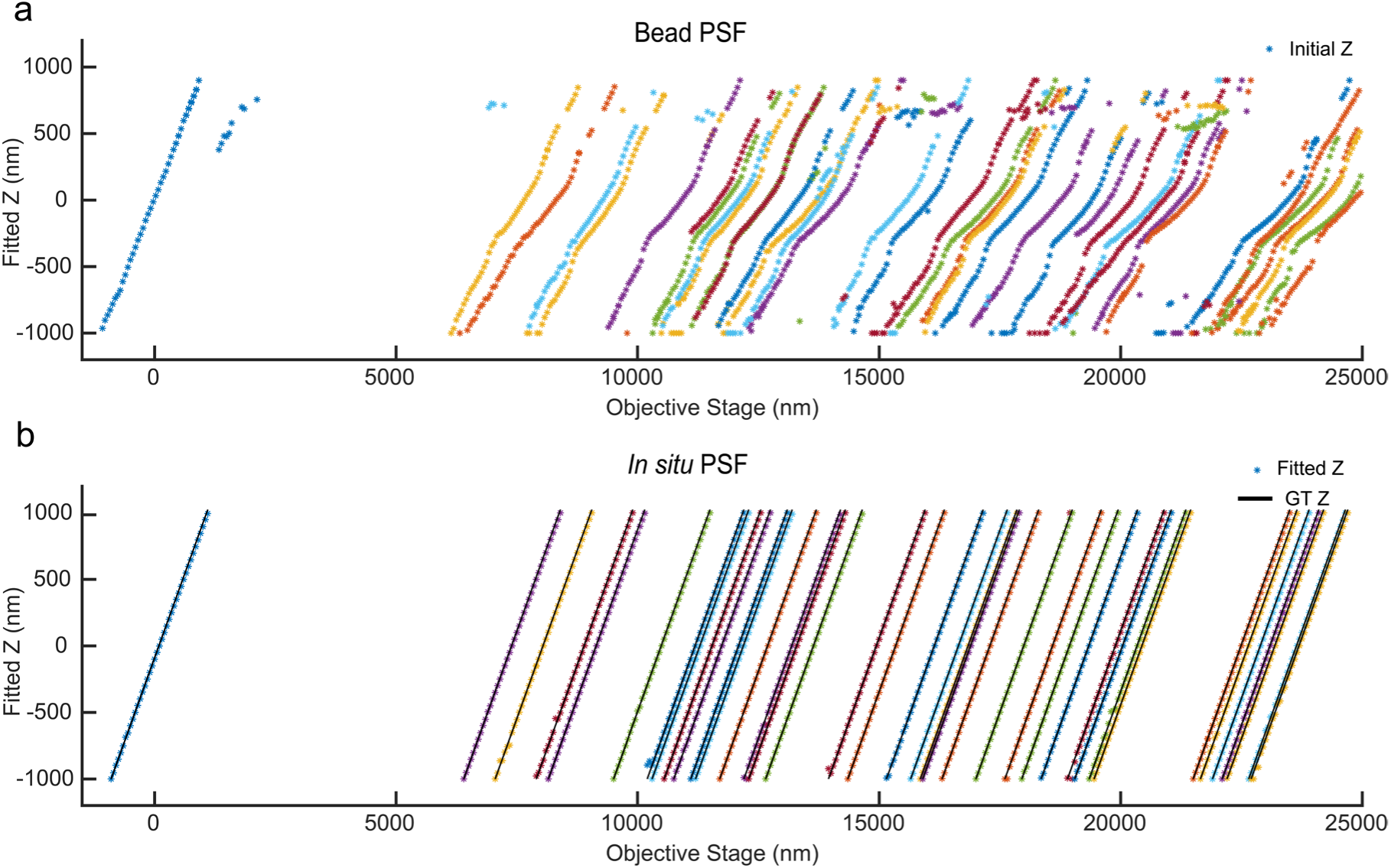
Validation of *in situ* PSF estimation using bead data in agarose gel. The images of these beads were collected at stage positions from -1 µm to 25 µm. For each bead, the per-frame z position was used as an independent variable to simulate the *in situ* single molecule data. (a) The PSF estimated from the beads on the cover glass was used to localize the bead data at different depths. (b) The *in situ* PSF estimated by uiPSF was used to localize the bead data at different depths. Compared to the PSF model estimated from the beads on the cover glass, the *in situ* PSF model fitted z-positions exhibit a good linear relationship with the objective stage positions.

**ED Fig 6.**
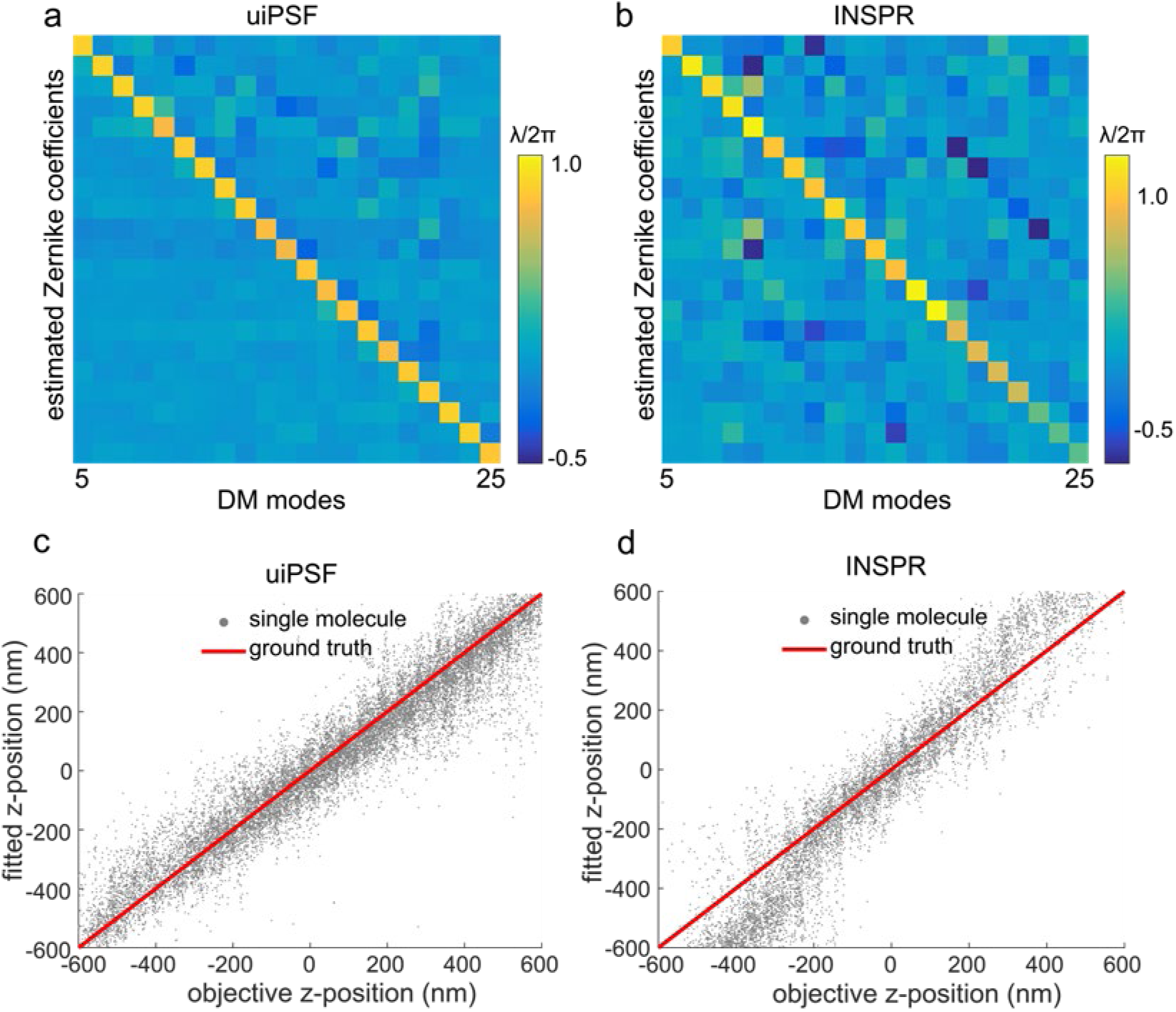
Comparison of Zernike-based PSF modelling using uiPSF and INSPR. Comparison between the input amplitude and the fitted amplitude of the *in situ* PSF modelling using the vectorial uiPSF (a) and the INSPR method (b) for 21 Zernike modes (fringe indices) of the deformable mirror input. Single-molecule data were collected when the DM was applied with 21 different Zernike modes separately. Such type of data was fitted by uiPSF and INSPR (b was adapted with permission from Fig. 3f in Xu et. al. ^41^). As shown in a and b, the Zernike aberrations returned by uiPSF have lower residual aberrations, compared to INSPR. Comparison of uiPSF (c) with the INSPR method (d) for fitted z-position and objective z-position. We imaged the dye AF647 immobilized on a coverslip and moved the objective the collect the z-stack imaging data. The z-position fitted by uiPSF is linearly related to the z-position of the objective (c), while there is a discontinuity at -300 nm for INSPR (d), indicating the mismatch of the INSPR PSF model.

**ED Fig 7.**
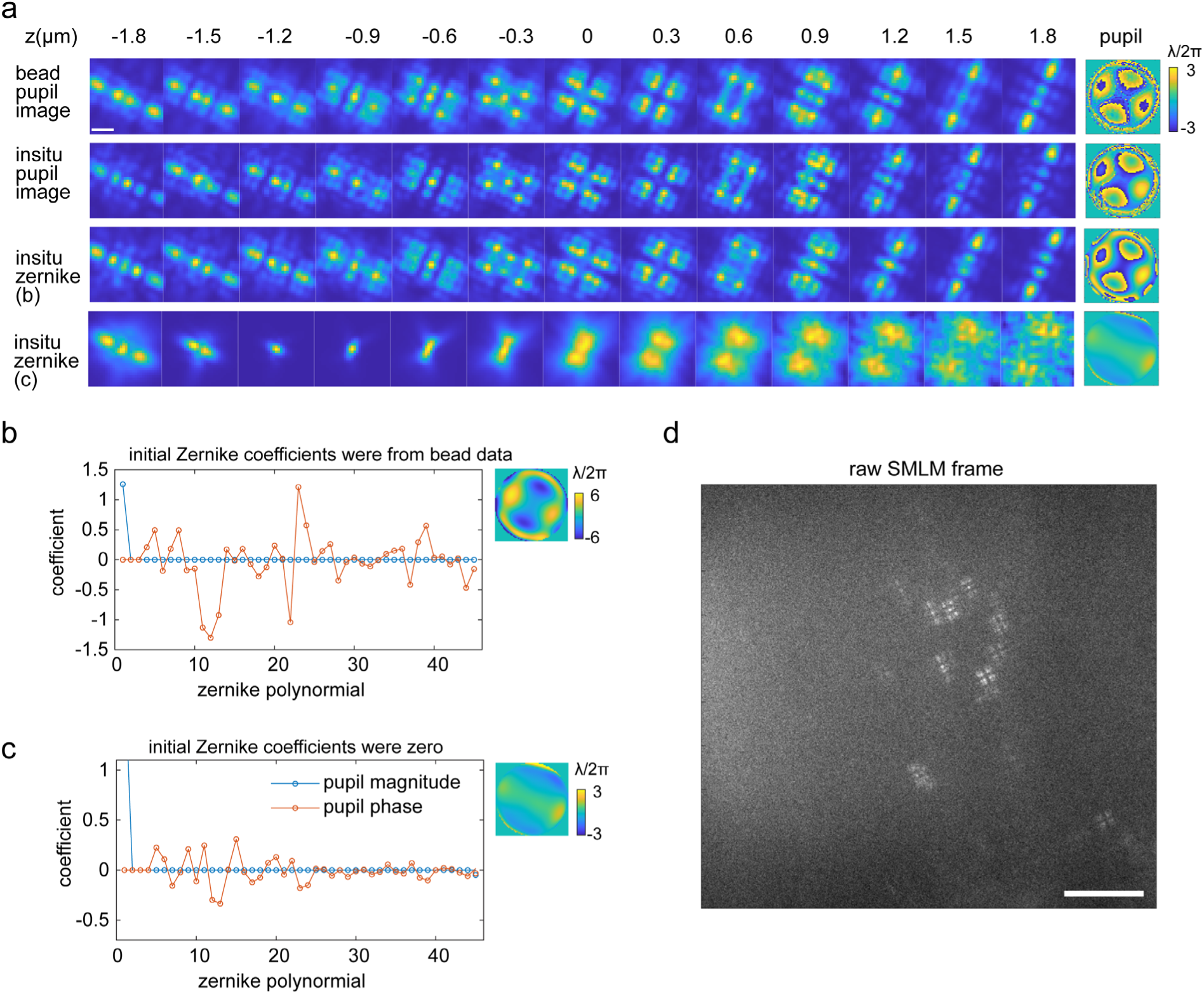
Comparison of pupil-image and Zernike-based *in situ* PSF estimation of Tetrapod blinking patterns generated from a phase plate. (a) Comparison of estimated PSF and pupil from bead data and single molecule blinking patterns. The phase patten generated from a phase plate is more complex than the ones from a deformable mirror, both the pupil-imaged based and Zernike-based modelling methods can retrieve the pupil function from single molecule blinking data. However, the Zernike-based method requires accurate initial values of the Zernike coefficients, otherwise, it fails to retrieve the accurate pupil. (b,c) Estimated Zernike coefficients from Zernike-based *in situ* PSF modelling with (b) or without (c) accurate initial values of the Zernike coefficients. (d) Example raw camera frame used for *in situ* PSF learning. Scale bar, 1 µm (a), 10 µm (d).

**ED Fig. 8.**
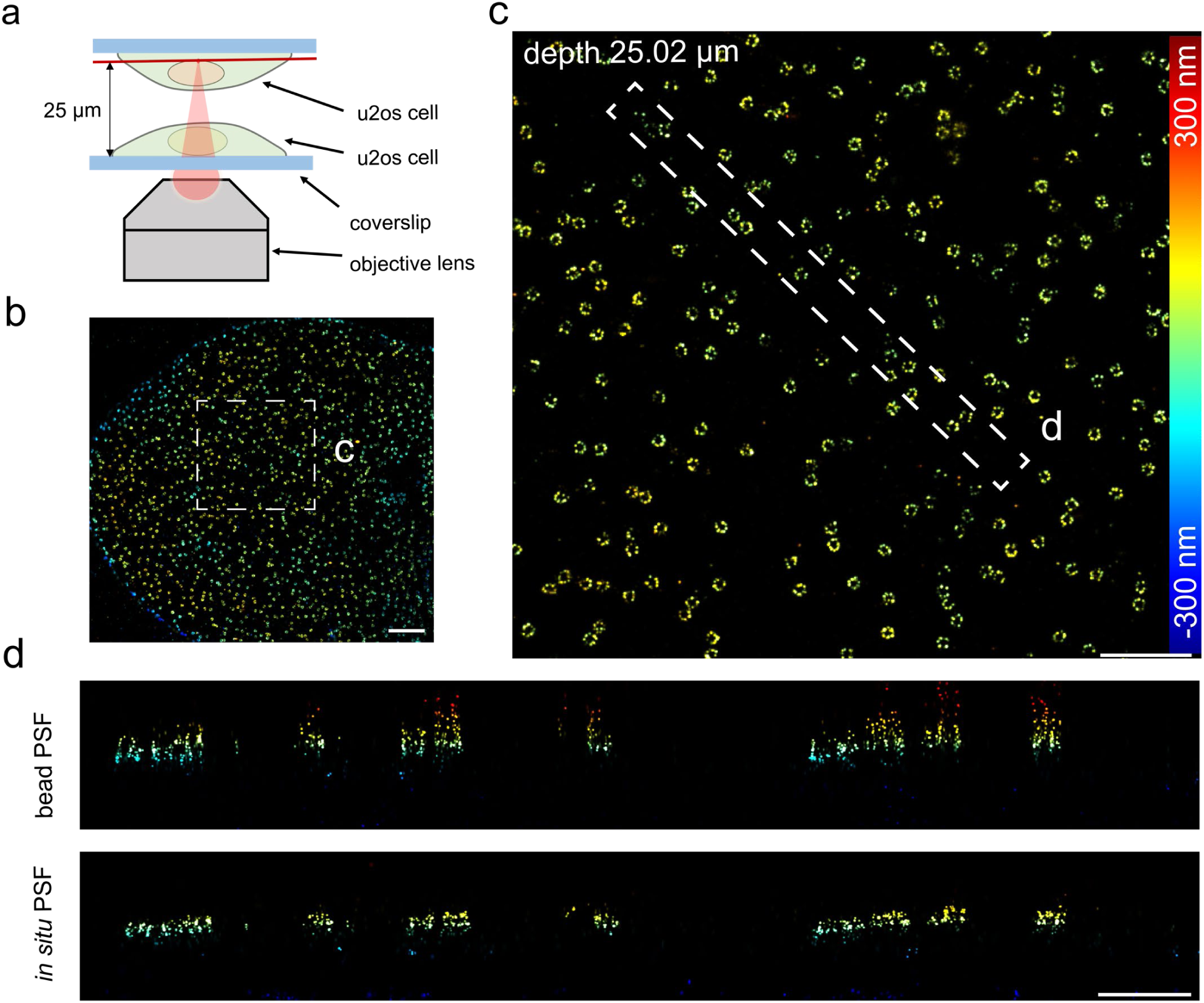
Validation of *in situ* PSF modelling for imaging a thick sample. Imaging of Nup96-SNAP-AF647 in U2OS cells was performed using an oil immersion objective lens (NA=1.5) on a single-channel SMLM system equipped with a DM. (a) A sandwiched sample of U2OS cells was prepared for an imaging depth of 25 µm. (b) Top view of the reconstructed Nup96 using the *in situ* PSF model. (c) Enlarged XY view of the selected region in (b). (d) XZ view of the selected region in (c) reconstructed from bead and *in situ* PSF models, respectively. Scale bars, 2 µm (b), 1 µm (c), and 0.5µm (d).

**ED Fig. 9.**
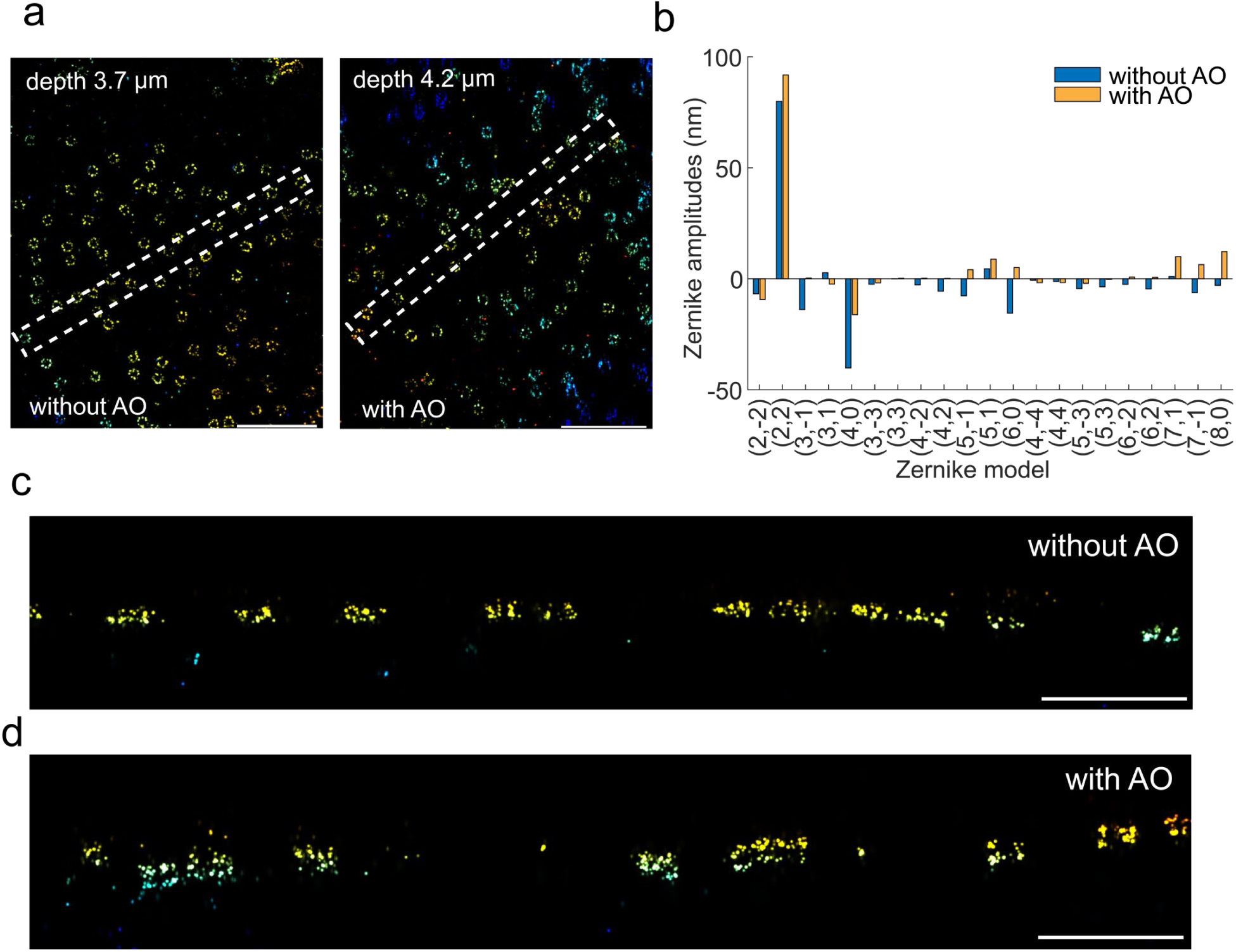
Application of uiPSF for aberration correction using a deformable mirror. (a) Top view of the reconstructed Nup96 using the *in situ* PSF model with and without uiPSF guided aberration correction by AO. (b) Zernike aberrations calculated from single molecule data using uiPSF with and without AO correction. (c-d) XZ view of the selected region in (a) for samples with and without AO correction. After uiPSF guided aberration correction, the reconstructed image quality improved significantly as demonstrated by the resolved double ring structure. Scale bars, 1 µm (a), 500 nm (c-d).

## Notes

https://github.com/ries-lab/uiPSF

### Competing Interest Statement

The authors have declared no competing interest.

