## Supplementary File for "Universal inverse modelling of point spread functions for SMLM localization and microscope characterization"

### Supplementary Figures

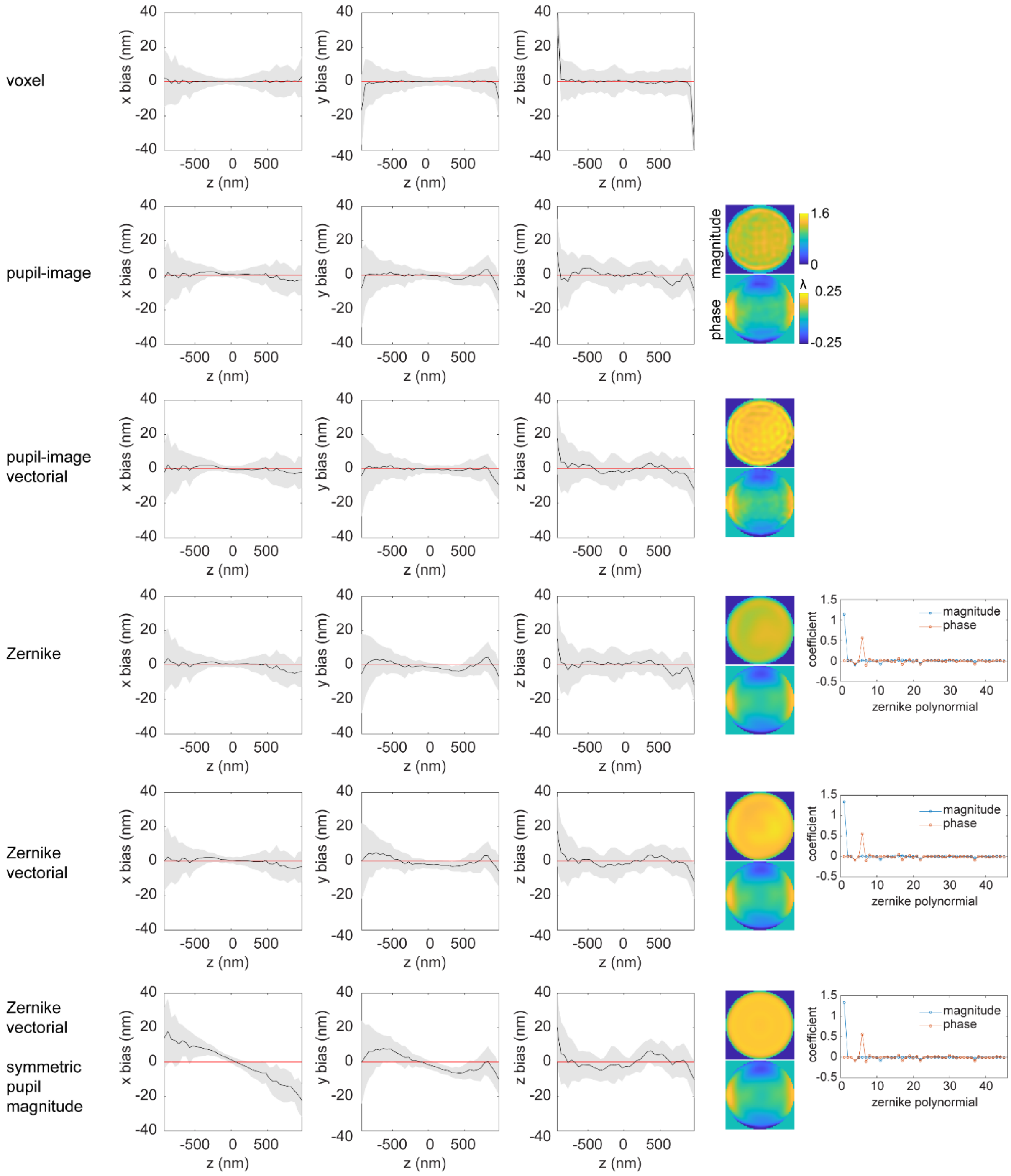

**SI Fig 1. Comparison of different PSF modelling methods** for estimating the PSF model of a single-channel system from experimental bead data. Bead data were collected by imaging 40 nm red bead at z positions from -1000 nm to 1000 nm, with a step size of 50 nm. Localization is performed on the bead data used for inverse modelling. In the last row, the pupil magnitude was modelled only by Zernike polynomials that are circular symmetric.

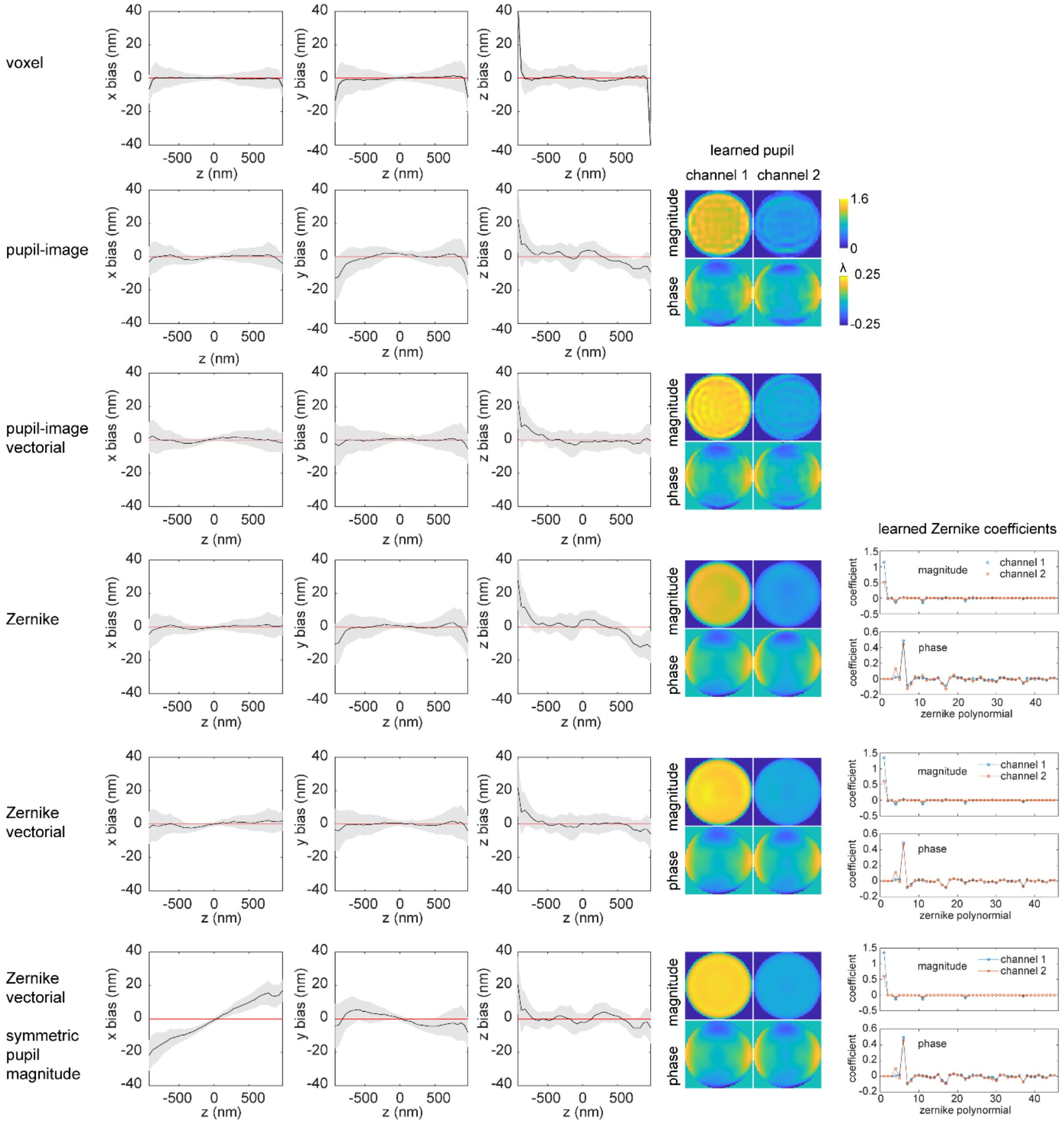

**SI Fig 2. Comparison of different PSF modelling methods for estimating the PSF model of a ratiometric dual-color system** from experimental bead data. Bead data were collected by imaging 40 nm red bead at z positions from -1000 nm to 1000 nm, with a step size of 50 nm. Localization is performed on the bead data used for inverse modelling. In the last row, the pupil magnitude was modelled only by Zernike polynomials that are circular symmetric.

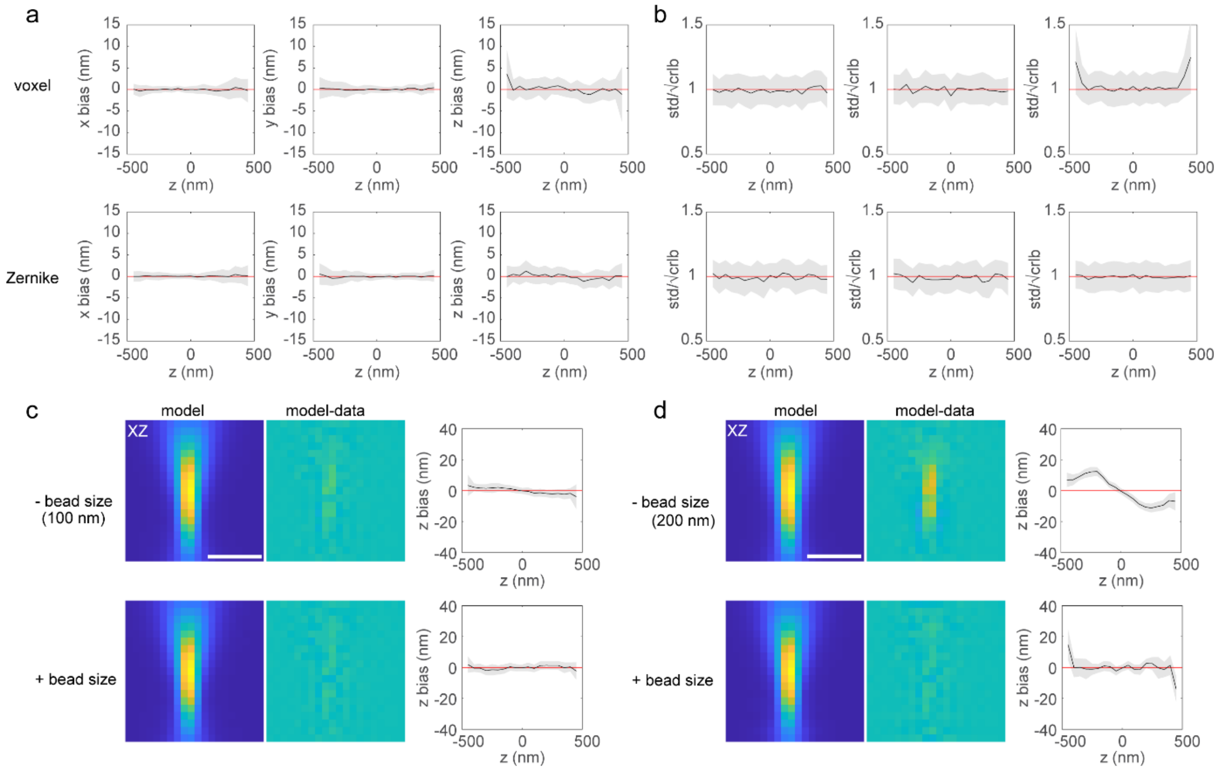

**SI Fig 3. Localization test of estimated voxel and Zernike based PSF model on simulated data of a single-channel system with astigmatism aberration.** (a) Localization of simulated data which were not used during inverse modelling process. The bead size used in the simulation is 50 nm. The data were simulated at z positions from -500 nm to 500 nm, with a step size of 50 nm and 40 images per z position. (b) Ratio of the standard deviation of the localized positions and the average theoretical estimation precision ($\sqrt{CRLB}$) over 40 repeats per z position. A value ~1 indicates that we reach the CRLB. (c) Comparison of estimated PSF models with (+) or without (-) incorporating bead size in the forward model and their axial localization biases. PSF models were estimated from simulated data with a bead size of 100 nm. Localization was performed on simulated data with a bead size of 40 nm. The residue shows the difference between the PSF model and the simulated data with a bead size of 40 nm. (d) Same as c, except that the PSF models were estimated from simulated data with a bead size of 200 nm. Scale bars: 1 µm.

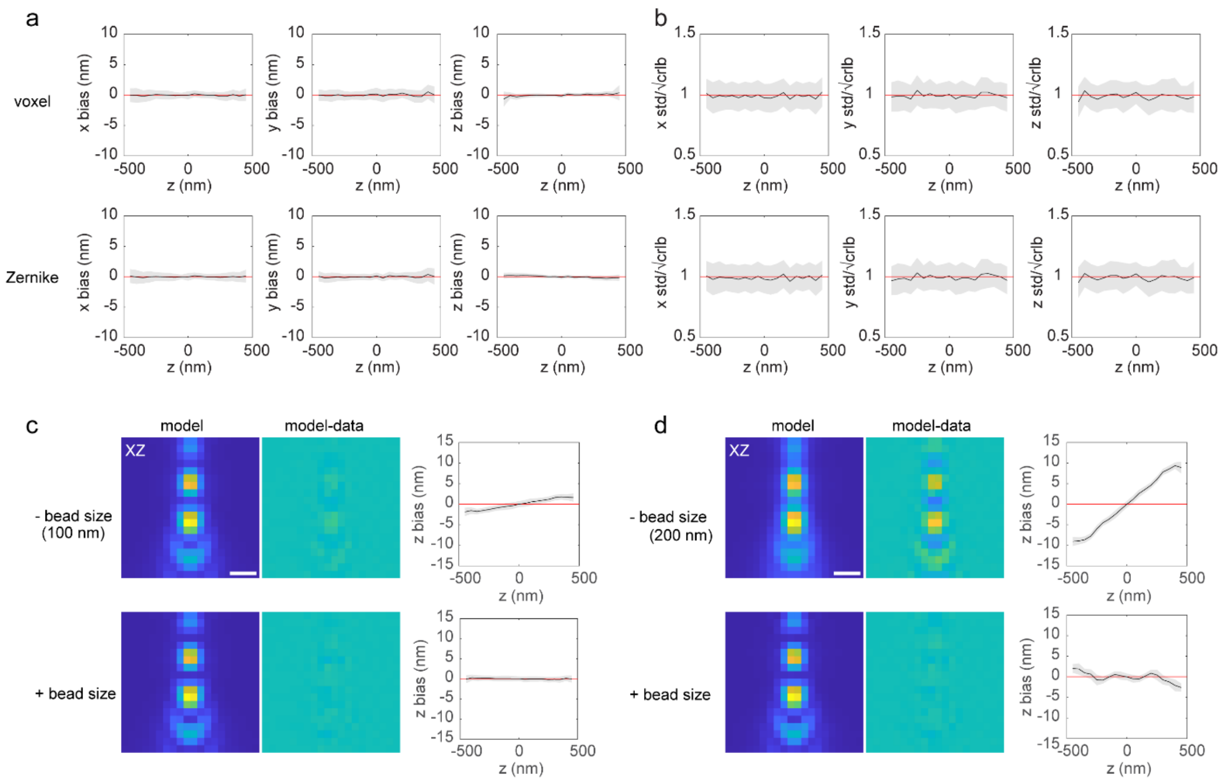

**SI Fig 4. Localization test of estimated voxel and Zernike based PSF model on simulated data of a 4Pi-SMLM system**. (a) localization of simulated data which were not used during inverse modelling process. The bead size used in the simulation is 50 nm. The data were simulated at z positions from -500 nm to 500 nm, with a step size of 50 nm and 40 images per z position. (b) Ratio of the standard deviation of the localized positions and the average theoretical estimation precision ($\sqrt{CRLB}$) over 40 repeats per z position. (c) Comparison of estimated PSF models with (+) or without (-) incorporating bead size in the forward model and their axial localization biases. PSF models were estimated from simulated data with a bead size of 100 nm. Localization was performed on simulated data with a bead size of 40 nm. The residue shows the difference between the PSF model and the simulated data with a bead size of 40 nm. (d) Same as c, except that the PSF models were estimated from simulated data with a bead size of 200 nm. Scale bars: 500 nm.

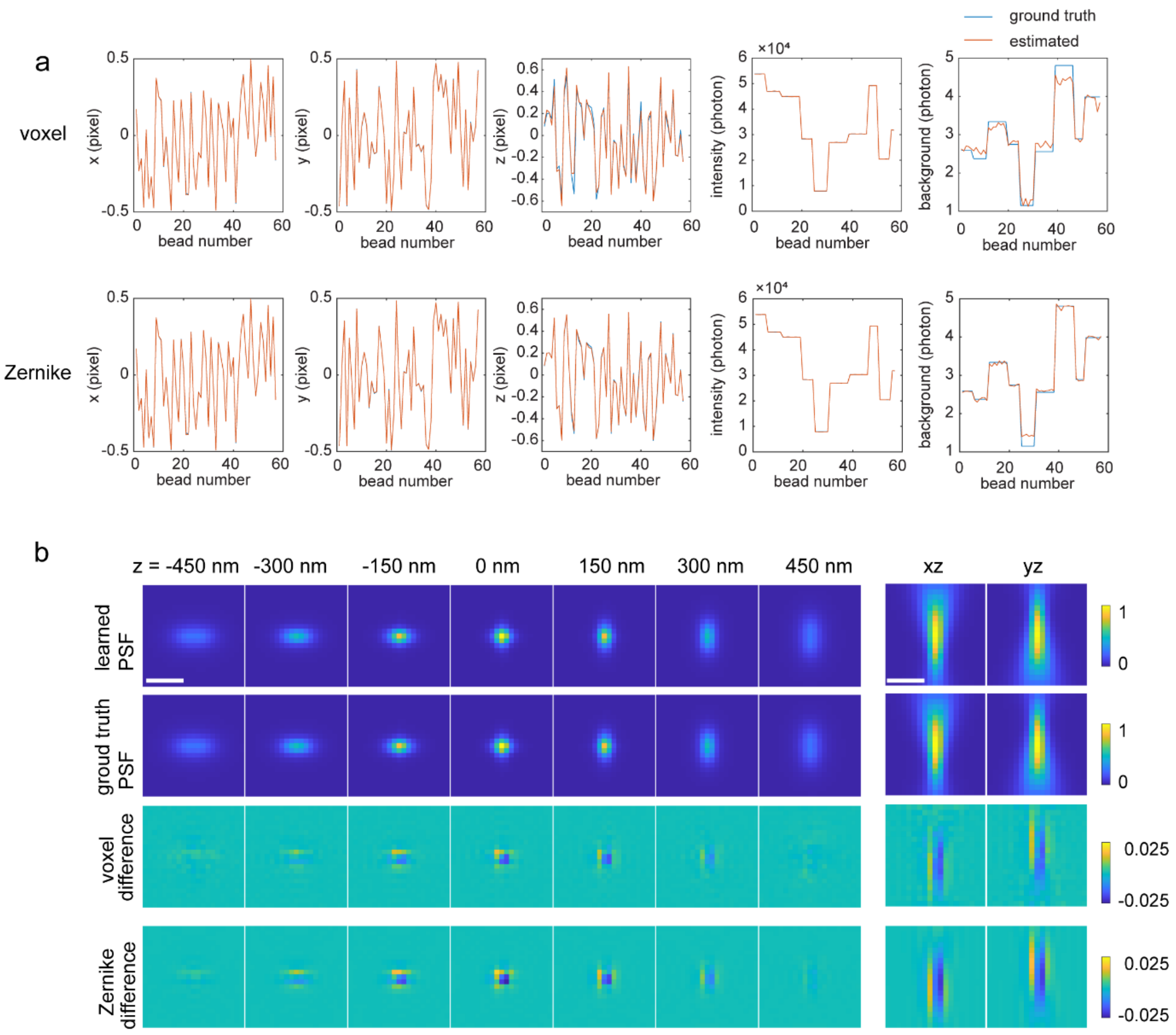

**SI Fig 5. Estimation accuracy of voxel and Zernike based PSF model on simulated data of a single-channel system with astigmatism aberration.** (a) Comparison of the ground truth and estimated values of the x, y, z, photon and background of each bead stack. (b) Comparison of the ground truth and the estimated PSF models. Scale bars: 1 µm.

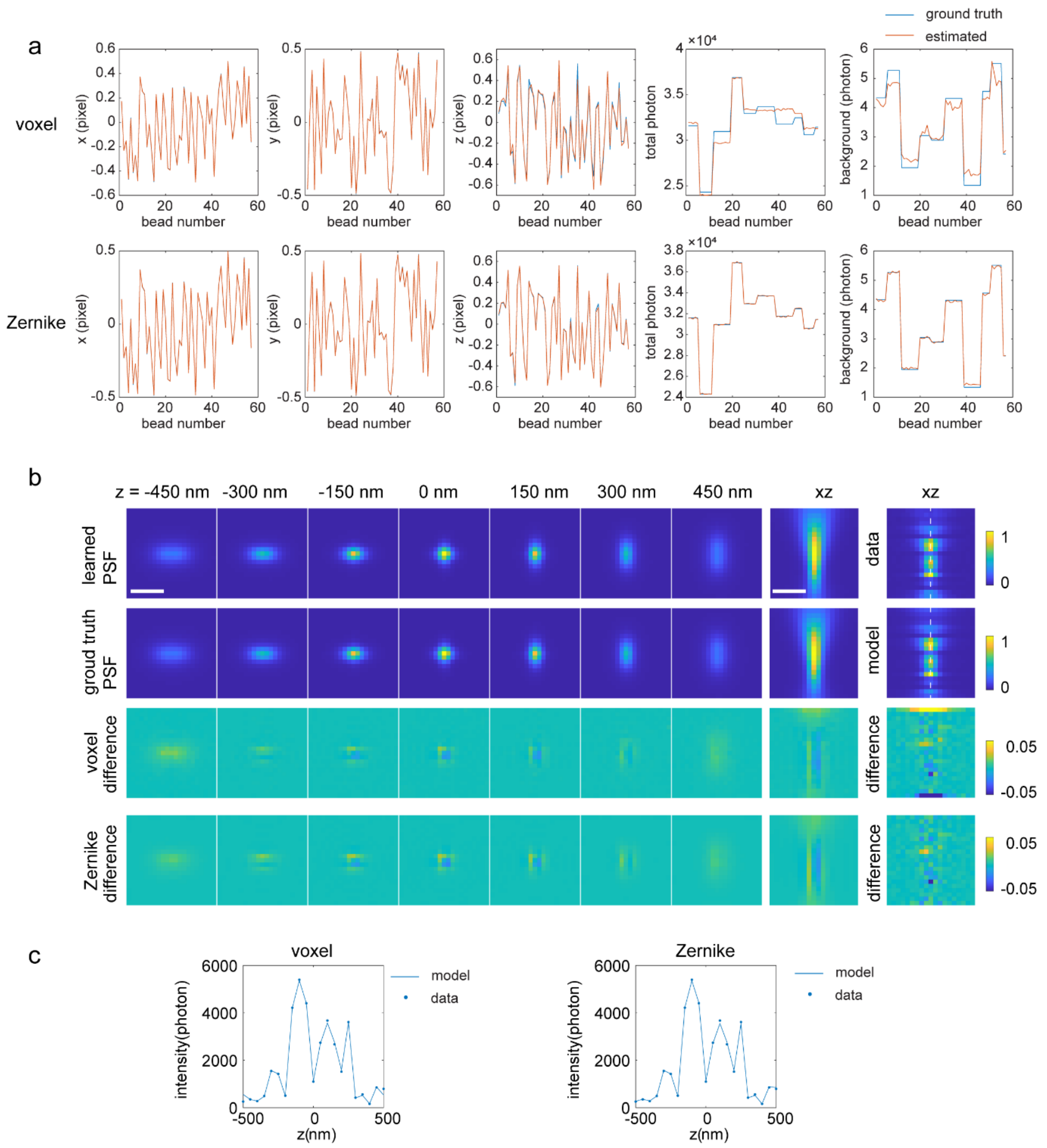

**SI Fig 6. Estimation accuracy of voxel and Zernike based PSF model on simulated data with large photon fluctuation in z.** (a) Comparison of the ground truth and estimated values of the x, y, z, average photon over z and background of each bead stack. (b) Comparison of the ground truth and the estimated PSF models. The last column shows an example comparison of one bead data and its corresponding forward model. (c) Example comparison of the intensity at the center pixel over z between the data and its forward model. Scale bars: 1 µm.

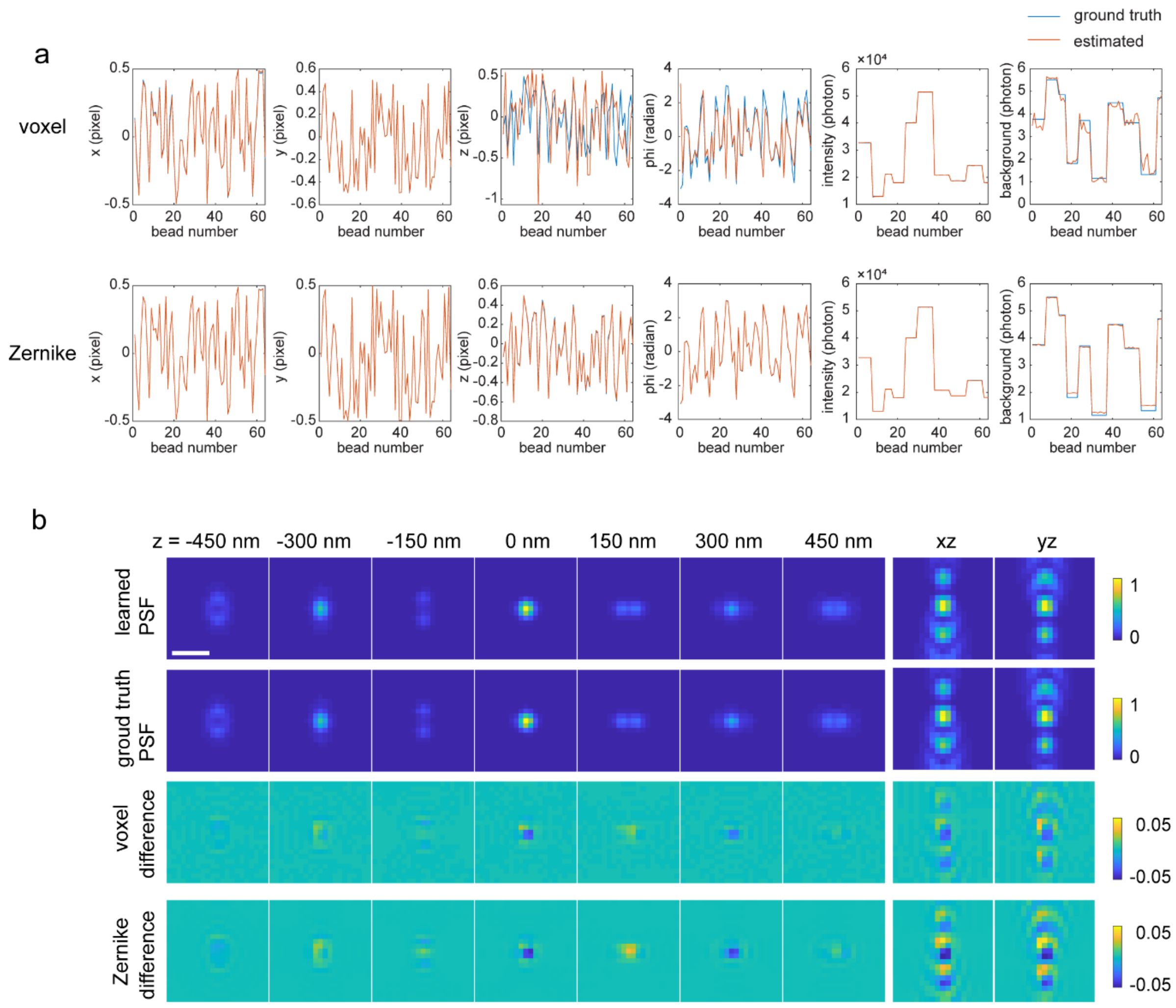

**SI Fig 7. Estimation accuracy of voxel and Zernike based PSF model on simulated data of a 4Pi-SMLM system.** (a) Comparison of the ground truth and estimated values of the x, y, z, phase, photon and background of each bead stack. (b) Comparison of the ground truth and the estimated PSF models. Scale bar: 1 µm.

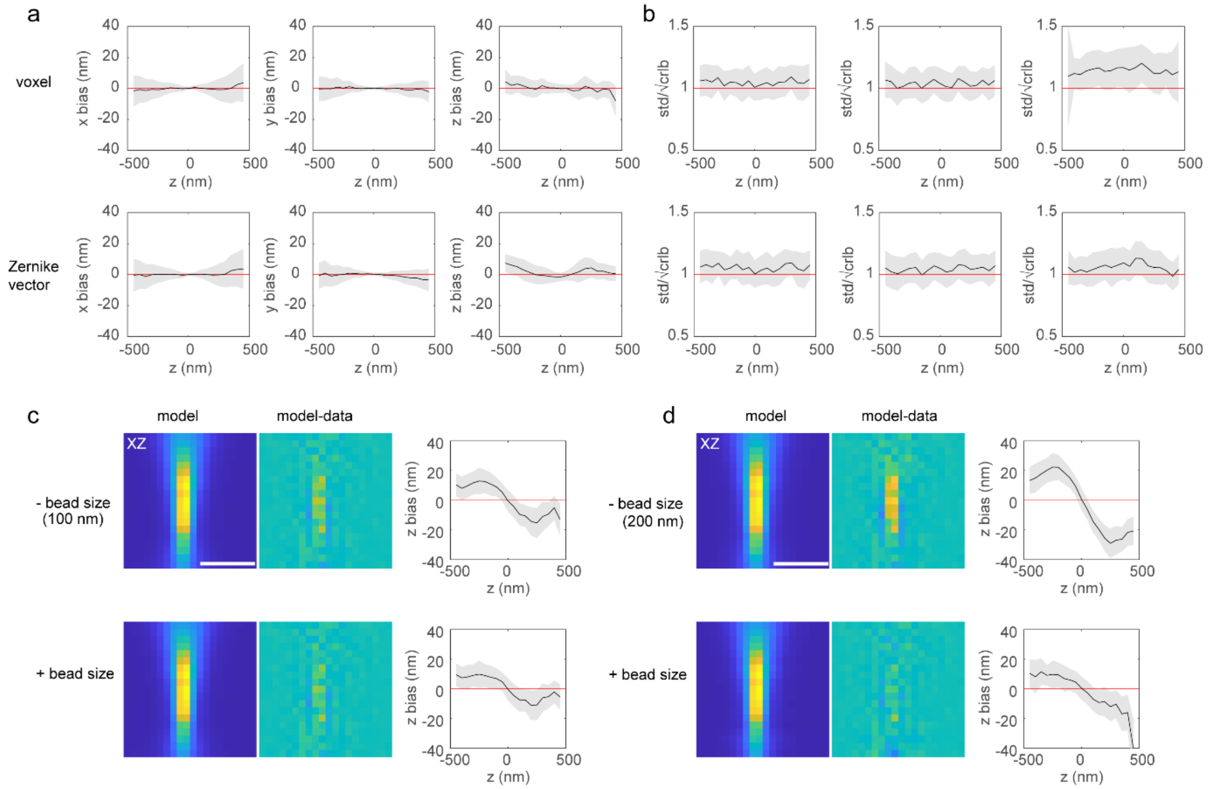

**SI Fig 8. Experimental localization test of estimated voxel and Zernike-vector based PSF model from a single-channel system.** (a) Localization of bead data which were not used during inverse modelling process. The bead size is 40 nm. The data were collected at z positions from -500 nm to 500 nm, with a step size of 50 nm and 40 frames per z position. (b) Ratio of the standard deviation of the localized positions and the average theoretical estimation precision ($\sqrt{CRLB}$) over 40 repeats per z position. (c) Comparison of estimated PSF models with (+) or without (-) incorporating bead size in the forward model and their axial localization biases. Voxel-based PSF models were estimated from 100 nm bead data. Localization was performed on 40 nm bead data. The residue shows the difference between the PSF model and the 40 nm bead data. (d) same as c, except that the PSF models were estimated from 200 nm bead data. Scale bars: 1 µm.

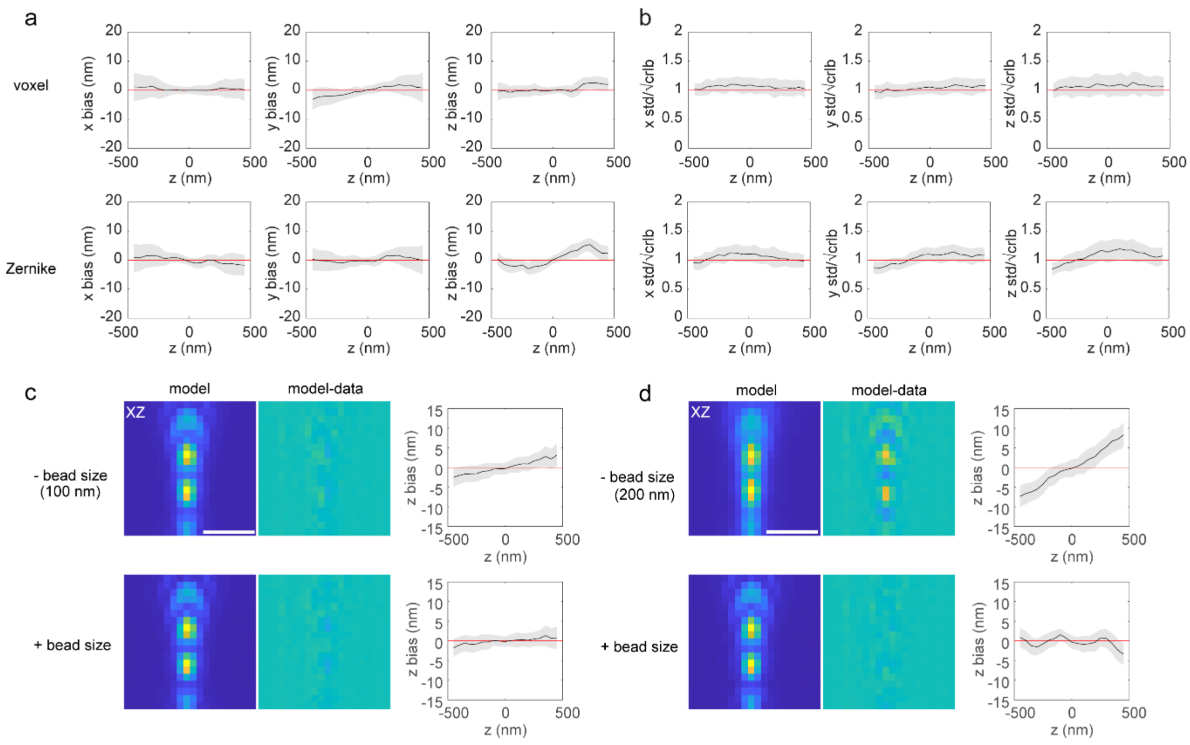

**SI Fig 9. Experimental localization test of estimated voxel and Zernike based PSF model from a 4Pi-SMLM.** (a) Localization of bead data which were not used during inverse modelling process. The bead size is 40 nm. The data were collected at z positions from -500 nm to 500 nm, with a step size of 50 nm and 40 frames per z position. (b) Ratio of the standard deviation of the localized positions and the average theoretical estimation precision ( $\sqrt{\text{CRLB}}$) over 40 repeats per z position. (c) Comparison of estimated PSF models with (+) or without (-) incorporating bead size in the forward model and their axial localization biases. Voxel-based PSF models were estimated from 100 nm bead data. Localization was performed on 40 nm bead data. The residue shows the difference between the PSF model and the 40 nm bead data. (d) Same as c, except that the PSF models were estimated from 200 nm bead data. Scale bar: 1 µm.

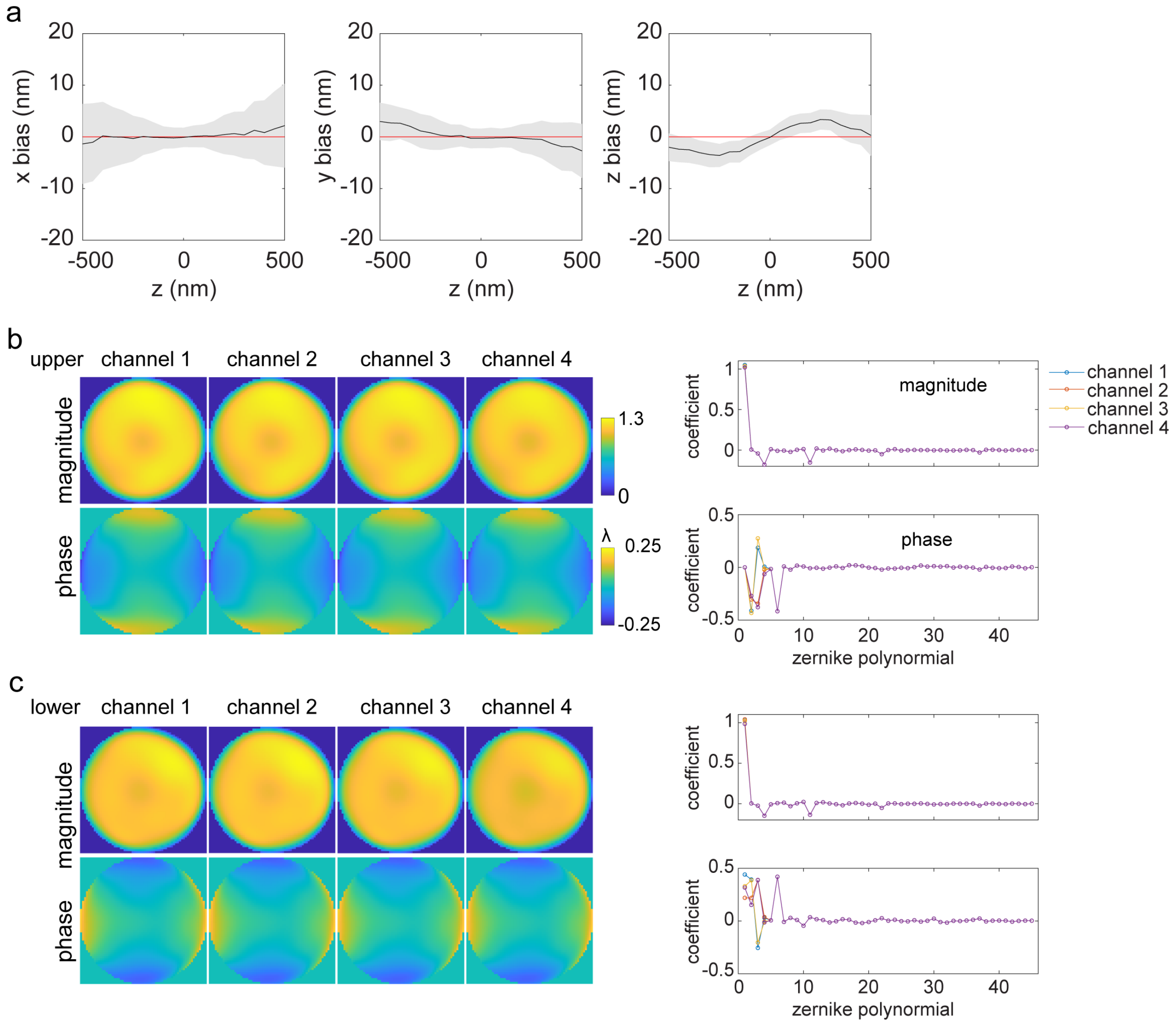

**SI Fig 10. Estimation of Zernike-based PSF model of a 4Pi-SMLM system from experimental bead data**. Bead data were collected by imaging 40 nm red bead at z positions from -500 nm to 500 nm, with a step size of 50 nm and three phase positions at -2π/3, 0, 2π/3 were collected at each z position. (a) Localization on bead data which were used for inverse modelling. Here the z position is converted from estimation of the phase. (b) Estimated Zernike coefficients of each channel from the upper emission path and the corresponding pupil function. (c) Estimated Zernike coefficients of each channel from the lower emission path and the corresponding pupil function.

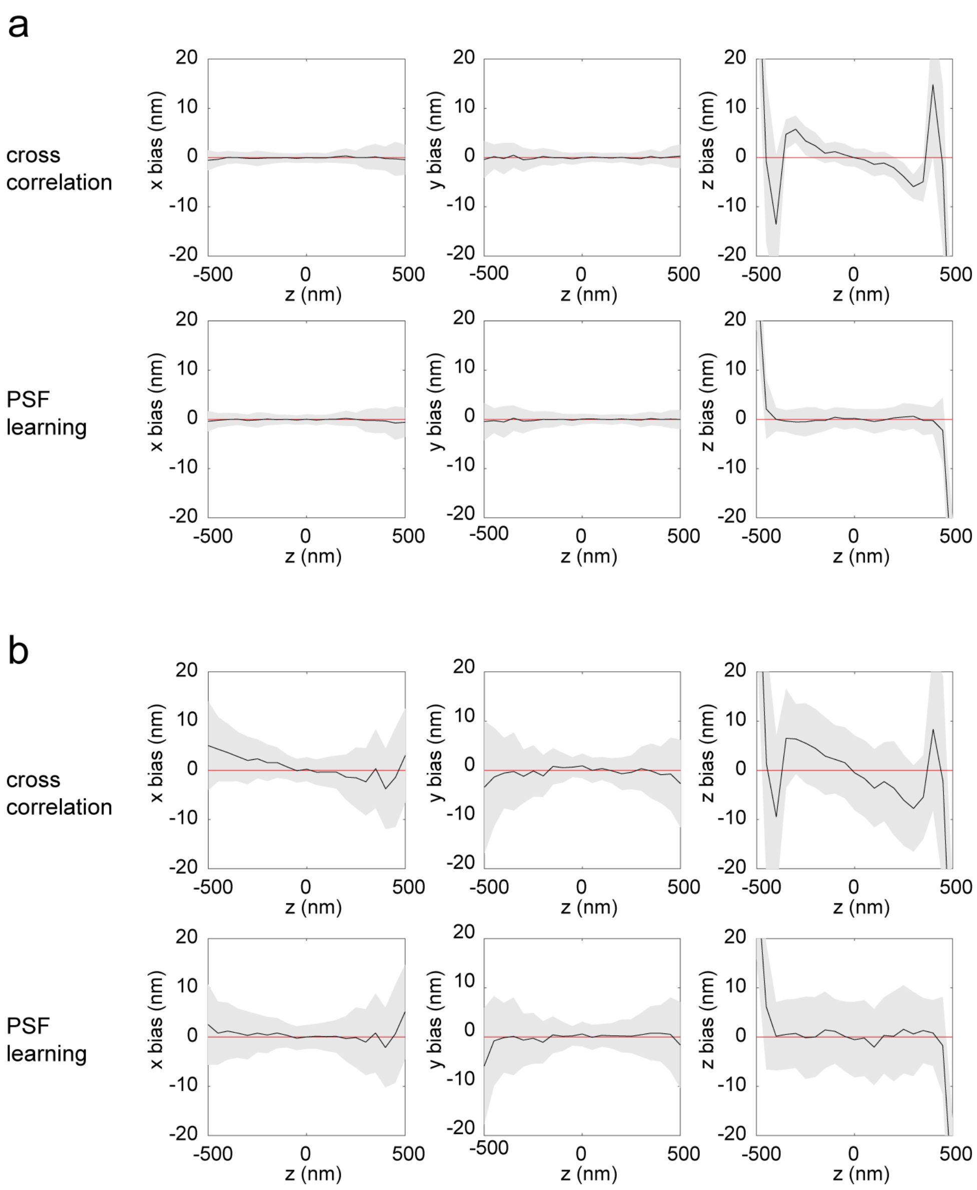

**SI Fig 11. Comparison of voxel-based PSF modeling with uiPSF and traditional cross-correlation based method.** (a) Test on simulated data. The data were simulated at z positions from -500 nm to 500 nm, with a step size of 50 nm. An astigmatism aberration of 0.5 and a bead size of 50 nm were used in the simulation (b) Test on experimental data. 40 nm bead data were collected at z positions from -500 nm to 500 nm, with a step size of 50 nm. The cross-correlation based method was described previously in Li *et al.*[1] and its implementation in SMAP was used.

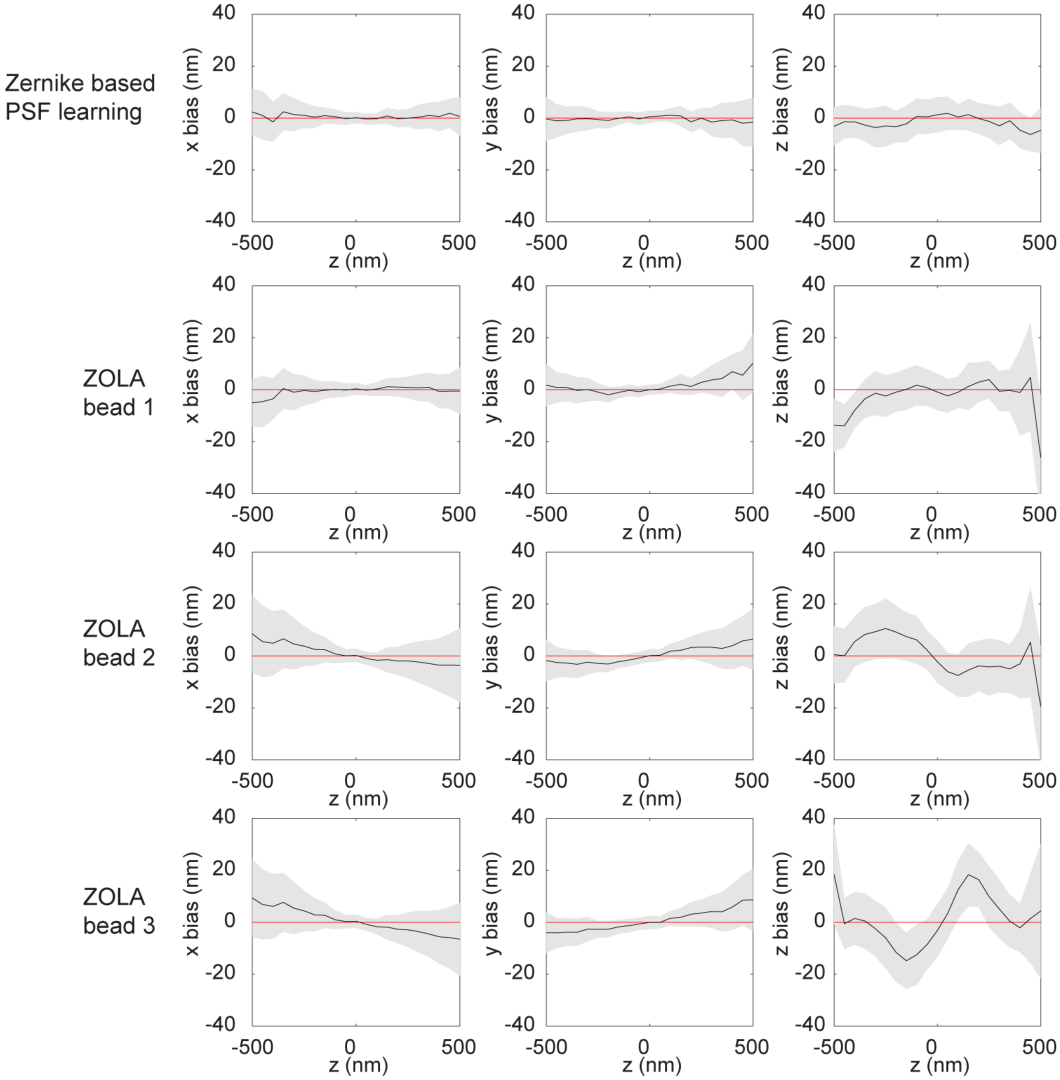

**SI Fig 12. Comparison of Zernike-based PSF modelling using uiPSF and Zola 3D.** 40 nm bead data were collected at z positions from -500 nm to 500 nm, with a step size of 50 nm. Here we show the localization performance on the bead data used for PSF modelling (56 bead data). For uiPSF, the vectorial PSF modelling method was used and the PSF model was estimated from 82 bead stacks. For Zola 3D, the imageJ plugin was used and the PSF model was estimated from a single bead stack (which performs better than using multiple beads). Here we tested three PSF models generated from three different beads. It shows that the localization bias using Zola 3D is bead dependent.

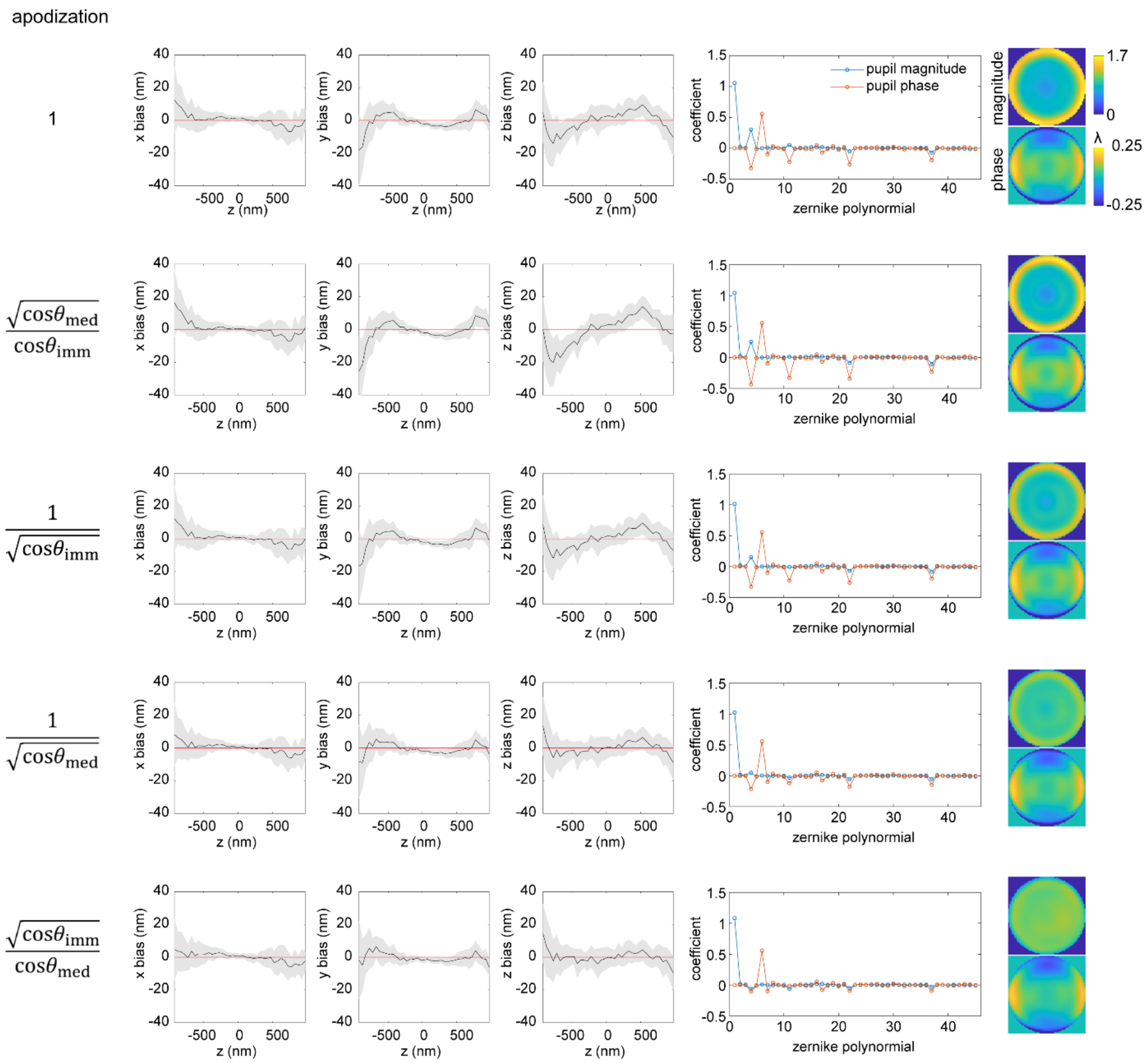

**SI Fig 13. Comparison of different apodization terms at the presence of index mismatch aberration.** The Zernike vectorial PSF model was used. Bead data were collected by imaging 40 nm red bead at z positions from -1000 nm to 1000 nm, with a step size of 50 nm. Localization was performed on the bead data used for inverse modelling. All tested apodization terms are identical if refractive indices are matched.

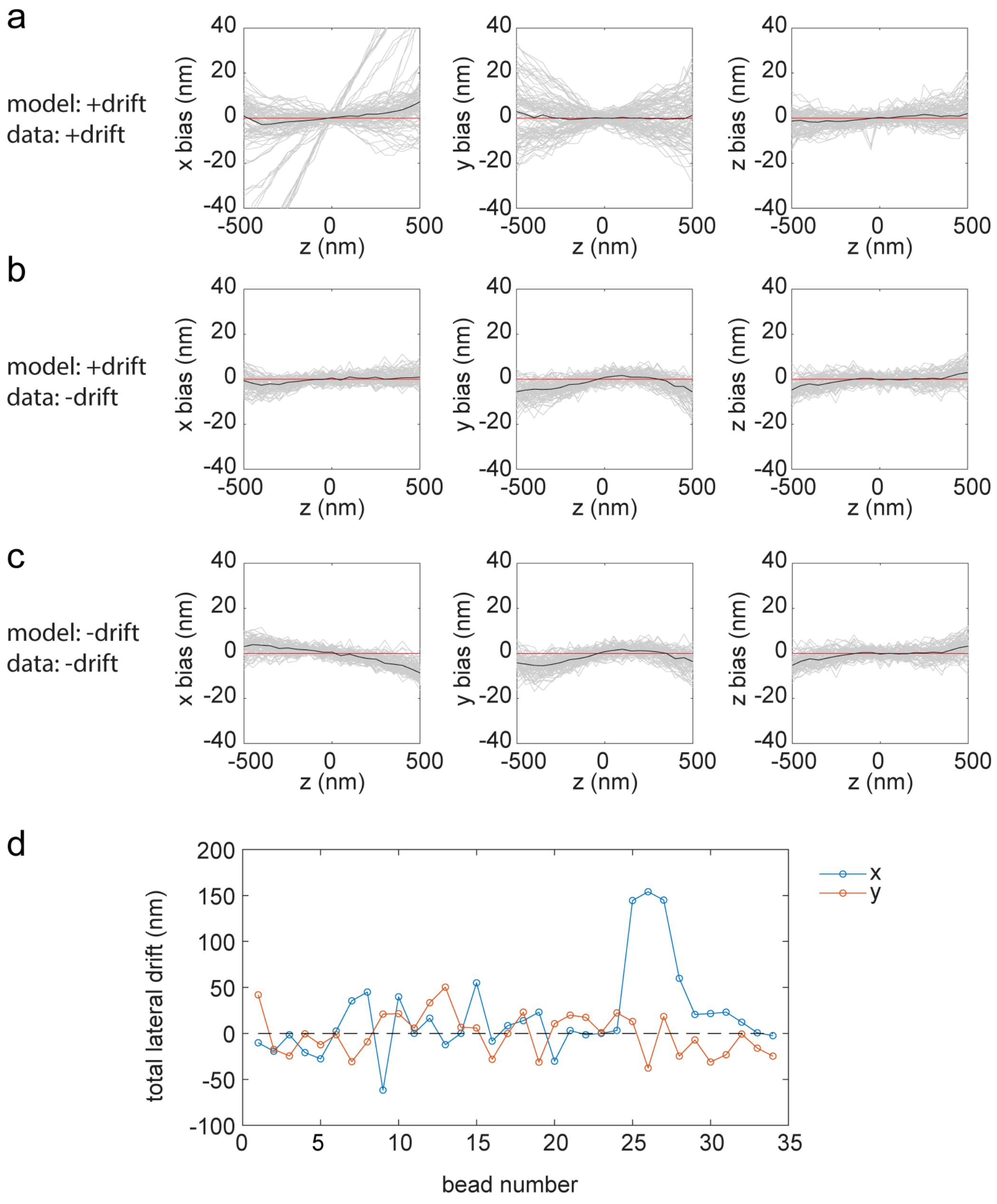

**SI Fig 14. Effect of considering lateral drift in the forward model.** Data were collected from a 4Pi-SMLM system. Voxel-based PSF modelling method was used. (a) Localization of the data used for inverse modelling. The data contains bead stacks with large drift in x. The inverse modelling includes the estimation of the lateral drift. (b) Localization of a different dataset that contains small lateral drifts. The inverse modelling included the estimation of the lateral drifts, but from the data in a. (c) Localization of the same data in b. The inverse modelling assumed no lateral drifts and used the data in a. The localization bias in x is larger than the one in b, which indicates that without estimating the lateral drifts during inverse modelling, the estimated model will be affected by the lateral drifts in the data. (d) The estimated lateral drifts from inverse modelling using the data in a.

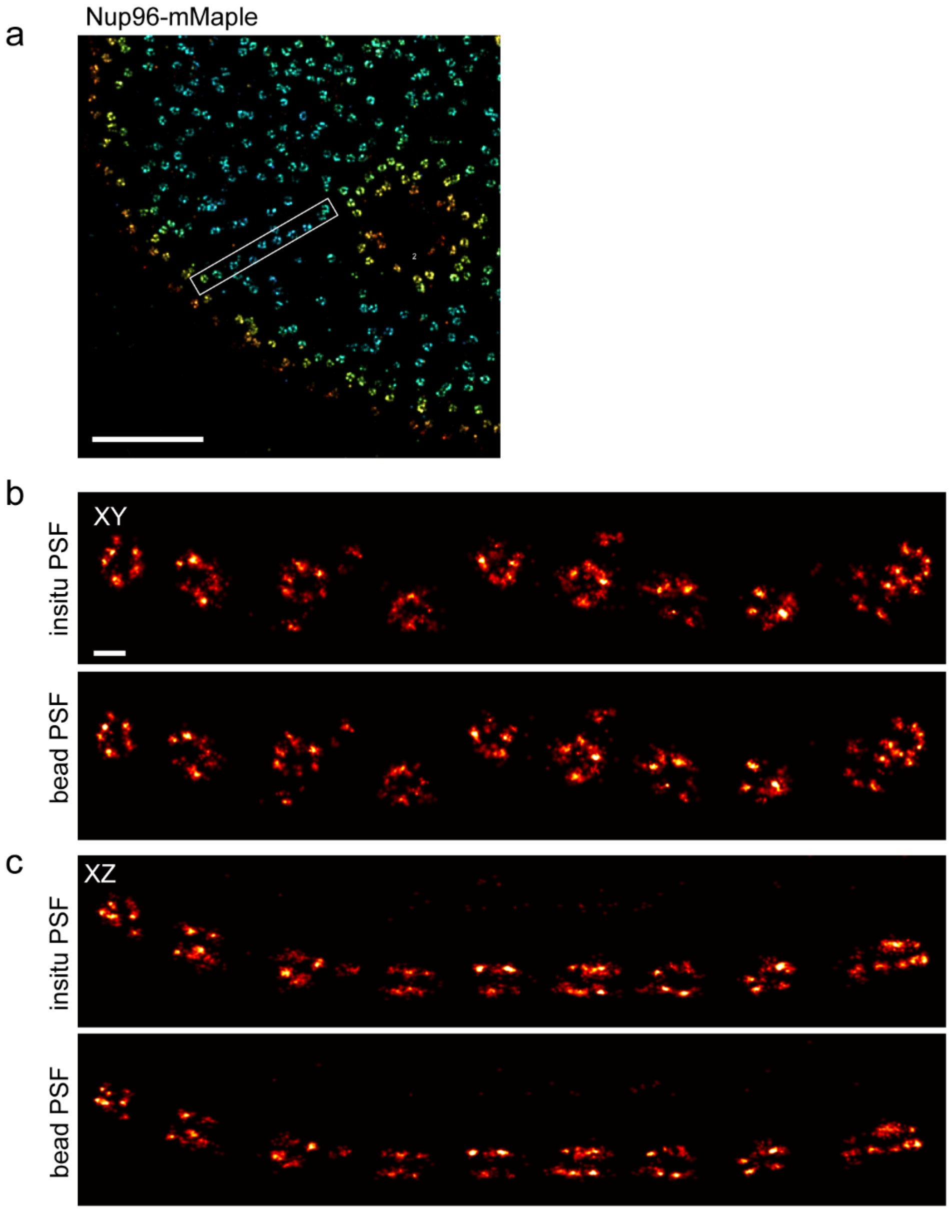

**SI Fig 15. Reconstruction of Nup96-mMaple in U2OS cells from 4Pi-SMLM using the PSF models estimated from bead data and single-molecule blinking data.** (a) A subregion of the reconstructed Nup96 using the bead PSF model. Same as Fig .2h. (b) XY view of the selected region in a from bead and *in situ* PSF models. (c) XZ view of the selected region in a from bead and *in situ* PSF models. Here, the refractive index of sample medium is matched with the silicone oil immersion medium. For *in situ* model, the PSF is directly estimated from the single-molecule blinking data without the need for the additional PSF calibration. Scale bar: 2 µm (a), 100 nm (b,c).

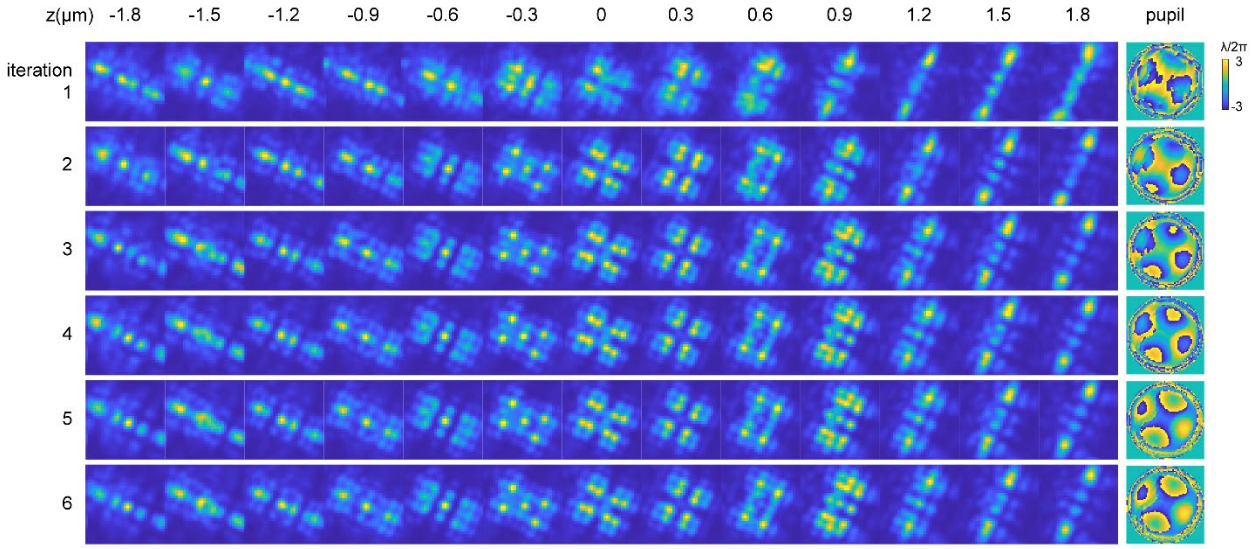

**SI Fig. 16. Pupil-image based *in situ* PSF estimation of Tetrapod blinking patterns generated from a phase plate.** Estimated PSF models and pupil images from iteration 1 to 6 are shown. PSF model converges after four iterations. The initial pupil function was generated from the 13$^{th}$ Zernike polynomial (Noll order, diagonal 2$^{nd}$ astigmatism) with an amplitude of -2. The PSF model and pupil from the 6$^{th}$ iteration are shown in Fig. 3c.

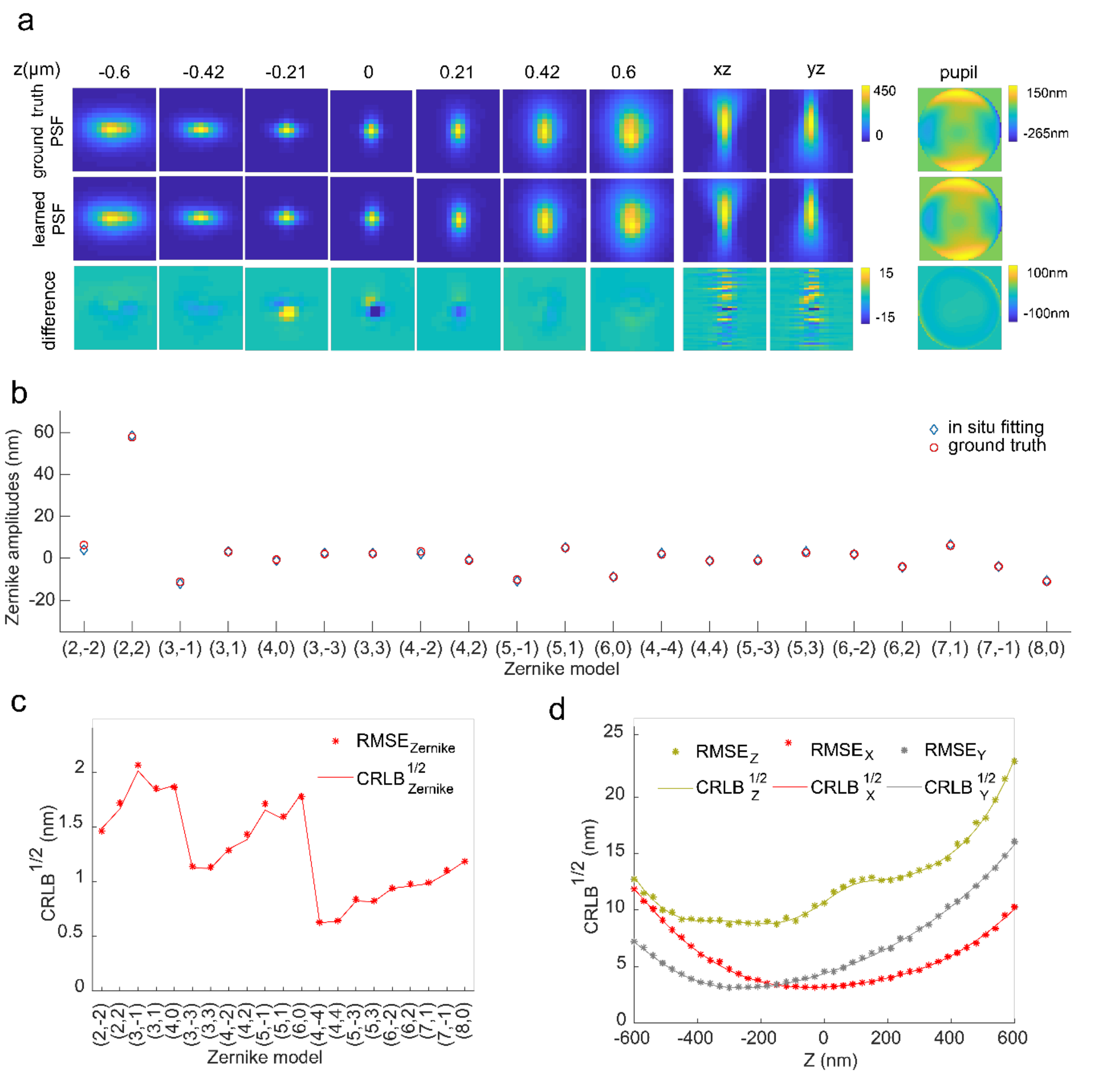

**SI Fig. 17. Performance of *in situ* PSF modeling estimated from single-molecule blinking data**. (a) Comparison of the ground truth and the estimated PSF models from simulated single molecules located in the axial range from -600 nm to +600 nm. The simulated data consist of 41 single molecules equally located at z positions between -600 nm to 600 nm with a total of 5000 photons and a background level of 10. (b) Comparison of the amplitudes of the 21 Zernike modes (blue diamonds) returned from *in situ* PSF modeling and the ground truth Zernike coefficients (red circles). (c) and (d) have the same parameters as (a) and (b). 1000 repeated fitting calculations were performed to determine the localization precision. (c) Localization precision of the Zernike aberrations. (d) Localization precision of 3D positions. CRLB is the Cramér-Rao lower bound.

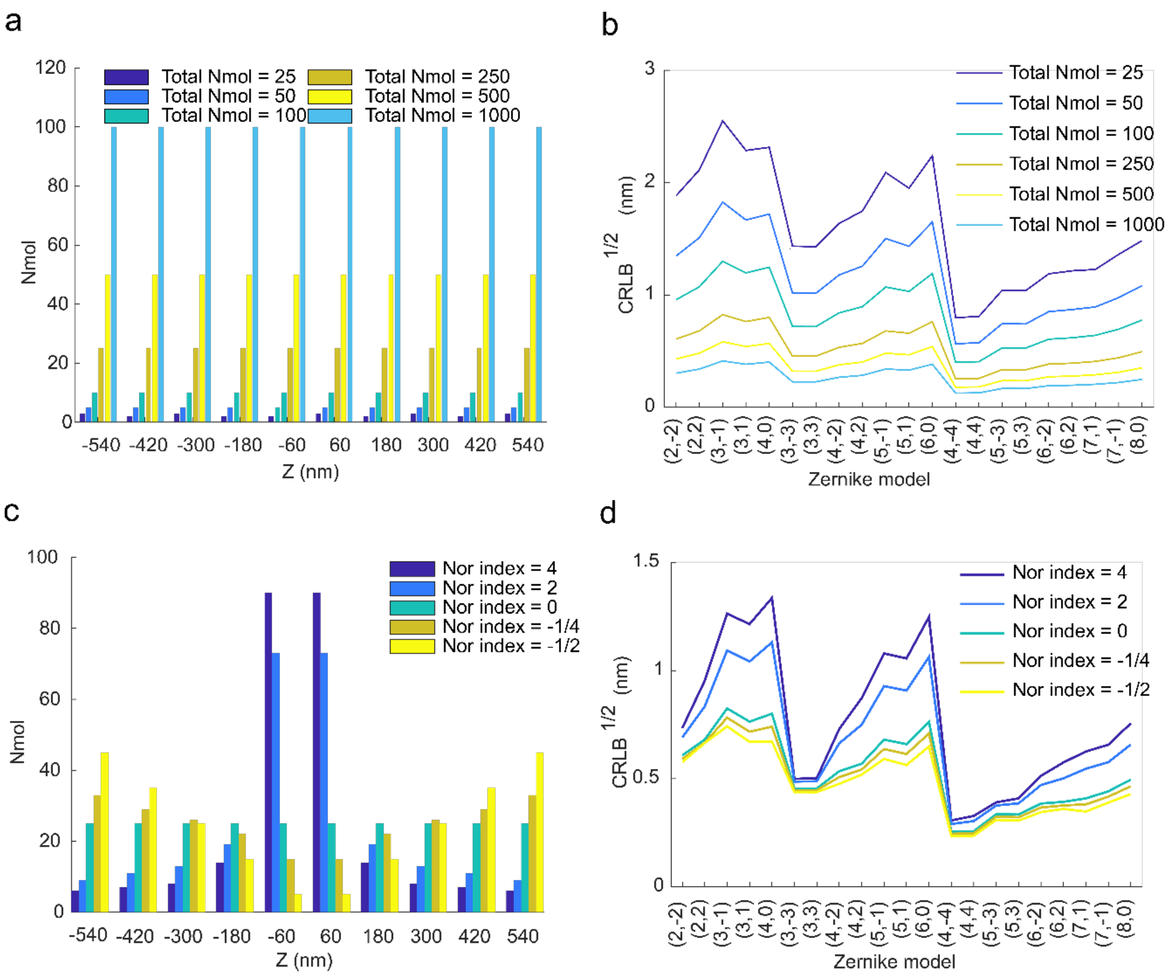

**SI Fig 18. Effect of number and distribution of single molecules along the z-axis on the fitting accuracy.** (a) Uniform distribution of single molecules with varying total numbers along the z-axis. (b) Comparison of fitting accuracy of Zernike coefficients under the distribution described in (a). Total Nmol is the total number of single molecules in fitting. (c) Uneven distribution of 250 single molecules along the z-axis. (d) Comparison of fitting accuracy of Zernike coefficients under the distribution described in (c). Nor index is a measure of the level of aggregation of single molecules near z=0. (a) and (b) have the same simulation parameters as SI Fig. 17. It shows that single molecules with more defocus positions result in better estimation accuracy of the aberrations.

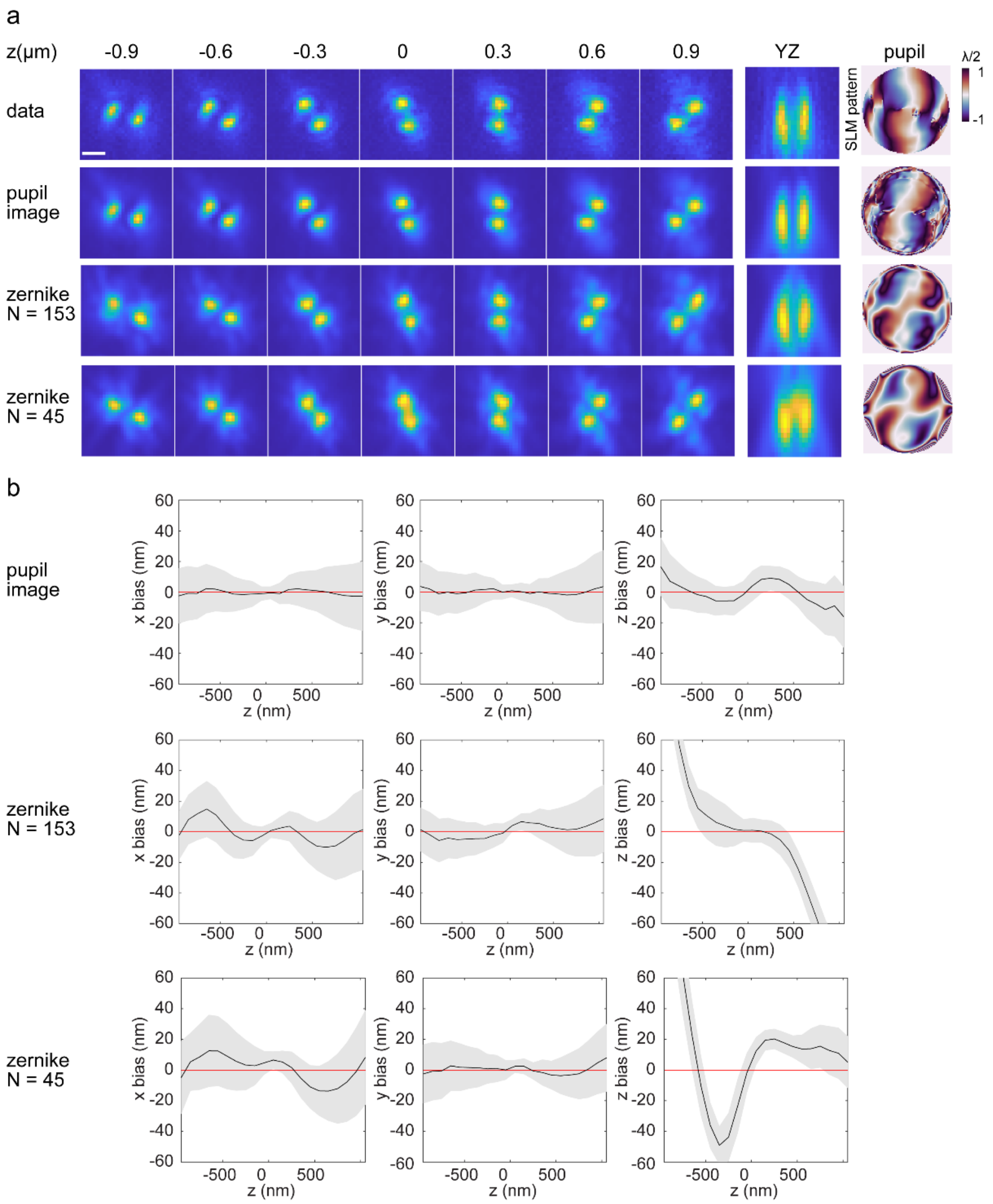

**SI Fig. 19. Comparison of pupil-image and Zernike based learning of double-helix PSFs from bead data.** (a) Comparison of estimated PSF models and the pupil functions with one example bead data and the SLM pattern used to generate the DH-PSFs. The estimated pupil functions show a slight rotation compared to the SLM pattern, this could be caused by misalignment of the SLM and aberrations in the optical system. N represents the number of Zernike polynomials used in learning. (b) Comparison of localization biases on the bead data using different PSF models in a. Zernike-based PSF models result in large bias, even with 153 Zernike polynomials, Zernike expanded pupil function still fails to model the correct pupil for DH-PSFs. The data were collected by imaging 200 nm crimson bead (F8806, Thermo Fisher) dried on coverslip. The beads were imaged by moving the sample stage from -1 to 1 μm, with a step size of 100 nm. A silicone oil objective (NA=1.35) was used. The DH-PSFs were generated from the phase pattern in a using a spatial light modulator (Meadowlark).

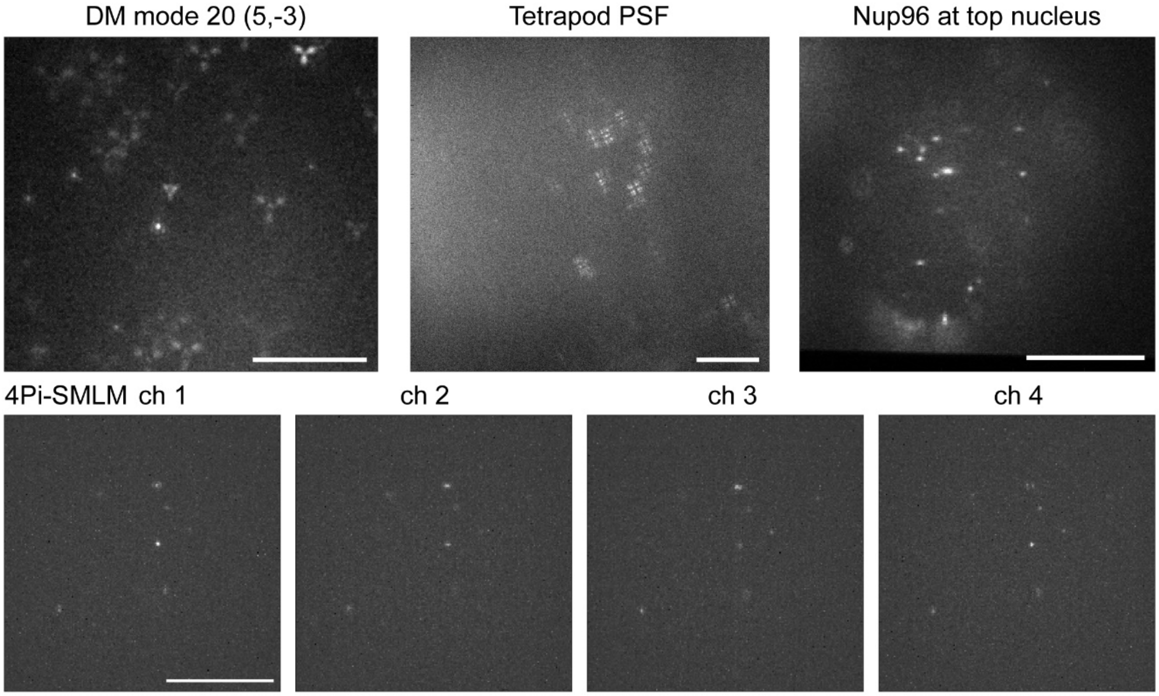

**SI Fig 20. Example SMLM frames from different imaging systems.** Example raw camera frames used for *in situ* PSF modelling of various imaging systems corresponding to the results in Fig. 3. Scale bars, 10 µm.

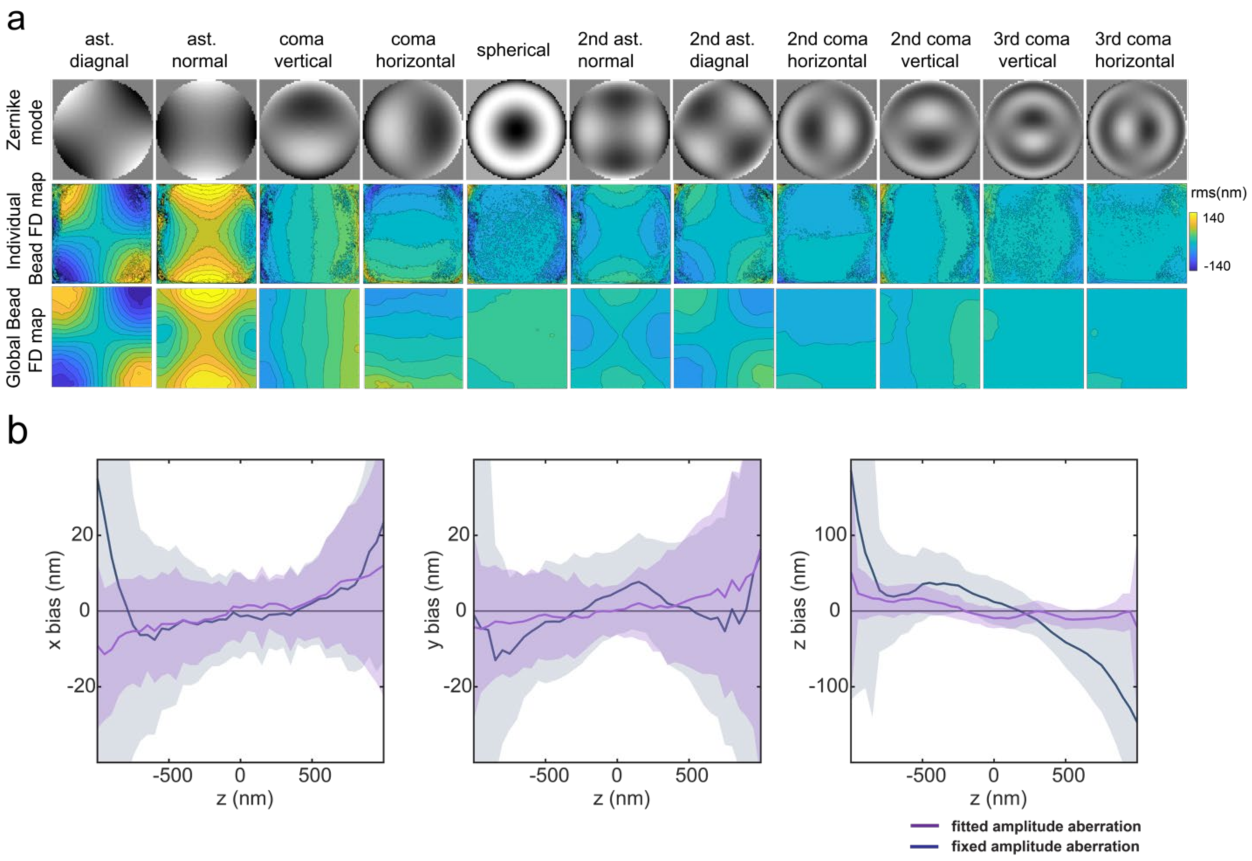

**SI Fig. 21. Comparison of Zernike-based FD PSF modelling using uiPSF and FD-DeepLoc.** (a) FD-DeepLoc individually fits each bead, where each bead corresponds to a set of Zernike coefficients. These coefficients are interpolated to generate an FD map. However, the presence of clustered or unevenly distributed beads during the fitting process often leads to singular values on the FD map. In contrast, uiPSF performs a global fitting on all beads, optimizing the FD map globally and resulting in a relatively smoother FD map. (b) Comparison of the impact of fixing and fitting the amplitude aberration on localization accuracy. The uiFSF obtained PSF models by fitting and fixing the amplitude aberration parameter, and then used them to localize FD bead data. By comparing the results, it was found that relaxing the amplitude aberration parameter resulted in a smaller bias in PSF model localization, indicating that it better represents the true PSF.

### Supplementary Notes

In the supplementary notes we used 'learning' to stand for inverse modelling, not machine learning or deep learning. As in our method, the PSF models are based on physics models. The following conventions for symbols are used: bold for a vector, regular for a scalar or the length of a vector; symbol $i$ in subscript denotes index and in exponential denotes a complex value.

#### 1. Data preprocessing

Data preprocessing includes conversion of the raw camera pixel value to photon counts, segmentation and negative value correction. The pixel value from the camera output has a unit of analog-to-digital unit (ADU), conversion from ADU to photon count is obtained by $D = (D_{\mathrm{raw}} - o)/\gamma$, where o is a constant offset value that can be estimated from the mean of the pixel values at no photon collection, and $\gamma$ is a gain factor quantified through gain calibration[2]. Here we ignore the pixel dependent offset and gain that might exist for sCMOS cameras[3] and use a single-valued offset and gain for data collected from both sCMOS and EMCCD. We also found that when gain calibration is not accessible, setting $\gamma = 1$ will not affect the inverse modeling results.

##### *1.1 Segmentation for single-channel bead data*

Segmentation is to select and crop the candidate bead images from the raw data. For each data stack, a maximum projection is applied along the z dimension to generate a high SNR image. The resulting image is then smoothed by a 2D Gaussian filter with a standard deviation of e.g. 2 (user defined) pixels. The local maxima within a region of e.g. 3 by 3 (user defined) pixels are found by applying a maximum filter of a kernel size of 3 by 3 pixels to the smoothed image. The local maxima with a peak value higher than a set threshold are selected as the coordinates of the candidate bead images. We used a peak threshold at e.g. 20% (user defined) of the maximum value of the smoothed image.  A bead image stack of given subregion size was cropped around the center of each selected coordinate. We used a subregion size of 20-30 pixels along each dimension.

To prevent estimation of negative background values during PSF learning, we subtracted a negative value from the cropped bead stacks, which is estimated from 0.001 quantile of all the pixel values of all bead stacks.

##### *1.2 Segmentation for multi-channel bead data*

For multi-channel data, the candidate bead coordinates for each channel are first selected independently as described above. Then a coordinate pairing procedure is applied to ensure that the coordinates from all channels associated with the same bead are paired together. The pairing procedure consists of two steps. First, find lateral shift between channels: 1) set channel 1 as the reference channel; 2) calculate the channel shift $\boldsymbol{d}$ from the average of coordinate positions $\boldsymbol{X}$ (a vector a two integers) of the current channel $l$ and the reference channel $l = 1$, $\boldsymbol{d}_l = \mathrm{avg}(\boldsymbol{X}_i) - \mathrm{avg}(\boldsymbol{X}_1)$; 3) calculate the pairwise distance between $\boldsymbol{X}_i$ and $\boldsymbol{X}_1 + \boldsymbol{d}_{\mathrm{c}}$; 4) remove coordinates with a pairwise distance $|\boldsymbol{d}|$ larger than a set threshold; 5) repeat steps 2-4 five times and save the final channel shift; 6) repeat steps 2-5 for each channel. Second, pair the coordinates: 1) for each coordinate in the current channel, find its closest coordinate in the reference channel corrected by channel shift, pair the two coordinates if their distance is less than 5 pixels; 2) remove unpaired coordinates from all paired channels; 3) repeat steps 1-2 for all channels.

##### *1.3 Segmentation for SMLM data*

Segmentation for *in situ* PSF learning is to select and crop the candidate emitters from the raw SMLM data. For each frame, we first subtract a background image, which is estimated as the average of 100 frames after the current frame. This step is to remove unblinking emitters or large aggregates that can last for a long time, so that the remaining emitters are mostly from single fluorophores. Then a difference of Gaussian filter (i.e. bandpass filter) is applied to the background subtracted image $I_{\mathrm{raw}}$,

$$I_{\mathrm{filter}} = I_{\mathrm{raw}}(x,y) \otimes G(x,y,0.75\sigma_x,0.75\sigma_y) - I_{\mathrm{raw}}(x,y) \otimes G(x,y,\sigma_x,\sigma_y), \tag{1.1}$$

where $G$ represents a 2D Gaussian function with standard deviations defined by $\sigma_x, \sigma_y$, which are user defined parameters. The resulting smoothed image is then filtered by a maximum filter of a set kernel size and the candidate emitters are selected following the same method as for bead data. Usually, 10,000 to 50,000 emitters are selected for one dataset. Different from bead data, here each selected emitter is a 2D image instead of a 3D stack. The axial position of each single molecule emitter is unknown, while the axial step size within the 3D stack is known for bead data. This additional unknown parameter for each emitter complicates the *in situ* PSF learning.

For SMLM data from multi-channel systems a similar coordinate pairing process as described above was applied. However, the method for finding the channel shift is image-based instead of coordinate-based: 1) set channel 1 as the reference channel; 2) calculate the maximum intensity projection (MIP) image along the frame dimension for each channel; 3) calculate the channel shift from cross-correlation (CC) between the MIP images of the current channel and the reference channel. The CC image is set to have the same dimension as the MIP image; 4) the channel shift is the center of the MIP image minus the coordinate of the maximum pixel value of the CC image. 5) repeat steps 3-4 for all channels.

#### 2. PSF modelling in the spatial domain

##### *2.1 PSF learning for single-channel system*

###### *2.1.1 Calculation of initial values*

The initial position for each bead is

$$x_i = 0, y_i = 0, z_i = 0. \tag{2.1}$$

The background value for each bead is estimated as

$$b_i = \min\left(D_i(x,y,z) \otimes G\big(x,y,z,\sigma_x = \sigma_y = \sigma_z = 2\big)\right), \tag{2.2}$$

where $D_i(x,y,z)$ is the cropped 3D image stack of bead $i$. G is a 3D Gaussian kernel with its standard deviations in $x, y$ and $z$ equal to $\sigma_x$, $\sigma_y$, and $\sigma_z$ pixels respectively. $\otimes$ denotes convolution. The minimum is taken over all voxels of the cropped and filtered region. The initial background for each bead is set to the median value of $b_i$ from all bead data,

$$b_{\text{init}} = \underset{i}{\text{median}}(b_i). \tag{2.3}$$

The initial photon value for each bead is estimated as

$$s_{\text{i}}^0 = \underset{z}{\text{avg}}\left(\sum_{x,y} D_i(x,y,z) - b_i\right). \tag{2.4}$$

The initial PSF model is a 3D array with each element set to $1/N_s^2$ with $N_s$ the ROI size. For example, if the ROI size is $N_s \times N_s = 21 \times 21$ pixels, the initial value is 0.0023. This assumes that the sum of each axial slice of the PSF model is one. In fact, for most data, $N_s$ is around 15 to 31, we set the initial value to 0.002 for simplicity.

###### *2.1.2 Calculation of the forward model*

The PSF model at a given bead location $(x_i,\ y_i,\ z_i)$ can be calculated as follows,

$$U_{\text{shift},\,i}(x - x_i, y - y_i, z - z_i) = \mathcal{F}_{3D}^{-1}\big(\widetilde{U} \cdot e^{i\varphi_{\text{shift}}}\big) \tag{2.5}$$

where $\widetilde{U}$ is the OTF, equal to the 3D Fourier transform of the PSF model $U(x,y,z)$,

$$\widetilde{U} = \mathcal{F}_{3D}\big(U(x,y,z)\big). \tag{2.6}$$

The shifting phase is calculated from,

$$\varphi_{\text{shift}} = 2\pi\big(q_x x_i + q_y y_i + q_z z_i\big). \tag{2.7}$$

where $q_x, q_y, q_z$ are the Cartesian coordinates of the frequency space as given by,

$$q_x = \frac{x}{L_x}, \qquad q_y = \frac{y}{L_y}, \qquad q_z = \frac{z}{L_z}, \tag{2.8}$$

where $L_x, L_y$ and $L_z$ are length of the PSF model in $x, y$ and $z$ directions. Considering the bead size, we modified the above equation to,

$$U_{\text{blur},\,i}(x - x_i, y - y_i, z - z_i) = \mathcal{F}_{3D}^{-1}\left(\widetilde{U} \cdot g(q)\ \cdot e^{i\varphi_{\text{shift}}}\right). \tag{2.9}$$

Here we model the bead as a sphere and $g(q)$ is the analytical Fourier transform of a solid sphere of radius $r_0$,

$$g(q) = \frac{J_{\frac{3}{2}}(2\pi q r_0)}{(q r_0)^{\frac{3}{2}}} r_0^3, \tag{2.10}$$

where $J$ is the Bessel function of the first kind and $q$ is the is the spherical coordinate in the frequency space and is calculated from

$$q = \sqrt{q_x^2 + q_y^2 + q_z^2}. \tag{2.11}$$

To ensure the summation of $U_{\text{blur}}$ is close to $U_{\text{shift}}$, the $g(q)$ is normalized by its maximum value.

Considering the photon $s_i$ and background $b_i$, the final forward model for each bead stack is

$$U_i = U_{\text{blur},i} \cdot s_i + b_i. \tag{2.12}$$

##### *2.1.3 Variable scaling*

We used the L-BFGS-B method provided by the optimize package in SciPy for optimization. During optimization, the gradients of the loss function with respect to the variables are calculated for each iteration. However, the gradient values are at different scales for different types of variables. To ensure all variables can be updated at equal rate during each iteration, we scaled each type of variables by a given factor and replace the original variables in the forward model through variable substitution, for example,

$$\begin{aligned} U &= \frac{U}{w_U} w_U = U_w w_U, \\ s_i &= \frac{s_i}{w_s} w_s = s_{w,i} w_s, \\ b_i &= \frac{b_i}{w_b} w_b = b_{w,i} w_b, \end{aligned} \tag{2.13}$$

where $U_w, s_{w,i}, b_{w,i}$ are scaled variables, with respect to which the gradient will be calculated, and $w_U, w_s, w_b$ are scaling factors for the PSF model, intensity and background (see SI Tables for settings of scaling factors for different PSF models). Variable scaling (or parameter scaling) is critical in PSF learning, which ensures the optimization achieves global minimum and all variables can be optimized.

##### *2.1.4 Calculation of the Loss function*

Our loss function for voxel-based learning is constructed based on the mean square error (MSE) between the forward model and the bead data. To let the variables satisfy additional constrains, several terms were added to the loss function. The final loss function is calculated from,

$$\begin{aligned} loss = a_1 MSE_1 + a_2 MSE_2 + a_{\text{sm}} f_{\text{sm}} + a_{\text{edge}} f_{\text{edge}} + a_{\text{drift}} f_{\text{drift}} \\ + \beta(a_{\text{Umin}} f_{\text{Umin}} + a_{\text{bmin}} f_{\text{bmin}} + a_{\text{smin}} f_{\text{smin}} + a_{\text{norm}} f_{\text{norm}}), \end{aligned} \tag{2.14}$$

where $a_{\text{x}}$ are the weighting factors for each loss term. To gradually increase the importance of some constrains during iteration, an additional factor $\beta$ is multiplied, which increases every iteration by $\beta_{n+1} = 1.1\beta_n, \beta_0 = 1$.

Here we will explain every term explicitly. We used two MSE terms,

$$MSE_1 = \frac{\underset{x,y,z,i}{\text{avg}}\,(U_i - D_i)^2}{\underset{x,y,z,i}{\text{avg}}\,(D_i)},$$

$$MSE_2 = \underset{i}{\text{avg}}\left(\frac{\sum_{x,y,z}(U_i - D_i)^2}{N_z \cdot \underset{x,y,z}{\max}(D_i^2)}\right), \tag{2.15}$$

where $N_z$ is the number of slices in z for each bead data and $z$ is from 2 to $N_z - 1$. From simulated data, we found that $MSE_1$ induces a more accurate estimation of the emitter's photon, while $MSE_2$ has less emphasis on bright bead data (which are often caused by bead aggregates) and results in better position estimation. Here we set $a_1 = 1$ and $a_2 = 200$, so that the two MSE terms have similar values.

To reduce the effect of data noise and to obtain a smooth PSF model, we calculated the first difference of the PSF model along z and define,

$$f_{\text{sm}} = \sum_{x,y,z}\left(\frac{U(x,y,z+\Delta z) - U(x,y,z)}{\Delta z}\right)^2 \tag{2.16}$$

where $\Delta z = 1$ pixel (equal to 1 frame of the bead stack) and $a_{\text{sm}} = 1$.

As the PSF model is a finite 3D array, shifting of the PSF model in $x, y$ and $z$ will create abrupt value change at the edges especially along the axial dimension. Therefore, in the calculation of $MSE_1$and $MSE_2$, we ignored the first and the last axial slices in $U_i$ and $D_i$. We then add a loss term to constrain those edge values of the PSF model,

$$f_{\text{edge}} = \sum_{x,y}(U_{z=1} - U_{z=2})^2 + \left(U_{z=N_z} - U_{z=N_z-1}\right)^2. \tag{2.17}$$

This term constrains the border values along the axial direction to its neighboring values along z. We note that although we only ignore the edge slices at $z = 1$ and $z = N_z$ in the MSE calculations, our method can estimate axial shift >1 pixels. Here we set $a_{\text{edge}} = 0.01$.

When sample drift is considered (see section 2.1.5), $f_{\text{drift}}$ is equal to the $L^1$ norm of all drift rates over the bead data, this term is to constrain the estimated drift rates to be close to zero, to avoid adding an arbitrary constant drift rate.

The next three terms serve to constrain the values of PSF model, photon and background to be positive,

$$f_{\text{Umin}} = \sum_{x,y,z}\min(U(x,y,z),0)^2,$$

$$f_{\text{smin}} = \sum_{i}\min(s_i,0)^2, \tag{2.18}$$

$$f_{\text{bmin}} = \sum_{i}\min(b_i,0)^2.$$

Here min denotes element-wise minimum comparing each value in an array with zero. As default we set $a_{\text{Umin}} = 1, a_{\text{smin}} = 1, a_{\text{bmin}} = 1$.

The last term $f_{\text{norm}}$ in the loss function is to constrain the sum of each axial slice of the PSF model to a constant value. This is because the total photon of a single emitter should remain constant at different axial positions due to energy conservation. Therefore, we define,

$$f_{\mathrm{norm}} = \operatorname{avg}_{z}\left(\sum_{x,y} U - \sum_{x,y,z}\frac{U}{N_z}\right)^2. \quad (2.19)$$

This term is often used together with the estimation of $z$ dependent photon fluctuation (see section 2.1.5), to account for photo-bleaching and fluctuation of illumination intensity during the data collection. Normally, we set $a_{\mathrm{norm}} = 0$.

##### *2.1.5 Optional learning variables*

The instabilities of the system, such as drift, illumination fluctuation and photo-bleaching, increase the variations in PSF model. To address this issue, our learning method provides optional variables that partially incorporate the system instabilities.

One option is to estimate the lateral drift during the collection of one bead stack. We define the bead's lateral position at each axial slice as,

$$x_{i,z} = g_{x,i} z, \quad y_{i,z} = g_{y,i} z, \quad (2.20)$$

where $z = 1,2 \ldots N_z$, and $g_{x,i}$ and $g_{y,i}$ are additional learning variables that define the drift rates along $x$ and $y$ for each bead stack. To apply the $z$ dependent lateral drift, the obtained forward model is shifted slice wise through a 2D Fourier transform,

$$U_{\mathrm{drift},i}\left(x - x_{i,z}, y - y_{i,z}, z\right) = \mathcal{F}_{2D}^{-1}\left[\mathcal{F}_{2D}\left(U_i(x,y,z)\right) e^{i2\pi(k_x x_{i,z} + k_y y_{i,z})}\right]. \quad (2.21)$$

Due to additional number of Fourier transforms, with drift estimation the learning speed slows down by ~10%.

Another option is to incorporate $z$ dependent photon fluctuation. With this option, the intensity variable for each bead stack becomes a vector, including one variable $s_{i,z}$ for each axial slice. The initial value for $s_{i,z}$ are the same and equal to the initial value for $s_i$. The calculation for the forward model stays the same, except replacing $s_i$ with $s_{i,z}$. With $z$ dependent photon variable, it is possible to apply $f_{\mathrm{norm}}$ in the loss function, therefore under this option one can set $a_{\mathrm{norm}} = 1$. However, we found that adding $f_{\mathrm{norm}}$ sometimes leads to early termination of the optimization function. Therefore, we advise that when optimization terminates at iteration number less than 60, set $a_{\mathrm{norm}} = 0$.

#### ***2.2 PSF learning for a multi-channel system***

For multi-channel learning, first a single-channel learning is performed for each channel. We selected the first channel as the reference channel, then a transformation matrix $T_l$ (affine transform) of the reference channel to each target channel $l$ was calculated from the estimated positions. The obtained matrix was set as the initial value of $T_l$. For multi-channel learning, variables include positions, detected photons and the optional drift rates from the reference channel, and backgrounds and PSF models from all channels, and the transformation matrices for all target channels (exclude the reference channel). The forward model of a multi-channel system calculates the forward-model of each channel as in single-channel learning, and the $x$ and $y$ positions for the target channel are calculated from

$$[x_{il}, y_{il}, 1] = [x_{i1}, y_{i1}, 1]T_l \quad (2.22)$$

where $i$ is the index of the bead and $l$ is the index of the target channel. As all channels share the same detected photon estimates, the relative intensity ratio between the target channel and the reference channel will be automatically incorporated in the learned PSF models.

#### ***2.3 PSF learning for a 4Pi-SMLM system***

4Pi-SMLM (short as 4Pi in the following text) is a multi-channel system. To use the same framework of the multi-channel learning described in section 2.2, we generated a single-channel learning procedure for the 4Pi system.

##### *2.3.1 Single-channel learning of the 4Pi-PSF*

**Calculation of initial values**. Each 4Pi bead data contains three z-stacks, collected at three different piston phases $\varphi_p = [2\pi/3, 0, -2\pi/3]$ applied by a deformable mirror (DM). We formulated our 4Pi-PSF model based on the IAB model[4], where 4Pi-model is defined by

$$\begin{aligned} U(x,y,z,\varphi) &= I(x,y,z) + A(x,y,z)\cos(\varphi) + B(x,y,z)\sin(\varphi) \\ &= I(x,y,z) + 2\mathrm{Re}[E(x,y,z)e^{i\varphi}], \end{aligned} \tag{2.23}$$

So that each 4Pi-PSF model contains three 3D matrices representing $I$ and $\mathrm{Re}(E)$ and $\mathrm{Im}(E)$. The component $I$ describes the incoherence of the 4Pi-PSF, while component $E$ is a complex matrix and includes the coherence feature of the 4Pi-PSF. The $A$ and $B$ matrices defined in the IAB model are related to $E$ by $A = 2\mathrm{Re}(E), B = -2\mathrm{Im}(E)$.

Component $I$ and $E$ of each 4Pi bead data are obtained as follows,

$$\begin{aligned} G_{i0} &= \sum_{p=0}^{2} D_{ip} e^{i2\pi(p-1)/3}, \\ G_{i1} &= \sum_{p=0}^{2} D_{ip}, \end{aligned} \tag{2.24}$$

where $D_{ip}$ represents $p$th z-stack in $i$th 4Pi bead data, and $G_{i0}$ and $G_{i1}$ are Fourier components of $D_{ip}$ along the $p$ dimension. Note that the symbol $i$ in subscript denotes index and in exponential denotes complex value. Then we have,

$$\begin{aligned} I_{i,\mathrm{data}} &= G_{i1}/3, \\ E_{i,\mathrm{data}} &= G_{i0}/3. \end{aligned} \tag{2.25}$$

Similar to single-channel learning of incoherent PSFs, we used $I_{i,\mathrm{data}}$ instead of $D_i$ to calculate the initial values of photon and background. The initial position $(x_i, y_i, z_i)$ and phase $\varphi_i$ of each 4Pi bead data are set to zero.

We also set the piston phase $\varphi_p$ as a variable to compensate residue errors in DM calibration. The initial value of $\varphi_p = [2, 0, -2]$ and all bead data share the same $\varphi_p$.

The initial 4Pi-PSF model contains three 3D matrices, where we separate the real part and the imaginary part of the component $E$ to avoid complex variables, and each element of those matrices are set to a constant value,

$$\begin{aligned} I(x,y,z) &= 0.002, \\ \mathrm{Re}(E) = \mathrm{Im}(E) &= 0.001/\sqrt{2}. \end{aligned} \tag{2.26}$$

Therefore, for single channel learning of the 4Pi-PSF, the variables are $x_i, y_i, z_i, \varphi_i, s_i, b_i, I, \mathrm{Re}(E), \mathrm{Im}(E)$, and $\varphi_p$.

**Calculation of the forward model**. First, we obtain the 4Pi-PSF model from $I$ and $E$ as follows,

$$U_p = I(x,y,z) + 2\mathrm{Re}\left[E(x,y,z)e^{i\varphi_z + i\varphi_i + i\varphi_p}\right], \tag{2.27}$$

where $\varphi_z = 2\pi z/z_T$ describing the modulation feature of the 4Pi-PSF along the axial dimension. Thus, $E$ has a slow modulation in $z$. The modulation period $z_T$ is a user-defined constant value, with a unit of pixel. $z_T$ can be estimated from the modulation period of the 4Pi bead data, and we slightly adjust the value to minimize the loss. Here we used $z_T$ equals to $260\ \mathrm{nm}/\delta z$ and $290\ \mathrm{nm}/\delta z$ for bead data measured from 600 nm and 676 nm channels respectively, where $\delta z$ is the step size in $z$.

To shift the PSF model in x, y and z, we apply the same shift to both $I$ and $E$ as follows,

$$I_{\mathrm{shift},i}(x - x_i, y - y_i, z - z_i) = \mathcal{F}^{-1}\left[\mathcal{F}\big(U(x,y,z)\big)e^{i\varphi_{\mathrm{shift}}}\right], \tag{2.28}$$

$$E_{\text{shift},i}(x-x_i,y-y_i,z-z_i)=\mathcal{F}^{-1}[\mathcal{F}(E(x,y,z)e^{i\varphi_z+i\varphi_i})e^{i\varphi_{\text{shift}}}],$$

Then the forward model for each bead data becomes,

$$U_{\text{shift},i,p}=I_{\text{shift},i}+2\text{Re}\left[E_{\text{shift},i}e^{i\varphi_p}\right] \tag{2.29}$$

The shifted forward model is then blurred with the bead kernel,

$$U_{\text{blur},i,p}=\mathcal{F}_{3D}^{-1}\left[\mathcal{F}_{3D}\left(U_{\text{shift},i,p}\right)\cdot g(q)\right] \tag{2.30}$$

Considering the photon and background, the final forward model for each bead data is

$$U_i=U_{\text{blur},i,p}\cdot s_i+b_i. \tag{2.31}$$

As there are three piston phases $\varphi_p$, $p=0,1,2$, the final forward model for each 4Pi bead data contains three z-stacks.

**Calculation of the loss function**. The loss function for learning 4Pi-PSF is the same as the one for learning incoherent PSF. Here we explicitly give the calculation of each loss term,

$$MSE_1=\frac{\underset{x,y,z,p,i}{\text{avg}}\left(U_{ip}-D_{ip}\right)^2}{\underset{x,y,z,p,i}{\text{avg}}\left(D_{ip}\right)},$$

$$MSE_2=\underset{i,p}{\text{avg}}\left(\frac{\sum_{x,y,z}\left(U_{ip}-D_{ip}\right)^2}{N_z\cdot\underset{x,y,z}{\max}(D_{ip}^2)}\right),$$

$$\begin{aligned}f_{\text{sm}}=&\sum_{x,y,z}\left(\frac{I(z+\Delta z)-I(z)}{\Delta z}\right)^2+\sum_{x,y,z}\left(\frac{\text{Re}[E(z+\Delta z)]-\text{Re}[E(z)]}{\Delta z}\right)^2\\&+\sum_{x,y,z}\left(\frac{\text{Im}[E(z+\Delta z)]-\text{Im}[E(z)]}{\Delta z}\right)^2\end{aligned} \tag{2.32}$$

$$\begin{aligned}f_{\text{edge}}=&\sum_{x,y}(I_{z=1}-I_{z=2})^2+\left(I_{z=N_z}-I_{z=N_z-1}\right)^2\\&+\sum_{x,y}(\text{Re}(E)_{z=1}-\text{Re}(E)_{z=2})^2+\left(\text{Re}(E)_{z=N_z}-\text{Re}(E)_{z=N_z-1}\right)^2\\&+\sum_{x,y}(\text{Im}(E)_{z=1}-\text{Im}(E)_{z=2})^2+\left(\text{Im}(E)_{z=N_z}-\text{Im}(E)_{z=N_z-1}\right)^2,\end{aligned}$$

$$f_{\text{Umin}}=\sum_{x,y,z}\min(I-2|E|,0)^2,$$

$$f_{\text{norm}}=\underset{z}{\text{avg}}\left(\sum_{x,y}I-\sum_{x,y,z}\frac{I}{N_z}\right)^2.$$

The remaining terms are defined the same as in learning incoherent PSFs.

##### 2.3.2 Multi-channel learning of 4Pi-PSF

Multi-channel learning of the 4Pi-PSF follows the same framework of multi-channel learning of the incoherent PSF. It includes single-channel learning of the 4Pi-PSF from each of the four channels and initial calculation of the transformation matrices between the target channels to the reference channel. The variables include positions, phases, photons, piston phases and the optional drift rates from the reference channel, backgrounds, $I$, real part and imaginary part of $E$ from all channels, and the transformation matrices for all target channels (exclude the reference channel).

#### 2.4 Learning of lattice light-sheet PSF

Learning of lattice light-sheet (LLS) PSFs is based on single-channel PSF learning. Due to the geometry of the LLS microscope, moving of the sample stage results in translating the bead in both z and one of the lateral dimensions, respective to the detection objective or the camera. Therefore, the forward model in LLS PSF needs to incorporate this lateral translation. The method is similar to adding drift to the forward model (see section 2.1.5). However, here instead of estimating the drift rate, a constant drift is applied to all bead data. This constant drift is termed as skew constants, $g_{sx}$ and $g_{sy}$ , and are equal to the translation (in pixels) of a bead along the x and y axis per axial slice. The skew constants are given by the user, which are related to the angle between the detection axis and the sample plane. As the lateral translation can be tens of pixels over the whole z stack, the raw bead stack can take ~60 pixels in one of the lateral dimensions. To reduce the PSF size, we deskewed the raw bead stack where each axial slice is shifted by integer pixels as follows,

$$\begin{aligned} \Delta x_{\text{int}} &= -\text{round}(g_{sx}z), \\ \Delta y_{\text{int}} &= -\text{round}(g_{sy}z), \end{aligned} \tag{2.33}$$

The deskewed bead data are used in the learning and a 2D shift is applied to the forward model to account for the skew effect,

$$U_{\text{skew},i}\left(x - x_{i,z}, y - y_{i,z}, z\right) = \mathcal{F}_{2D}^{-1}\left[\mathcal{F}_{2D}(U_i(x, y, z))e^{i2\pi(k_x x_{i,z} + k_y y_{i,z})}\right], \tag{2.34}$$

where

$$\begin{aligned} x_{i,z} &= g_{sx}z - \text{round}(g_{sx}z), \\ y_{i,z} &= g_{sy}z - \text{round}\left(g_{sy}z\right). \end{aligned} \tag{2.35}$$

The learned PSF model is the deskewed PSF as seen from a conventional single-objective system.

#### 2.5 Localization test

To evaluate the accuracy of the learned PSF model, we performed a localization test on the same bead data used for learning. Details of the localization algorithms are described in previous studies[1,5]. In brief, first we calculated the cubic spline coefficients of the learned PSF model, where each voxel contains $64 \times N_{\text{channel}}$ coefficients, while for 4Pi-PSF, it is $64 \times 3 \times N_{\text{channel}}$ coefficients, where $N_{\text{channel}}$ is the number of channels; then a maximum likelihood estimation (MLE) is used to estimate the $x, y, z$ positions (and $\varphi$ for 4Pi-PSF) for each 2D bead image using the spline PSF model (see section 7). We evaluated the accuracy of the learned PSF model using the MSE of the axial localization,

$$MSE_z = \operatorname*{avg}_z \left( z_{\text{bias},i,z} - \operatorname*{median}_i z_{\text{bias},i,z} \right)^2, \tag{2.36}$$

where the localization bias in z is calculated from,

$$z_{\text{bias},i,z} = z_{i,z} - z_{GT,i,z}, \tag{2.37}$$

where $z_{GT}$ is the ground truth position and equals to the stage position, the subscripts $i$ and $z$ denote the indices of the beads and the axial slices in each bead stack respectively. For 4Pi bead data, $z_{i,z} = \operatorname*{avg}_p z_{i,z,p}$, where $p$ is the index of the piston phases. $MSE_z$ is a vector of $N_{\text{bead}} \times 1$ elements where $N_{\text{bead}}$ is the number of the bead data.

##### *2.6 Outlier removal*

Due to the presence of field-dependent aberrations of the imaging system and the aggregates within the bead sample, it is necessary to remove outlier beads. We used three criteria to identify outliers. The first one is based on the MSE between the forward models and the bead data,

$$MSER = \frac{MSE}{\text{median}(MSE)}\ , \tag{2.38}$$

Where $MSE$s for incoherent PSFs (e.g. astigmatism PSF) and 4Pi PSFs were calculated from,

$$MSE_w = \frac{\underset{x,y,z}{\text{avg}}(U_i - D_i)^2}{\underset{x,y,z}{\text{avg}}(D_i)} \text{ and } MSE_{4\text{Pi}} = \frac{\underset{x,y,z,p}{\text{avg}}\left(U_{ip} - D_{ip}\right)^2}{\underset{x,y,z,p}{\text{avg}}\left(D_{ip}\right)}. \tag{2.39}$$

Both $MSE$ and $MSER$ are vectors of $N_{bead} \times 1$ elements. The second criterion is based on the MSE of the axial localization bias, $MSE_z$ calculated above,

$$MSER_z = \frac{MSE_z}{\text{median}(MSE_z)}. \tag{2.40}$$

The third criterion is based on the estimated photons from the PSF learning. This criterion is to remove bead measurements that are too bright or too dim, and can be calculated from,

$$R_s = \frac{[s - \text{median}(s)]^2}{s \cdot \text{median}(s)}, \tag{2.41}$$

where $s$ contains the estimated photon $s_i$ of each bead data. Both $s$ and $R_s$ are vectors of $N_{bead} \times 1$ elements.

The bead data are considered as outliers if their metric values of above defined criteria are greater than the user defined thresholds, for example,

$$MSER_i > 3 \text{ or } MSER_{z,i} > 3 \text{ or } R_{s,i} > 1.5. \tag{2.42}$$

After outlier removal, the remaining bead data were used to re-learn the PSF model. The obtained PSF model and its corresponding spline coefficients (and the transformation matrices for multi-channel learning) from the second learning step will be used for localization of SMLM data.

#### 3. PSF modelling in the Fourier domain

##### *3.1 Pupil-image based PSF learning*

In this representation, pupil function is represented by two 2D matrices.

###### *3.1.1 Calculation of the forward model*

The PSF model is calculated from a Fourier transform of the pupil function,

$$U(x - x_i, x - y_i, z - z_i) = \left|\mathcal{F}_{\text{czt}}\left[h\left(k_x, k_y\right)e^{-i2\pi k_z(z-z_i)}e^{i2\pi(k_x x_i + k_y y_i)}\right]\right|^2,$$
$$k_z = \sqrt{k^2 - k_x^2 - k_y^2} \tag{3.1}$$

where $h\left(k_x, k_y\right)$ is the pupil function and can be written as,

$$h\left(k_x, k_y\right) = T_a A\left(k_x, k_y\right)e^{i\Phi(k_x,k_y)}, \tag{3.2}$$

where $A$ and $\Phi$ are the magnitude and phase components of the pupil function, each of them is a 2D array. $T_a$ is the apodization factor of the objective lens and is equal to

$$T_a = \frac{\sqrt{\cos\theta_{\text{imm}}}}{\cos\theta_{\text{med}}}, \tag{3.3}$$

where $\theta_{\text{med}}$ and $\theta_{\text{imm}}$ are the angles of the optical rays in the sample medium and the immersion medium respectively, which are limited by the numerical aperture (NA) of the objective lens. The $k_x, k_y, k_z$ are the Cartesian components of the wave vector $\boldsymbol{k}$, with its magnitude $k = n_{\text{imm}}/\lambda$, where $n_{\text{imm}}$ is the refractive index of the immersion medium and $\lambda$ is the central wavelength of the emission filter. And $e^{-i2\pi k_z(z-z_i)}$ is the propagation term and accounts for the defocus phase. With pupil-based PSF learning, we can directly add the bead position $(x_i, y_i, z_i)$ to the pupil function and eliminate the edge effect from voxel-based PSF learning. And $z$ denotes the stage positions and is a vector $N_z \times 1$ elements. Here we used Chirp Z-transform based on Bluestein's algorithm to calculate the 2D Fourier transform of the pupil function, denoted as $\mathcal{F}_{\text{czt}}$. The benefit of using the BS algorithm is that the size of the pupil function is not dependent on the pixel size of the camera and a pupil size of 64 × 64 pixels can represent a reasonable sampling in the pupil function.

To account for the effect of fluorescence dipole emission, pixilation, bead size, dispersion and chromatic aberration, the PSF model is blurred by,

$$U_{\text{blur}} = \mathcal{F}_{3D}^{-1}\left[\mathcal{F}_{3D}(U) \cdot g(q) \cdot e^{-i2(\sigma_x q_x \pi)^2 - i2(\sigma_y q_y \pi)^2}\right], \tag{3.4}$$

where $g(q)$ is the bead kernel in Fourier space as defined in Eq 2.10 and $\sigma_x$ and $\sigma_y$ are the standard deviation (in pixel unit) of a 2D Gaussian kernel in real space. By including the photon count and background of each bead stack, the final forward model for pupil-image based learning is

$$U_i = U_{\text{blur}} \cdot s_i + b_i. \tag{3.5}$$

##### *3.1.2 calculation of the Loss function*

The loss function for pupil-image based learning is,

$$loss = a_1 LL + a_{\text{sm}} f_{\text{sm}} + a_{\text{drift}} f_{\text{drift}} + a_{\text{norm}} f_{\text{norm}} + \beta(a_{\text{bmin}} f_{\text{bmin}} + a_{\text{smin}} f_{\text{smin}}), \tag{3.6}$$

where $LL$ is based on the log-likelihood function, assuming the converted pixel values (see section 1) follow the Poisson distribution,

$$LL = \underset{x,y,z,i}{\text{avg}}\, U_i - D_i - D_i \log(U_i) + D_i \log(D_i). \tag{3.7}$$

Note that pure data terms such as $D_i$ and $D_i \log(D_i)$ are present for improved numerical stability to sum up smaller values. Since they are constant, they do not influence the position of the optimum. Here $f_{\text{sm}}$ is to smooth the pupil magnitude and is defined as,

$$f_{\text{sm}} = \sum_{k_x, k_y} \left(\frac{\Delta A}{\Delta k_x}\right)^2 + \left(\frac{\Delta A}{\Delta k_y}\right)^2. \tag{3.8}$$

where $\Delta k_x, \Delta k_y$ and $\Delta k_z$ are equal to one pixel in the Fourier space. The smoothness of the pupil phase is controlled by $f_{\text{norm}}$, which constrains the sum of the in-focus PSF close to one,

$$f_{\text{norm}} = \min\left(\frac{\sum U(z=0)}{\sum U_{\text{ideal}}(z=0)} - 0.97, 0\right)^2, \tag{3.9}$$

where $U_{\text{ideal}}$ is the unaberrated PSF with pupil phase equal to zero and $U$ is the PSF calculated from the Fourier transform of the current pupil phase $\Phi$. We found that without $f_{\text{norm}}$, the learned pupil phase tends to have random noise pixels. Those random noise pixels will not affect the shape of the PSF model but will lower its total intensity[6]. The rest of the terms are the same as the ones for voxel-based learning.

The variables for pupil-imaged based learning are $x_i, y_i, z_i, s_i, b_i, A, \Phi, \sigma_x$ and $\sigma_y$. The initial values of $x_i, y_i, z_i, s_i$ were determined as described in voxel-based learning (see section 2.1.1). The initial values for each element in $A$ and $\Phi$ are 1 and 0, respectively. The initial values of $\sigma_x$ and $\sigma_y$ are user defined and set to 0.5 pixel as default.

##### *3.2 Zernike-based PSF learning*

The calculation of the forward model for Zernike-based PSF learning is like the one for pupil-image based PSF learning, except that we decompose the magnitude and the phase components of the pupil function into Zernike polynomials,

$$A = \sum_p C_{A,p} Z_p,$$
$$\Phi = \sum_p C_{\Phi,p} Z_p, \quad (3.10)$$

where $Z_p$ are Zernike polynomials in Noll order. Here we estimate the coefficients $C_{A,p}$ and $C_{\Phi,p}$ for each Zernike polynomial.

As the expansion of pupil functions into Zernike polynomials has an effect of smoothing the pupil function, therefore in the loss function, we omit the smoothing and normalization terms (Eqs. 3.8 and 3.9),

$$loss = a_1 LL + a_{\text{drift}} f_{\text{drift}} + \beta(a_{\text{bmin}} f_{\text{bmin}} + a_{\text{smin}} f_{\text{smin}}). \quad (3.11)$$

The variables for Zernike-based PSF learning are $x_i, y_i, z_i, s_i, b_i, C_{A,p}, C_{\Phi,p}$ and $\sigma$. The initial values $C_{A,p}$ and $C_{\Phi,p}$ are equal to zero, except the first coefficient of pupil amplitude, $C_{A,0} = 1$. The initial values of other variables were determined as described for pupil-image based learning.

##### *3.3 Vectorial PSF model*

To account for the broadening effect of fluorescence dipole emission, we also incorporated the vectorial PSF model in pupil-image and Zernike based PSF learning. Here we assume that the fluorescence bead is an ensemble of freely rotating dipoles, and the vectorial PSF model can be written as,

$$U(x - x_i, y - y_i, z - z_i) = \sum_{\substack{m=x,y \\ n=p_x,p_y,p_z}} \left| \mathcal{F}_{\text{czt}} \left[ h(k_x, k_y) w_{mn} e^{-i2\pi k_z (z-z_i)} e^{i2\pi (k_x x_i + k_y y_i)} \right] \right|^2, \quad (3.12)$$

where $w_{mn}$ is the $m$ component of the electric field at the pupil plane generated by the $n$ component of the dipole moment of the fluorophore. The calculation of $w_{mn}$ is as follows,

$$w_{xn} = P_n \cos\varphi - S_n \sin\varphi,$$
$$w_{yn} = P_n \sin\varphi + S_n \cos\varphi, \quad (3.13)$$

Where $\varphi$ is the angular component in the polar coordinate of the frequency space. $P_n$ and $S_n$ are electric field components in p and s polarizations relative to the incident plane at the sample space,

$$P_{p_x} = T_P \cos\theta_1 \cos\varphi,$$
$$P_{p_y} = T_P \cos\theta_1 \sin\varphi,$$
$$P_{p_z} = -T_P \sin\theta_1,$$
$$S_{p_x} = -T_S \sin\varphi, \quad (3.14)$$
$$S_{p_y} = T_S \cos\varphi,$$
$$S_{p_z} = 0,$$

where $p_x, p_y, p_z$ are the Cartesian components of the dipole moments, $T_P$ and $T_S$ are the total transmission coefficients of p- and s-polarized light. The fluorescence light starting from the dipole emitter propagates through the sample medium, the coverslip and the immersion medium. In this three-layer system, we ignore the multiple reflections at the two interfaces, then $T_P$ and $T_S$ can be calculated from,

$$\begin{aligned} T_P &= \tau_{P12}\tau_{P23}, \\ T_S &= \tau_{S12}\tau_{S23}, \end{aligned} \tag{3.15}$$

where $\tau_{Pij}$ and $\tau_{Sij}$ are the Fresnel transmission coefficients of s- and p-polarized light travel from medium $i$ to medium $j$,

$$\begin{aligned} \tau_{Pij} &= \frac{2n_i\cos\theta_i}{n_i\cos\theta_j + n_j\cos\theta_i}, \\ \tau_{Sij} &= \frac{2n_i\cos\theta_i}{n_i\cos\theta_i + n_j\cos\theta_j}, \end{aligned} \tag{3.16}$$

where $n_i$ and $\theta_i$ are the refractive index and the light propagation angle in medium $i$, and the subscript 1,2,3 denotes the sample medium, the coverslip and the immersion medium respectively.

The remaining calculations of the forward model and the definition of the loss functions are the same as in the pupil-image and Zernike based methods (sections 3.1 and 3.2).

##### *3.4 Fourier domain PSF learning of multi-channel systems*

The PSF learning methods based on the scalar or vectorial PSF models can also be extended to multi-channel systems. Similar as in the voxel-based PSF learning, the x and y positions of the beads between the target channel and the reference channel are related by a 3×3 transformation matrix. The learning variables are positions, photons and the optional drift rates from the reference channel, and backgrounds, the blurring factor, the pupil function or the Zernike coefficients from all channels, and the transformation matrices for all target channels (exclude the reference channel).

##### *3.5 Fourier domain PSF learning for the 4Pi-SMLM system*

The Fourier domain PSF learning methods can also be extended to the 4Pi-SMLM system. The learning framework is the same as in voxel-based 4Pi-PSF learning.

###### *3.5.1 Single-channel learning of pupil based 4Pi-PSF*

Pupil-based model including both pupil-image and Zernike based models.

**Calculation of the forward model**. The 4Pi system collects fluorescence emission from both lower and upper objectives, the coherent superposition of the electric fields (pupil function) from the two emission paths forms a coherent PSF, while the incoherent superposition of the two electric fields forms an incoherent PSF. The 4Pi PSF in each channel is a summation of the coherent and incoherent PSFs,

$$U = \alpha U_I + (1-\alpha)U_w, \tag{3.17}$$

where $U_I$ is the coherent PSF,

$$\begin{aligned} &U_I(x-x_i, y-y_i, z-z_i, \varphi_i) \\ &= \left|\mathcal{F}_{\mathrm{czt}}\left[\left(h_1 e^{i2\pi k_z(z-z_i)+i\varphi_i} + h_2 e^{-i2\pi k_z(z-z_i)+i\varphi_{pl}}\right)e^{i2\pi(k_x x_i + k_y y_i)}\right]\right|^2, \end{aligned} \tag{3.18}$$

where $h_1$ and $h_2$ are pupil functions from the upper and lower emission paths, the term $2\pi k_z(z-z_i)$ is the defocus phase and $k_z$ is defined in Eq. (3.1), $\varphi_i$ is the modulation phase of each PSF data and $\varphi_{pl} = \varphi_p + \varphi_l$ is the piston phase applied by the deformable mirror plus a phase delay ($\varphi_l$, see section 3.5.2) relative to the reference channel. $U_w$ is the incoherent PSF,

$$U(x - x_i, y - y_i, z - z_i)$$
$$= \left|\mathcal{F}_{\mathrm{czt}}\left[h_1 e^{i2\pi k_z(z-z_i)} e^{i2\pi(k_x x_i + k_y y_i)}\right]\right|^2 + \left|\mathcal{F}_{\mathrm{czt}}\left[h_2 e^{-i2\pi k_z(z-z_i)} e^{i2\pi(k_x x_i + k_y y_i)}\right]\right|^2 \quad (3.19)$$

The factor $\alpha$ describes the modulation contrast of the 4Pi-PSF. When $\alpha = 1$, the 4Pi-PSF is completely coherent, when $\alpha < 1$, the 4Pi-PSF is partially coherent. Due to the broad fluorescence emission spectrum and the finite band width of the emission filter, the measured 4Pi-PSF is always partially coherent.

Learning of the 4Pi-PSF model involves learning of both upper and lower pupil functions, $h_1$ and $h_2$, and they can be calculated from,

$$h_1 = T_a A_1 e^{i\Phi_1},$$
$$h_2 = T_a A_2 e^{i\Phi_2}, \quad (3.20)$$

The apodization term $T_a$ is the same as previously defined. For Zernike-based learning, $A_1, A_2, \Phi_1$ and $\Phi_2$ are expressed in terms of Zernike polynomials,

$$X_j = \sum_{p=0}^{N_{\mathrm{zern}}-1} C_{X_j,p} Z_p \ , \quad (3.21)$$

where $j$ can be 1 or 2 denotes upper and lower paths, and $X_j$ can be $A_j$ or $\Phi_j$. Here we set the piston phase in $\Phi_1$ to zero, which is $C_{\Phi_1,0} = 0$, and the relative phase delay between $h_1$ and $h_2$ is estimated from the piston phase in $\Phi_2$, which is $C_{\Phi_2,0}$.

The obtained PSF model will also be blurred by a bead kernel and a Gaussian kernel as for the case of voxel-based 4Pi-PSF learning.

**Calculation of the Loss function**. The loss function of pupil-image based 4Pi-PSF learning is,

$$loss = a_1 LL + a_{\mathrm{sm}} f_{\mathrm{sm}} + a_{\mathrm{drift}} f_{\mathrm{drift}} + \beta(a_{\mathrm{b}} f_{\mathrm{bmin}} + a_{\mathrm{s}} f_{\mathrm{smin}} + a_{\alpha} f_{\alpha\mathrm{min}}), \quad (3.22)$$

where $f_{\mathrm{sm}} = f_{\mathrm{sm1}} + f_{\mathrm{sm2}}$ is to smooth both the lower and upper pupil functions, and $f_{\mathrm{sm1}}$ and $f_{\mathrm{sm2}}$ can be calculated from Eq. (3.8). The term $f_{\alpha\mathrm{min}}$ is to constrain the modulation contrast $\alpha$ to be positive,

$$f_{\alpha\mathrm{min}} = \min(\alpha, 0)^2. \quad (3.23)$$

The remaining terms are the same as in pupil-image based learning for incoherent PSFs (section 3.1).

The loss function for Zernike-based 4Pi-PSF learning is

$$loss = a_1 LL + a_{\mathrm{drift}} f_{\mathrm{drift}} + \beta(a_{\mathrm{b}} f_{\mathrm{bmin}} + a_{\mathrm{s}} f_{\mathrm{smin}} + a_{\alpha} f_{\alpha\mathrm{min}}). \quad (3.24)$$

##### *3.5.2 Multi-channel learning of the 4Pi-PSF in the Fourier domain*

The multi-channel learning of a pupil-based 4Pi-PSF model is like the one for voxel-based learning. To account for the constant phase delay between each channel, we add $\varphi_l$ to $\Phi_2$ in each channel,

$$\varphi_l = -l\frac{\pi}{2}, \ \ l = 0,1,2,3, \quad (3.25)$$

where $l$ denotes the channel index, and the residual phase delay between each channel will be incorporated into the piston component of $\Phi_2$. Above definition of $\varphi_l$ assumes the phase delay increment by $-\pi/2$ over the four channels. User can define the increment to be $-\pi/2$ or $\pi/2$ depending on their systems. The learning variables include positions, phases, photons, piston phases and the optional drift rates of the reference channel, and backgrounds, pupil functions or Zernike polynomials, blurring factor and the modulation contrast from all channels, and the transformation matrices for all target channels (exclude the reference channel).

For Zernike-based learning, the user can choose to link the Zernike coefficients between the four channels, note that we do not link the Zernike coefficients between the lower and upper paths,

$$C_{A_j,l,p} = C_{A_j,0,p}\ , p > 0, l = 0,1,2,3, j = 1,2$$
$$C_{\Phi_j,l,p} = C_{\Phi_j,0,p}\ , p > 3, l = 0,1,2,3, j = 1,2 \tag{3.26}$$

Where, $p$ denotes the Zernike index, $l$ the channel index, $j$ the lower or the upper path. For the magnitude part, only the piston term is unlinked to account for intensity differences between the four channels. For the phase part, the piston phase, x and y tilts and defocus are unlinked, to account for the relative phase delay and the x, y and z shifts between the four channels. Although the x and y shifts between channels can be accounted for by the transformation matrices, we noticed that the relative x and y shifts between the upper and the lower objectives are also channel dependent. Therefore, we unlink the x and y shifts of the Zernike coefficients between the four channels.

##### *3.6 Learning of field dependent PSFs*

For high-NA objectives, field dependent (FD) aberrations become prominent with large field of views (FOVs). The PSF model within a FOV of 10 × 10 µm can often be treated as spatial invariant. However, in a FOV of 100 × 100 µm, a single PSF model sometimes can no longer represent all PSF patterns. Here we extend our method to learn a spatial variant PSF model across a large FOV. The forward model for learning the FD aberrations is based on Zernike-based PSF models (either scalar or vectorial model). For each Zernike coefficient, a 2D map is learned instead of a scalar value, where each pixel value of the map represents the aberration amplitude at the field location defined by the pixel coordinate. Therefore, in FD PSF learning, a set of 2D aberration maps were learned. The forward model was calculated similarly as in Zernike-based PSF learning (section 3.2), except that the Zernike coefficients are bead-dependent and were calculated from a linear interpolation of the aberration map,

$$C_{A,i,p} = Z_{\mathrm{map},A,p}(X_i, Y_i)\ ,$$
$$C_{\Phi,i,p} = Z_{\mathrm{map},\Phi,p}(X_i, Y_i)\ . \tag{3.27}$$

Here $X_i$ and $Y_i$ are the pixel coordinates of the $i$th bead in the raw camera frame. $Z_{\mathrm{map},A,p}$ and $Z_{\mathrm{map},\Phi,p}$ are the aberration maps for the $p$th Zernike polynomial of the amplitude and the phase parts of the pupil function, respectively. Each aberration map is a 2D matrix of $N \times N$ pixels, where $N$ is user defined, e.g. $N = 20$ means the FOV is divided by 20 × 20 subregions. The larger the $N$, the finer the aberration map, however, as the aberration maps vary smoothly across the FOV, $N$ can be chosen so that each subregion takes ~10 × 10 µm.

The loss function for learning the FD PSF model is,

$$loss = a_1 LL + a_{\mathrm{sm}} f_{\mathrm{sm}} + a_{\mathrm{drift}} f_{\mathrm{drift}} + \beta(a_{\mathrm{bmin}} f_{\mathrm{bmin}} + a_{\mathrm{smin}} f_{\mathrm{smin}})\ , \tag{3.28}$$

where $f_{\mathrm{sm}}$ is to smooth the aberration maps,

$$f_{\mathrm{sm}} = \sum_{k_x,k_y,p} \left(\frac{\Delta Z_{\mathrm{map},A,p}}{\Delta k_x}\right)^2 + \left(\frac{\Delta Z_{\mathrm{map},A,p}}{\Delta k_y}\right)^2 + \left(\frac{\Delta Z_{\mathrm{map},\Phi,p}}{\Delta k_x}\right)^2 + \left(\frac{\Delta Z_{\mathrm{map},\Phi,p}}{\Delta k_y}\right)^2, \tag{3.29}$$

And the rest terms are the same as the ones in Zernike-based PSF learning (section 3.2).

##### *3.7 Learning of refractive index mismatch aberrations*

Here we apply our learning algorithm to data from beads embedded in an agarose gel to extract the bead's axial position relative to the coverslip. Fluorescence beads were immobilized throughout the agarose gel and are imaged through an oil immersion objective. The refractive indices of the agarose gel and the immersion oil are 1.334 and 1.516. Therefore, the deeper the bead inside the gel, the larger the refractive index mismatch aberration. The forward model, e.g. vectorial model, to incorporate the index mismatch aberration is,

$$U(x - x_i, y - y_i, z_i, z_s + z) = \sum_{\substack{m=x,y\\ n=p_x,p_y,p_z}} \left|\mathcal{F}_{\mathrm{czt}}\left[h(k_x,k_y) w_{mn} e^{i2\pi k_{z\mathrm{med}} z_i - k_z(z_s+z)} e^{i2\pi(k_x x_i + k_y y_i)}\right]\right|^2, \tag{3.30}$$

The term $e^{i2\pi(k_{z\mathrm{med}} z_i - k_z z_s)}$ accounts for the index mismatch aberration,

$$k_{z\text{med}} = \sqrt{k_{\text{med}}^2 - k_x^2 - k_y^2}\,,$$

$$k_z = \sqrt{k^2 - k_x^2 - k_y^2}\;, \tag{3.31}$$

$$k_{\text{med}} = \frac{n_{\text{med}}}{\lambda}\;, k = \frac{n_{\text{imm}}}{\lambda}$$

where $n_{\text{med}}$ and $n_{\text{imm}}$denote the refractive indices of the sample medium and the immersion medium, respectively. $z_s$ defines the stage translation relative to the nominal focal position, where the objective is focused at the coverslip. Here we define $z_s = 0$ at the nominal focal position, and $z_s > 0$ when the objective is moved closer to the coverslip, equivalent to when objective is focused at beads away from the coverslip.

And $z_i$ is the bead's axial position to the coverslip, $z$ is a vector of $N_z \times 1$ relative to $z_s$. During the data segmentation, a 3D bead stack was cropped around the 3D pixel coordinate of the bead. The pupil function $h(k_x, k_y)$ was fixed and learned from the bead data collected with bead at the coverslip. Both $z_i$ and $z_s$ are estimation parameters during the learning.

### 4. Learning of *in situ* PSF models

*In situ* PSF learning is to extract the PSF model from raw SMLM data, i.e., camera frames with single blinking emitter patterns. Here our method is based on the vectorial PSF model.

#### *4.1 In situ PSF learning for single-channel systems*

##### *4.1.1 Calculation of initial z values*

As the PSF pattern varies with the emitter's axial position, learning of the *in situ* PSF model is dependent on the initial estimations of the emitter's axial positions. Here we localize all emitters using an initial PSF model. The localization method is based on MLE with a spline-interpolated PSF model (sections 2.5 and 7). Here we provide three options on estimating the initial PSF model.

The first option is to learn the initial PSF model from the bead data collected from the same imaging system. The resulting file can be set as an input parameter for the subsequent *in situ* PSF learning. This method can give an accurate initial z estimation; however, it requires collection of bead data.

The second option is to generate the initial PSF model based on a set of user defined Zernike polynomials. Here the user needs to select the Zernike polynomials that represent the dominant aberrations of the imaging system and set the approximate amplitudes to the selected Zernike polynomials. For example, most 3D-SMLM systems have astigmatism aberration, then the user can set the Zernike polynomial to 5 (Noll order) and its coefficient to 0.5 (radian, its absolute value quantifies the RMS wavefront error). Although this method requires no bead measurement, the user needs to know the dominant aberration of the imaging system. We found that the learning algorithm is quite robust to a wrong initial aberration if the tendency of the initial z positions of single molecules agrees with the real situation. However, the convergence might be a bit slower.

The third option requires minimum knowledge of the imaging system from the user, it will search through a set of Zernike polynomials to determine the dominant aberration. The user only needs to define the search range is within the lower order (5-21) or the higher order Zernike polynomials (22-45). During the searching procedure, a PSF model for each Zernike polynomial at an amplitude of 0.5 or -0.5 is generated and used for localizing all emitters. The median value of the log-likelihood ratios[7] (LLR) of all localizations is used as a quality metric of the PSF model. The PSF model that gives the highest quality metric will be used as the initial PSF model. The searching procedure takes typically 2-3 minutes.

In general, the second option is recommended over the third if the dominant aberration is known.

##### *4.1.2 Partitioning of data*

Due to overlapping emitters and emitters of low photon counts, many selected emitters are not useful for learning the PSF model. Here we selected emitters by partitioning them into small axial segments, and selecting a small number of emitters within each segment that has the highest log-likelihood ratio from initial localization results. The number of axial segments and the number of emitters in each segment are user defined, here we use 21-31 and 100-200, respectively. After partition, 2000-3000 emitters were selected and used for PSF learning.

##### *4.1.3 Calculation of the forward model*

The forward model for each emitter results in a 2D image and is based on the vectorial PSF model,

$$U(x-x_i, x-y_i, z_i, z_s) = \sum_{\substack{m=x,y \\ n=p_x,p_y,p_z}} \left|\mathcal{F}_{\text{czt}}\left[h(k_x,k_y) w_{mn} e^{i2\pi(k_{z\text{med}}z_i - k_z z_s)} e^{i2\pi(k_x x_i + k_y y_i)}\right]\right|^2 , \quad (4.1)$$

where $w_{mn}$ is the electric field from dipole radiation as defined in Eq. (3.13), $k_{z\text{med}}$ and $k_z$ are defined in Eq. (3.31), and $x_i, y_i, z_i$ define the position of each emitter in the sample medium, in particular, $z_i$ defines the distance of the emitter to the coverslip and is constrained to be positive. $z_s$ is the stage position and defined in section 3.7, $z_s$ is positive when objective is focused on emitters above the coverslip. Since no emitter is below the coverslip during SMLM data acquisition, we constrain $z_s$ to be positive.

In the case of index matching or water dipping objective, $n_{\text{med}} = n_{\text{imm}}$ , changing of $z_s$ and $z_i$ are equivalent, so $z_s$ will be fixed to a user defined value. Independent of index matching, the initial value of $z_s$ should be large enough to ensure $z_i$ to be positive.

The pupil function $h(k_x, k_y)$ can be expressed either as 2D matrices or in terms of Zernike polynomials. For common aberrations from the imaging system, the Zernike-based pupil function is sufficient, while for engineered PSFs where the applied phase variation is not smooth, such as Tetrapod and double-helix PSFs, learning of image-based pupil functions is necessary. Here, user can also choose to set the magnitude part of the pupil function to a unit circle, so that the learning process only learns the phase part of the pupil function. This is a good approximation for most imaging systems and can greatly improve the convergence especially for complex PSF models. Therefore, for most learning results, we fix the magnitude part of the pupil function.

Similar to learning from bead data, the PSF model is blurred with a 2D Gaussian kernel,

$$U_{\text{blur}} = \mathcal{F}_{2D}^{-1}\left[\mathcal{F}_{2D}(U) \cdot e^{-i2(\sigma_x q_x \pi)^2 - i2(\sigma_y q_y \pi)^2}\right] . \quad (4.2)$$

Note that here we use 2D Fourier transform, as for *in situ* PSF learning, $U$ is a 2D matrix instead of a 3D array like in Eq 3.4. We found that the estimated $\sigma_x$ and $\sigma_y$ are slightly smaller to that from the bead data. Although *in situ* PSF learning is expected to extract the PSF model from single fluorophores which are much smaller than fluorescence beads, the extra blurring effect still exists. Finally, including the photon and background of each emitter, the forward model is,

$$U_i = U_{\text{blur}} \cdot s_i + b_i . \quad (4.3)$$

The variables for *in situ* PSF learning are $x_i, y_i, z_i, s_i$ and $b_i$ for each emitter, $\sigma_x$ and $\sigma_y$, $z_s$ and pupil magnitude and phase, $A$ and $\Phi$ , for learning image-based pupil functions, or the Zernike coefficients of the pupil magnitude and phase, $C_{A,p}, C_{\Phi,p}$ , for learning Zernike-based pupil functions.

##### *4.1.4 Calculation of the loss function*

The loss function for *in situ* PSF learning with a Zernike-based pupil function is,

$$loss = a_1 LL + \beta(a_{\text{bmin}} f_{\text{bmin}} + a_{\text{smin}} f_{\text{smin}} + a_{\text{zmin}} f_{\text{zmin}}) , \quad (4.4)$$

where the definition of $LL$, $f_{\text{bmin}}$ and $f_{\text{smin}}$ are the same as the ones for bead data. The last term constrains $z_s$ and $z_i$ to be positive,

$$f_{\text{zmin}} = \min(z_s, 0)^2 + \sum_i \min(z_i, 0)^2. \tag{4.5}$$

The loss function for *in situ* PSF learning with an image-based pupil function is,

$$loss = a_1 LL + a_{\text{sm}} f_{\text{sm}} + \beta(a_{\text{bmin}} f_{\text{bmin}} + a_{\text{smin}} f_{\text{smin}} + a_{\text{zmin}} f_{\text{zmin}}) , \tag{4.6}$$

where $f_{\text{sm}}$ is for smoothing the pupil function and is the same as the one for bead data. However, for learning a pupil function with discontinuous phase variations, $a_{\text{sm}}$ is set to zero.

###### 4.1.5 Iterative learning

Similar to learning PSF from bead data, one iteration of *in situ* PSF learning consists of initial learning, outlier removal and re-learning. However, for *in situ* PSF learning, after one learning iteration, the learned PSF model is biased towards the initial PSF model. Furthermore, when the PSF pattern is complex and the photon counts of the emitters are low, one learning iteration is not sufficient to converge to the correct PSF model. Therefore, for *in situ* PSF learning, multiple learning iterations were used: for the first iteration, the initial PSF model is determined as described in section 4.1.1, for the subsequent iterations, the initial PSF model is the learned PSF model from previous iteration. The minimum photon count, defined by the quantile of all photon counts, used in data partition is set to a user defined value, e.g. 0.4, and decreases by 0.1 for each subsequent iteration until it reaches 0.2. For most tested data, 2 iterations were sufficient, but for the Tetrapod PSF generated from a phase mask, 4-6 iterations were required to converge (SI Fig. 16).

##### 4.2 In situ PSF learning for multi-channel systems

For *in situ* PSF learning of a multi-channel system, a single-channel learning as describe above is performed. However, here in the single channel learning, no data partition is applied, this is to maintain a sufficient number of emitters. After single-channel learning, an initial transformation matrix between each target channel to the reference channel is calculated. As the selected emitters after each single-channel learning might be different between all the channels, a coordinate pairing process is performed again (section 1.3). With the transformation matrices and the learned PSF model from each channel, a multi-channel localization is performed. Based on the localization results, outlier emitters are removed and the remaining emitters are partitioned as described above. The partitioned emitters are used for subsequent multi-channel PSF learning similar to the multi-channel learning from bead data.

##### 4.3 In situ PSF learning of 4Pi-SMLM systems

Learning a 4Pi-PSF model from SMLM data is more complicated than learning the one from bead data. In SMLM data, there is no known sampling of the PSF in z and phase positions. To decouple the phase and z positions of a 4Pi-PSF, the forward model of the 4Pi-PSF is slightly modified.

###### 4.3.1 Single-channel learning of in situ 4Pi-PSF

**Calculation of the forward model**. The 4Pi-PSF is still calculated as the summation of the incoherent and coherent PSF (section 3.5). The coherent PSF is calculated as,

$$U_I(x_i, y_i, z_i, \varphi_i) = \left|\mathcal{F}_{\text{czt}}\left[\left(h_1 e^{i(\varphi_d - \varphi_z) + i\varphi_i} + h_2 e^{i\varphi_z + i\varphi_l}\right) e^{i2\pi(k_x x_i + k_y y_i)}\right]\right|^2, \tag{4.7}$$

where

$$\begin{gathered}\varphi_z = 2\pi[(k_{\text{zmed}} - k_{\text{med}})z_i - k_z z_s], \\ \varphi_d = 2\pi\left(k_{\text{zmed}} - \frac{k_z n_{\text{imm}}}{n_{\text{med}}}\right) z_h.\end{gathered} \tag{4.8}$$

Here, $z_h$ measures the thickness of the sample chamber, $n_{\text{imm}}$ and $n_{\text{med}}$ are the refractive indices of the immersion oil and the sample medium. $\varphi_z$ accounts for index mismatch aberration, however, its calculation is different from the one defined in section 3.7 by a subtraction of $k_{\text{med}} z_i$ , where $k_{\text{med}}, k_z, k_{z\text{med}}$ are defined in Eq. (3.31). The term

$k_{\mathrm{med}} z_i$ only varies with $z_i$ and can be absorbed into the phase variable $\varphi_i$. Therefore $\varphi_z$ becomes a slow varying function of $z_i$ . This modification decouples the variables $z_i$ and $\varphi_i$, any phase drift during the data collection will not affect the learning. In another words, emitters collected at large separation in time can be used together for learning, and the learned PSF model can be used for the entire SMLM dataset. The term $\varphi_d$ accounts for index mismatch aberration in the upper emission path when the upper objective is focused at the bottom coverslip. Because when the emitter is at the bottom coverslip $z_i = z_s$ =0 and $\varphi_z$ =0, the only index mismatch aberration comes from the upper emission path. When the refractive index is matched, $\varphi_d$ equals to zero.

**Calculation of the loss function**. The loss function is

$$loss = a_1 LL + \beta(a_{\mathrm{bmin}} f_{\mathrm{bmin}} + a_{\mathrm{smin}} f_{\mathrm{smin}} + a_{\mathrm{zmin}} f_{z\mathrm{min}} + a_{\alpha} f_{\alpha\mathrm{min}}), \quad (4.10)$$

where

$$f_{\mathrm{zmin}} = \min(z_s, 0)^2 + \min(z_d, 0)^2 + \sum_i \min(z_i, 0)^2. \quad (4.11)$$

The remaining terms are the same as the ones for learning from bead data (section 3.5).

###### *4.3.2 Multi-channel learning of the in situ 4Pi-PSF*

First the 4Pi-PSF model from each channel is generated based on the initial Zernike coefficients. A 4Pi localization is performed using the generated model where the x and y localizations between the four channels are not linked. This is because the transformation matrix is unknown and initially set to an identity matrix. Outlier emitters are removed based on the localization results. A single channel 4Pi-PSF learning is performed on the remaining emitters in each channel where the localization results of the $z_i$, $\varphi_i$, photon and background are used as the initial values. After single-channel learning, an initial transformation matrix between each target channel to the reference channel is calculated. Notice that for *in situ* 4Pi-PSF learning, there is no outlier removal step during the single-channel learning. After all single-channel learnings, the emitters are then partitioned (section 4.1.2) and a multi-channel learning similar to the one for bead data is performed on partitioned emitters.

##### ***4.4 In situ PSF learning of field-dependent PSF***

The *in situ* PSF learning of a single-channel system can be easily extended to learn FD PSFs. The forward model is the same as Eq. (4.1), except that the Zernike coefficients of each emitter are calculated from a linear interpolation of a 2D aberration map (Eq. (3.27)). To ensure emitters are evenly sampled across the FOV, the data partition described in section 4.1.2 is extended to 3D, where the FOV is also divided into $N_x \times N_y$ segments, together with $N_z$ axial segments. The number of segments in each dimension is user defined. The total emitters are partitioned into each 3D segment based on their log-likelihood values as described previously.

The loss function is

$$loss = a_1 LL + a_{\mathrm{sm}} f_{\mathrm{sm}} + \beta(a_{\mathrm{bmin}} f_{\mathrm{bmin}} + a_{\mathrm{smin}} f_{\mathrm{smin}} + a_{\mathrm{zmin}} f_{\mathrm{zmin}}), \quad (4.12)$$

where the smooth constraint $a_{\mathrm{sm}} f_{\mathrm{sm}}$ is defined in Eq .3.29. The remaining process is the same as the one for *in situ* PSF learning of a single-channel system.

#### 5. Calibration of the deformable mirror

Based on the work described by Antonello *et al.*[8], we employed a rigorous methodology to accurately determine the influence function for each actuator in the DM (Boston Micromachines, M140A-35-P01). This ensures that we obtain reliable and precise measurements of the individual actuator's impact on the overall performance of the DM. To begin with, we constructed a Twyman-Green interferometer, a variant of the widely used Michelson interferometer, specifically designed for precise optical testing. This interferometer allowed us to measure the phase information induced by the DM in a reliable manner. To extract the phase information induced by the DM, we employed Fourier-based fringe analysis[9], a powerful technique that enables the precise determination of phase shifts in interferometric measurements. By carefully analyzing the fringes produced by the Twyman-Green interferometer, we were able to accurately quantify the influence of each individual actuator in the DM. In this work, we assumed that the responses of the DM actuators are linear, which is a commonly accepted approximation in similar studies. This assumption allowed us to simplify the analysis and establish a linear model to represent the relationship between the actuator inputs and the resulting output phase $\psi_{DM}$ :

$$\psi_{DM} = \sum \phi_m v_m \tag{5.1}$$

where $\phi_m$ represents the influence function of the $m$-th actuator, $v_m$ is the control voltage applied to the m-th actuator. A vectorized version of Eq. (6.1) is,

$$\Psi_{DM} = \Phi V \tag{5.2}$$

Here, $\Psi_{DM}$ is the discrete measured phase. The control signal V is a vector of size $\mathrm{N}_m$ whereas $\Psi_{DM}$ is a vector of size $\mathrm{N}_k$. The corresponding influence matrix $\Phi$ should be with a size $\mathrm{N}_k \times N_m$, each column of $\Phi$ is a squeezed version of an influence function. By sequentially applying different amplitudes to each actuator and capturing the resulting interference images, we can derive a series of output phases $[\Psi_1, \Psi_2 \dots \Psi_I]$ corresponding to their control signals $[\mathrm{V}_1, \mathrm{V}_2 \dots \mathrm{V}_I]$. Then the influence matrix $\Phi$ can be obtained by solving a simple least-squares problem,

$$\Phi = \underset{\Phi \in \mathbb{R}^{N_k \times N_m}}{\operatorname{argmin}} \sum_I \|\Psi_i - \Phi V_i\|^2 \tag{5.3}$$

Once the influence matrix $\Phi$ is established, we can efficiently determine the control signals required to achieve the desired wavefront shape by performing matrix calculations. This direct wavefront design approach, utilizing the DM influence functions, offers improved accuracy compared to the conventional method of projecting a theoretical Zernike phase onto the DM. To facilitate Zernike-based DM control, it is necessary to make the underlying assumption,

$$\Psi_i \approx \mathbb{Z}\,\xi \tag{5.4}$$

where $\mathbb{Z}$ is a matrix whose columns are Zernike polynomials sampled over the phase measurement grid. $\xi$ is a Zernike coefficients vector with $N_\xi$ components. Since the influence functions of the DM do not resemble Zernike polynomials, more Zernike polynomials ($N_\xi > N_m$) are necessary to describe the influence functions to avoid encountering large approximation errors. Zernike control needs to calculate the mapping between the control signals V and the Zernike coefficients $\xi$, which can be solved using linear regression.

#### 6. CRLB calculation and analytical gradients

The CRLB is calculated using the inverse of the Fisher information matrix $I(\Theta)$, which measures the amount of information that an observation (PSF) carries about the estimated parameters $\Theta$. The Fisher information matrix is defined as[10],

$$I(\Theta)_{ij} = \sum_k \frac{1}{U_k} \frac{\partial U_k}{\partial \Theta_i} \frac{\partial U_k}{\partial \Theta_j} \tag{6.1}$$

where $\Theta$ is a set of parameters being estimated, $k$ denotes pixel index. The parameters $\Theta$ for *in situ* PSF learning is $x_i, y_i, z_i, s_i$ and $b_i$ for each emitter, $\sigma_x$ and $\sigma_y$, $z_s$ and pupil magnitude and phase, $A$ and $\Phi$ , for learning pixel-wised pupil function, or the Zernike coefficients of pupil magnitude and phase, $C(A,p)$,$C(\Phi,p)$ , for learning Zernike expanded pupil function. The $U_k$ is the forward model as defined in Eq. (3.12). The first derivatives of the forward model $U_k$ with respect to the parameters $\Theta$ are given by,

$$\frac{\partial U_k}{\partial \Theta_{x,y,z}} = s_i \sum_{\substack{m=x,y \\ n=p_x,p_y,p_z}} Re\left\{E_{mn}^* \frac{\partial E_{mn}}{\partial \Theta_{x,y,z}}\right\} \tag{6.2}$$

$$\frac{\partial U_k}{\partial C(A/\Phi,p)} = s_i \sum_{\substack{m=x,y \\ n=p_x,p_y,p_z}} Re\left\{E_{mn}^* \frac{\partial E_{mn}}{\partial C(A/\Phi,p)}\right\}$$

$$\frac{\partial U_k}{\partial s_i} = U_{\text{blur}}$$

$$\frac{\partial U_k}{\partial b_i} = 1,$$

$$E_{mn} = \mathcal{F}_{\text{czt}}\left[h(k_x,k_y)w_{mn}e^{i2\pi(k_{\text{zmed}}z_i - k_z z_s)}e^{i2\pi(k_x x_i + k_y y_i)}\right]$$

$$\frac{\partial E_{mn}}{\partial \Theta_{x,y,z}} = \mathcal{F}_{\text{czt}}\left[ik_{x,y,\text{zmed}}h(k_x,k_y)w_{mn}e^{i2\pi(k_{\text{zmed}}z_i - k_z z_s)}e^{i2\pi(k_x x_i + k_y y_i)}\right]$$

$$\frac{\partial E_{mn}}{\partial C(A/\Phi,p)} = \mathcal{F}_{\text{czt}}\left[\frac{\partial h(k_x,k_y)}{\partial C(A/\Phi,p)}w_{mn}e^{i2\pi(k_{\text{zmed}}z_i - k_z z_s)}e^{i2\pi(k_x x_i + k_y y_i)}\right].$$

#### 7. Localization methods

For the localization step during the PSF learning, the data was analyzed with the cubic spline fitting method[1,5]. The cubic spline interpolation of a given PSF model is[4,11]

$$f_{i,j,k}(x,y,z) = \sum_{m=0}^{3}\sum_{n=0}^{3}\sum_{p=0}^{3} a_{i,j,k,m,n,p}\left(\frac{x-x_i}{\Delta x}\right)^m\left(\frac{y-y_i}{\Delta y}\right)^n\left(\frac{z-z_i}{\Delta z}\right)^p \tag{7.1}$$

where $\Delta x$, $\Delta y$ are the x and y pixel sizes, $\Delta z$ is the axial step size of the PSF model, $a_{i,j,k,m,n,p}$ are the spline coefficients and $x_i, y_i, z_i$ are the start position of each voxel $(i,j,k)$. After building the spline PSF model, maximum likelihood estimation (MLE) with Poisson statistics was used to localize beads or single molecules with the objective function given by,

$$\chi_{mle}^2 = 2\left(\sum_k(\mu_k - M_k) - \sum_{k,M_k>0} M_k \ln\left(\frac{\mu_k}{M_k}\right)\right), \tag{7.2}$$

where $\mu_k$ and $M_k$ are the expected photon number and measured photon number in the pixel $k$, respectively. We used a modified Levenberg-Marquardt (L-M) algorithm[11] to minimize $\chi_{mle}^2$ for the parameter estimation.

### Supplementary Tables

Supplementary Table 1. Parameters for single-channel PSF learning

| | Variables = initial | (Scaling factor) |
|---|---|---|
| voxel | $U = 0.002$<br>$x_i, y_i, z_i = 0$<br>$b_i = \text{Eq. } 2.3$<br>$s_i = \text{Eq. } 2.4$ | $(40/w^*)$<br>(1)<br>$(\bar{b})$<br>$(1000w)$ |
| pupil | $A = 1, \Phi = 0$<br>$x_i, y_i, z_i = 0$<br>$b_i = \text{Eq. } 2.3$<br>$s_i = \text{Eq. } 2.4$<br>$\sigma_x, \sigma_y = 0.5$ | $(1200/w)$<br>(1)<br>$(\bar{b})$<br>$(40w)$<br>(1) |
| Zernike | $C_{A,p} = \begin{cases} 1, p = 0 \\ 0, p > 0 \end{cases}$<br>$C_{\Phi,p} = 0$<br>$x_i, y_i, z_i = 0$<br>$b_i = \text{Eq. } 2.3$<br>$s_i = \text{Eq. } 2.4$<br>$\sigma_x, \sigma_y = 0.5$ | $(20/w)$<br>$(20/w)$<br>(1)<br>$(\bar{b})$<br>$(100 \cdot w)$<br>(1) |
| FD | $Z_{\text{map},A,p} = \begin{cases} 1, p = 0 \\ 0, p > 0 \end{cases}$<br>$Z_{\text{map},\Phi,p} = 0$<br>$x_i, y_i, z_i = 0$<br>$b_i = \text{Eq. } 2.3$<br>$s_i = \text{Eq. } 2.4$<br>$\sigma_x, \sigma_y = 0.5$ | $(800/w)$<br>$(800/w)$<br>(1)<br>$(\bar{b})$<br>$(100 \cdot w)$<br>(1) |
| Insitu-pupil | $A = 1, \Phi = 0$<br>$x_i, y_i = 0$<br>$z_{\text{med},i} = MLE$<br>$b_i = \text{Eq. } 2.3$<br>$s_i = \text{Eq. } 2.4$<br>$\sigma_x, \sigma_y = 0.5$<br>$z_s = \text{user defined}$ | $(1200/w)$<br>$(400/w)$<br>$(400/w)$<br>$(\bar{b})$<br>$(40w)$<br>$(4/w)$<br>$(80/w)$ |
| Insitu-Zernike | $C_{A,p} = \begin{cases} 1, p = 0 \\ 0, p > 0 \end{cases}$<br>$C_{\Phi,p} = 0$<br>$x_i, y_i = 0$<br>$z_{\text{med},i} = MLE$<br>$b_i = \text{Eq. } 2.3$<br>$s_i = \text{Eq. } 2.4$<br>$\sigma_x, \sigma_y = 0.5$<br>$z_s = \text{user defined}$ | $(40/w)$<br>$(40/w)$<br>$(6000/w)$<br>$(6000/w)$<br>$(\bar{b})$<br>$(200w)$<br>(1)<br>$(800/w)$ |
| Insitu-FD | $Z_{\text{map},A,p} = \begin{cases} 1, p = 0 \\ 0, p > 0 \end{cases}$<br>$Z_{\text{map},\Phi,p} = 0$<br>$x_i, y_i = 0$<br>$z_{\text{med},i} = MLE$<br>$b_i = \text{Eq. } 2.3$<br>$s_i = \text{Eq. } 2.4$<br>$\sigma_x, \sigma_y = 0.5$<br>$z_s = \text{user defined}$ | $(1600/w)$<br>$(1600/w)$<br>$(6000/w)$<br>$(6000/w)$<br>$(\bar{b})$<br>$(200 \cdot w)$<br>(1)<br>$(800/w)$ |

* $w = \sqrt{\text{median}(s_i)}$

Supplementary Table 2. Parameters for multi-channel PSF learning

| | Variables = initial | (Scaling factor) |
|---|---|---|
| voxel | $U, x_i, y_i, z_i, b_{i,l}, s_i$<br>$T_l = \mathrm{Eq.\,2.22}$ | same as single channel<br>(0.001) |
| pupil | $A_l, \Phi_l, x_i, y_i, z_i, b_{i,l}, s_i, \sigma_{x,l}, \sigma_{y,l}$<br>$T_l = \mathrm{Eq.\,2.22}$ | same as single channel<br>(0.001) |
| Zernike | $C_{A,l,p}, C_{\Phi,l,p}, x_i, y_i, z_i, b_{i,l}, s_i, \sigma_{x,l}, \sigma_{y,l}$<br>$T_l = \mathrm{Eq.\,2.22}$ | same as single channel<br>(0.001) |
| Insitu<br>pupil | $A_l, \Phi_l, x_i, y_i, z_{\mathrm{med},i}, b_{i,l}, s_i, \sigma_{x,l}, \sigma_{y,l}, z_s$<br>$T_l = \mathrm{Eq.\,2.22}$ | same as single channel<br>(0.001) |
| Insitu<br>Zernike | $C_{A,l,p}, C_{\Phi,l,p}, x_i, y_i, z_{\mathrm{med},i}, b_{i,l}, s_i, \sigma_{x,l}, \sigma_{y,l}, z_s$<br>$T_l = \mathrm{Eq.\,2.22}$ | same as single channel<br>(0.001) |

Supplementary Table 3. Parameters for 4Pi-PSF learning

| | Variables = initial | (Scaling factor) |
|---|---|---|
| voxel | $I_l = 0.002$ | $(40/w)$ |
| | $A_{l,\mathrm{Re}} = 0.001/\sqrt{2}$ | $(40/w)$ |
| | $A_{l,\mathrm{Im}} = 0$ | $(40/w)$ |
| | $x_i, y_i, z_i = 0$ | (1) |
| | $\varphi_i = 0$ | (1) |
| | $\varphi_p =$ user defined | (1) |
| | $b_{i,l} =$ Eqs. 2.3 and 2.25 | $(\bar{b})$ |
| | $s_i =$ Eqs. 2.4 and 2.25 | $(100w)$ |
| | $T_l =$ Eq. 2.22 | (0.0001) |
| pupil | $A_{1l}, A_{2l} = 1$ | $(1200/w)$ |
| | $\Phi_{1l}, \Phi_{2l} = 0$ | $(1200/w)$ |
| | $x_i, y_i, z_i = 0$ | (1) |
| | $\varphi_i = 0$ | (1) |
| | $\varphi_p =$ user defined | (1) |
| | $b_{i,l} =$ Eqs. 2.3 and 2.25 | $(\bar{b})$ |
| | $s_i =$ Eqs. 2.4 and 2.25 | $(40w)$ |
| | $\sigma_{x,l}, \sigma_{y,l} = 0.5$ | (1) |
| | $\alpha = 0.8$ | $(40/w)$ |
| | $T_l =$ Eq. 2.22 | (0.0001) |
| Zernike | $C_{\mathrm{A}_1,l,p}, C_{A_2,l,p} = \begin{cases} 1, p = 0 \\ 0, p > 0 \end{cases}$ | $(40/w)$ |
| | $C_{\Phi_1,l,p}, C_{\Phi_2,l,p} = 0$ | $(40/w)$ |
| | $x_i, y_i, z_i = 0$ | (1) |
| | $\varphi_i = 0$ | (1) |
| | $\varphi_p =$ user defined | (1) |
| | $b_{i,l} =$ Eqs. 2.3 and 2.25 | $(\bar{b})$ |
| | $s_i =$ Eqs. 2.4 and 2.25 | $(100w)$ |
| | $\sigma_{x,l}, \sigma_{y,l} = 0.5$ | (1) |
| | $\alpha = 0.8$ | $(40/w)$ |
| | $T_l =$ Eq. 2.22 | (0.0001) |
| Insitu Zernike | $C_{\mathrm{A}_1,l,p}, C_{A_2,l,p} = \begin{cases} 1, p = 0 \\ 0, p > 0 \end{cases}$ | (0.2) |
| | $C_{\Phi_1,l,p}, C_{\Phi_2,l,p} = 0$ | (0.2) |
| | $x_i, y_i = 0$ | (20) |
| | $z_{\mathrm{med},i} = 0$ | (20) |
| | $\varphi_i = 0$ | (1) |
| | $b_{i,l} =$ Eqs. 2.3 and 2.25 | (100) |
| | $s_i =$ Eqs. 2.4 and 2.25 | (1e4) |
| | $\sigma_{x,l}, \sigma_{y,l} = 0.5$ | (1) |
| | $\alpha = 0.8$ | (0.1) |
| | $z_s =$ user defined | (10) |
| | $z_h =$ user defined | (10) |
| | $T_l =$ Eq. 2.22 | (0.001) |
